# Activation of the *Plasmodium* egress effector subtilisin-like protease 1 is achieved by plasmepsin X destruction of the propiece

**DOI:** 10.1101/2023.01.13.524002

**Authors:** Sumit Mukherjee, Armiyaw S. Nasamu, Kelly Rubiano, Daniel E. Goldberg

**Author notes:** Center for Global Infectious Disease Research, Seattle Children’s Research Institute, Seattle, Washington, USA. Address correspondence: Daniel E. Goldberg. **Author Contributions:** S.M., A.S.N. and D.E.G. conceived and designed the study. S.M., A.S.N. and K.C.R. acquired data. S.M. and A.S.N. performed analyzed data. S.M., A.S.N. and D.E.G. wrote the manuscript. These authors contributed equally to this work (Sumit Mukherjee, Armiyaw S. Nasamu).

## Abstract

Following each round of replication, daughter merozoites of the malaria parasite *Plasmodium falciparum* escape (egress) from the infected host red blood cell (RBC) by rupturing the parasitophorous vacuole membrane (PVM) and the RBC membrane (RBCM). A proteolytic cascade orchestrated by the parasite’s serine protease, subtilisin-like protease 1 (SUB1) regulates the membrane breakdown. SUB1 activation involves primary auto-processing of the 82 kDa zymogen to a 54 kDa (p54) intermediate that remains bound to its inhibitory propiece (p31) post cleavage. A second processing step converts p54 to the terminal 47 kDa (p47) form of SUB1. Although the aspartic protease plasmepsin X (PM X) has been implicated in the activation of SUB1, the mechanism remains unknown. Here, we show that upon knockdown of PM X the inhibitory p31/p54 complex of SUB1 accumulates in the parasites. Using recombinant PM X and SUB1, we show that PM X can directly cleave both p31 and p54. We have mapped the cleavage sites on recombinant p31. Furthermore, we demonstrate that the conversion of p54 to p47 can be effected by cleavage at either a SUB1 or PM X cleavage site that are adjacent to one another. Importantly once the p31 is removed, p54 is fully functional inside the parasites suggesting that the conversion to p47 is dispensable for SUB1 activity. Relief of propiece inhibition via a heterologous protease is a novel mechanism for subtilisin activation.

**Significance Statement:** Malaria parasites replicate inside a parasitophorous vacuole within the host red blood cells. Exit of mature progeny from the infected host cells is essential for further dissemination. Parasite exit is a highly regulated, explosive process that involves membrane breakdown. To do this, the parasite utilizes a serine protease, called the subtilisin-like protease 1 or SUB1 that proteolytically activates various effector proteins. SUB1 activity is dependent on an upstream protease, called plasmepsin X (PM X), although the mechanism was unknown. Here we describe the molecular basis for PM X mediated SUB1 activation. PM X proteolytically degrades the inhibitory segment of SUB1, thereby activating it. Involvement of a heterologous protease is a novel mechanism for subtilisin activation.

## Introduction

Malaria remains the most significant parasitic disease throughout the world in terms of morbidity and mortality, with *Plasmodium falciparum* being the deadliest species^1^. During the symptomatic blood stage of infection, the parasites replicate by schizogony within a membrane enclosed parasitophorous vacuole (PV) inside the host red blood cells (RBCs). Following each round of replication, the daughter merozoites escape from the infected RBCs by rupturing the parasitophorous vacuole membrane (PVM) and RBC membrane (RBCM) through an explosive process called egress^2^. As egress allows the released merozoites to invade fresh RBCs, it is critical for the parasite’s dissemination.

Parasite egress follows an ‘inside-out’ route where the PVM disruption precedes the RBCM rupture^3,4^. Upon schizont maturation, and approximately 10 minutes prior to the actual release of the merozoites the PVM first rounds up due to an intracellular calcium oscillation^2^. Calcium also activates the cyclin-dependent protein kinase 5 (CDPK5)^5^. Within the next few minutes, the parasite’s cGMP-dependent protein kinase G (PKG) is activated. In coordination with CDPK5, the activated PKG triggers discharge of the subtilisin-like serine protease 1 (SUB1) from the exonemes into the PV^6–8^. PKG-mediated SUB1 discharge is a prerequisite for PVM rupture^9^. Once inside the PV, SUB1 proteolytically activates several merozoite surface proteins and soluble proteases including the MSPs and SERA6 respectively^9,10^. Coordinated action of these effectors leads to the disassembly of host cell cytoskeleton, perforation, and finally rupture of the RBCM.

SUB1 is synthesized as a precursor of 82 kDa (p82) with a C-terminal catalytic domain and an N-terminal extension called the prodomain (PD)^11^. Following translocation into the endoplasmic reticulum (ER), p82 undergoes autocatalytic cleavage to yield a C-terminal 54 kDa intermediate (p54) that remains non-covalently bound to the cleaved PD, also called the propiece (p31)^11^. Post ER, but before discharge, p54 is further processed to a 47 kDa form (p47). In parasites this second cleavage of SUB1 is dependent on the exoneme-localized aspartic protease plasmepsin X (PM X)^12,13^. Importantly, transgenic parasites lacking PM X or parasites treated with PM X inhibitors, fail to process SUB1 substrates, leading to an egress block^12–15^. This suggests that PM X is involved in the activation of SUB1, although the mechanism remains unknown.

Before SUB1 can be activated, the PD needs to be removed. For most subtilisins, the PD is required for folding and is inhibitory until removal^16^. In the apicomplexan *Toxoplasma gondii*, the microneme protein TgMIC5 acts as a PD for TgSUB1 and regulates the enzymatic activity of the latter^17,18^. TgMIC5 shares both structural and sequence similarities with the PfSUB1 PD, and the PDs from other *Plasmodium* species^17^. This hints at a similar regulatory role for the PfSUB1 PD in the activity of the protease. In fact, a recombinantly made p31 was shown to inhibit the activity of both the recombinant and native SUB1^19^. Consistent with its inhibitory function, crystal structures of SUB1 revealed tight interaction between p31 and the catalytic domain, occluding the active _site_^20,21^.

In this study we sought to understand the following mechanistic questions: how does the knockdown of PM X lead to the accumulation of the inactive form of SUB1? Does conversion of p54 to p47 activate SUB1? We report that PM X removes the inhibitory p31 from the p31/p54 complex by directly cleaving p31 at multiple sites. We further showed that once p31 is removed, the p54 form is fully functional inside the parasites. Involvement of an upstream protease in the removal of the propiece is a novel mechanism for subtilisin activation.

## Results

### PM X mediates cleavage of p31 in the p31/p54 complex of SUB1

Existing antibodies against SUB1 can detect the p54 and p47 species but not p31^6^. Therefore, to visualize p31 in cultured parasites, we tagged the N-terminus of SUB1 with triple hemagglutinin (3x HA). To accomplish this, we replaced the endogenous locus with a recodonized version of *Sub1* by CRISPR/Cas9 mediated genome editing. In the recodonized *Sub1*, the 3x HA was inserted between the predicted signal peptide sequence and the start of p31, thus labelling p31 at its N-terminal end (Fig. 1A). We edited the *Sub1* locus in the anhydrotetracycline (aTc)-regulated PM X^apt^ transgenic parasite line^12^ to track the fate of p31 in the presence or absence of PM X. As shown in Fig. 1B (left panel, top), we were able to pull down p31 from parasite lysates in the absence of the PM X. Additionally, we observed that the p54 coprecipitated with p31 nearly completely under PM X knockdown condition (Fig. 1B, left panel bottom). This validated the physical association between p54 and the p31, as reported previously^11^. In contrast, p47 was the major species of SUB1 in parasites expressing PM X and was overwhelmingly detected in the flow-through fraction (Fig.1, right panel, top). Together, these data suggest that PM X mediates the degradation of p31, thereby destabilizing the p31/p54 complex of SUB1.

**Figure 1.**
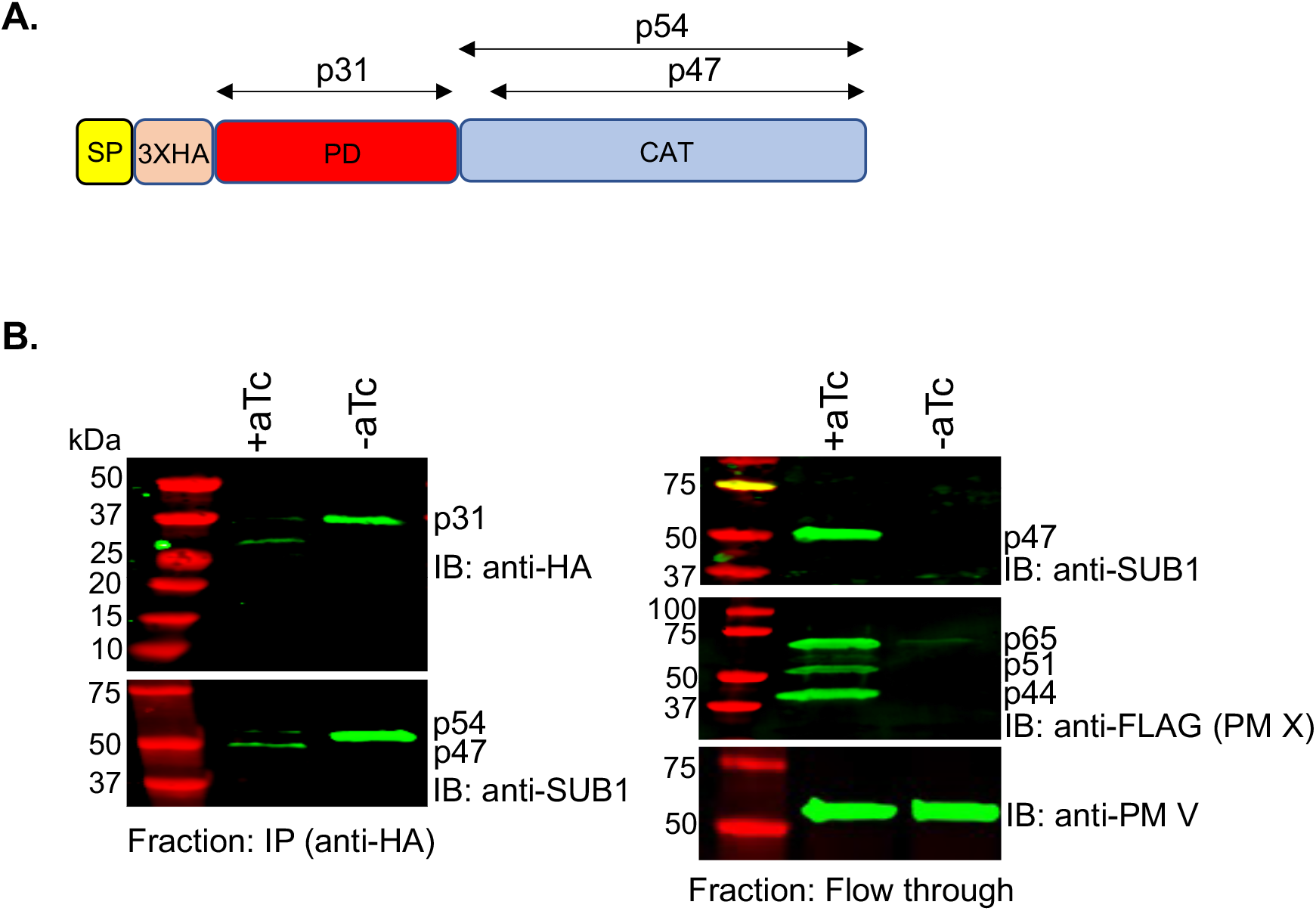
PM X mediates cleavage of SUB1 p31 *in vivo*. **(A)** Schematic of the modified endogenous SUB1 in the aTc-regulatable PM X^apt^ strain of *P. falciparum*. The triple hemagglutinin (3x HA) tag was placed between the signal peptide (SP) and the prodomain, thereby enabling us to detect p31 by western blot. CAT: catalytic domain. **(B)** Western blots. MACS-synchronized PM X^apt^ parasites with edited SUB1 were grown in the presence or absence of aTc for 45 h. Cell lysates prepared from parasite samples were subjected to immunoprecipitation with the anti-HA antibodies. The immunoprecipitated fractions (IP) were fractionated by SDS-PAGE and blotted with the indicated antibodies (left panel). Western blots were also performed with the flow-through (FT) fractions to detect the C-terminal domain of SUB1 (right panel, top), PM X (right panel, middle), and PM V (loading control, right panel, bottom). Experiments were repeated three times and shown is a representative blot.

### Recombinant PM X (rPM X) cleaves recombinant p31 (rp31) in vitro at multiple alternate sites

To further understand the action of PM X on p31 of SUB1, we carried out *in vitro* cleavage assay using recombinantly synthesized proteins. rPM X was produced from mammalian cells as reported earlier^13^. For rp31, the p31 sequence was expressed and purified from *E. coli*. This rp31 was tagged at the N and C-terminal ends with 3x HA and 6 histidines (6x His) respectively. As shown by Coomassie staining of an SDS-PAGE, and by western blotting with an anti-HA antibody (Fig. 2A, WT), the full-length form of purified rp31 ran close to the 37 kDa molecular weight marker. Additionally, a minor translation product of about 20 kDa was seen only by Coomassie staining (Fig. 2A, black arrow). N-terminal sequencing of this band by Edman degradation confirmed that it was translated from an internal start codon (Fig. 2B, met^63^, black underlined, Additional Supplementary file 1). As revealed by both Coomassie and western blot, the 37 kDa band disappeared when rp31 was treated with rPM X (Fig. 2A, WT). In addition, we observed two extra bands of approximately 25 kDa and 10 kDa by Coomassie blot but not by western blot with the anti-HA antibody (Fig. 2A, blue and green arrows respectively), indicating that both bands were C-terminal fragments. Finally, addition of the PM X inhibitor CW-117^12^ into the reaction mixture blocked the processing of rp31 to these shorter fragments. Together these data suggest that the rPM X is able to cleave rp31 at two sites, of which one is close to the N-terminus.

**Figure 2.**
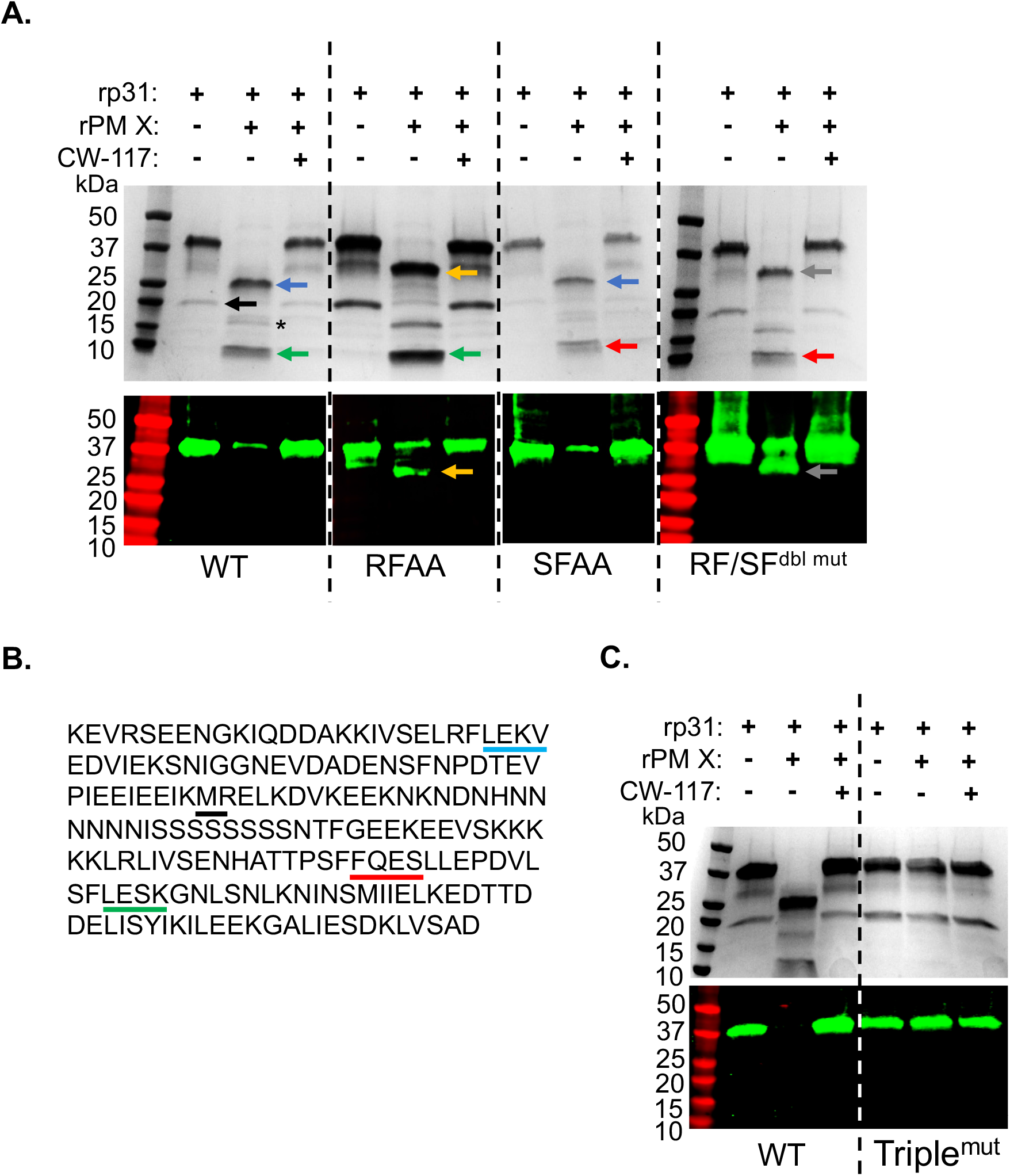
PM X cleaves p31 *in vitro* at alternate sites. **(A)** A Coomassie blot (top) and anti-HA antibody western blot (bottom) showing the cleavage of recombinant p31 (rp31), either wild type (WT) or the indicated mutants, by recombinant PM X (rPM X) following a 3 h assay in the PM X activity buffer. As shown, addition of the PM X inhibitor CW-117 inhibited PM X-mediated cleavage of p31. Blue arrows: cleavage product with an N-terminal tetrapeptide LEKV, green arrows: cleavage product with an N-terminal tetrapeptide LESK, orange arrows: the N-terminal cleavage product in the RFAA mutant, grey arrow: the N-terminal cleavage product in the RFAA/ SFAA double mutant (RF/SF^dbl mut^), red arrows: cleavage product with an N-terminal tetrapeptide FQES, black arrow and star: secondary polypeptide translated from an internal methionine and its cleavage product, respectively. **(B)** Sequence of SUB1 p31 with the N-terminal tetrapeptide sequences for the cleaved p31 fragments and the internal start codon for secondary translated polypeptide in **A** underlined. Colors correspond to the arrows highlighted by same colors in **A. (C)** Coomassie blot (top) and western blot with anti-HA antibody (bottom) showing that a triple cleavage mutant p31 (Triple^mut^) is resistant to cleavage by rPM X. Experiments were repeated three times and shown are representative blots.

To precisely determine the PM X cleavage sites on rp31, we excised the 25 kDa and 10 kDa bands from SDS-PAGE and subjected them to N-terminal sequencing. For both fragments unique profiles of residues were observed during the first four cycles of Edman degradation. The N-terminal tetrapeptide sequence for the 25 kDa fragment was LEKV (Fig. 2B, blue underlined, Additional Supplementary file 2) while that for the 10 kDa fragment was LESK (Fig. 2B, green underlined, Additional Supplementary file 3). Of note, the amino acid sequences that span the N-terminal ends of these fragments (Fig. 2B, ^48^RFLE^51^ and ^164^SFLE^167^) conform with the known PM X substrate cleavage specificity^15^. To test this further, we expressed two different mutant rp31 versions in *E. coli* by mutating the last two residues within the ^48^RFLE^51^ or ^164^SFLE^167^ motifs to alanine (^48^RFLE to ^48^RFAA, or ^164^SFLE to ^164^SFAA). Such di-mutation was shown to abolish PM X cleavage of a fluorogenic peptide or self-processing^13–15^. When treated with rPM X, the RFAA mutant was processed to generate two fragments, one running as 10 kDa similar to that in the WT, and the other running as approximately 27 kDa (Fig. 2A, green and orange arrows respectively). Importantly, the 27 kDa fragment retained the N-terminal 3x HA tag and was detected by the western blot with anti-HA antibody. Interestingly, when treated with rPM X, the processed fragments from ^164^SFAA mutant rp31 ran similarly to those from WT rp31. N-terminal sequencing of the shorter fragment (Fig. 2A, red arrow), however, revealed a different tetrapeptide (FQES in Fig.2B, red underlined, Additional Supplementary file 4), that is just 9 amino acids upstream of the ^164^SFLE^167^ sequence. Mutating both ^48^RFLE^51^ and ^164^SFLE^167^ sequences together (^48^RF/^164^SF^dbl mut^), on the other hand, generated a PM X cleavage pattern with an intact N-terminal fragment of about 27 kDa (Fig. 2A, grey arrow), and a C-terminal 10 kDa (Fig. 2A, single star) fragment with an N-terminal tetrapeptide FQES. This suggested that rPM X could cleave both the ^164^SFAA and the ^48^RF/^164^SF^dbl mut^ at the same site, generating identical C-terminal fragments. Consistent with this, the amino acids flanking the N-terminus of this fragment (^151^SFFQ^154^, Fig. 2B) also conforms with the PM X cleavage specificity. To check this further, we made a triple mutant rp31 by mutating the ^48^RFLE^51^, ^151^SFFQ^154^ and ^164^SFLE^167^ sites together (Triple^mut^). As shown by both Coomassie and western blots, Triple^mut^ rp31 was completely resistant to rPM X-mediated cleavage (Fig. 2C). Together the data suggest that the rPM X can directly cleave rp31 at multiple sites, of which the C-terminal sites are alternatively targeted by rPM X.

We were also able to purify a recombinant form of SUB1 (rSUB1) from mammalian cells as a secreted protein (Supplementary Fig. 1). This rSUB1 was tagged at the N and C-termini with 3x HA and 6x his respectively. As reported earlier^11^, in the presence of tunicamycin added to block N-glycosylation, the purified rSUB1 was a complex of the processed p31/p54. When the p31/p54 complex was incubated in the PM X activity buffer, only in presence of rPM X, we observed cleavage of p31 by Coomassie and western blots (Fig. 3). This suggested that the rPM X can access the target sites on p31 in the correctly folded rSUB1. Additionally, in the rPM X-treated sample we observed that the C-terminal fragment ran faster than that in the control sample (no PM X), indicating rPM X-mediated processing of the p54 fragment occured *in vitro*.

**Figure 3.**
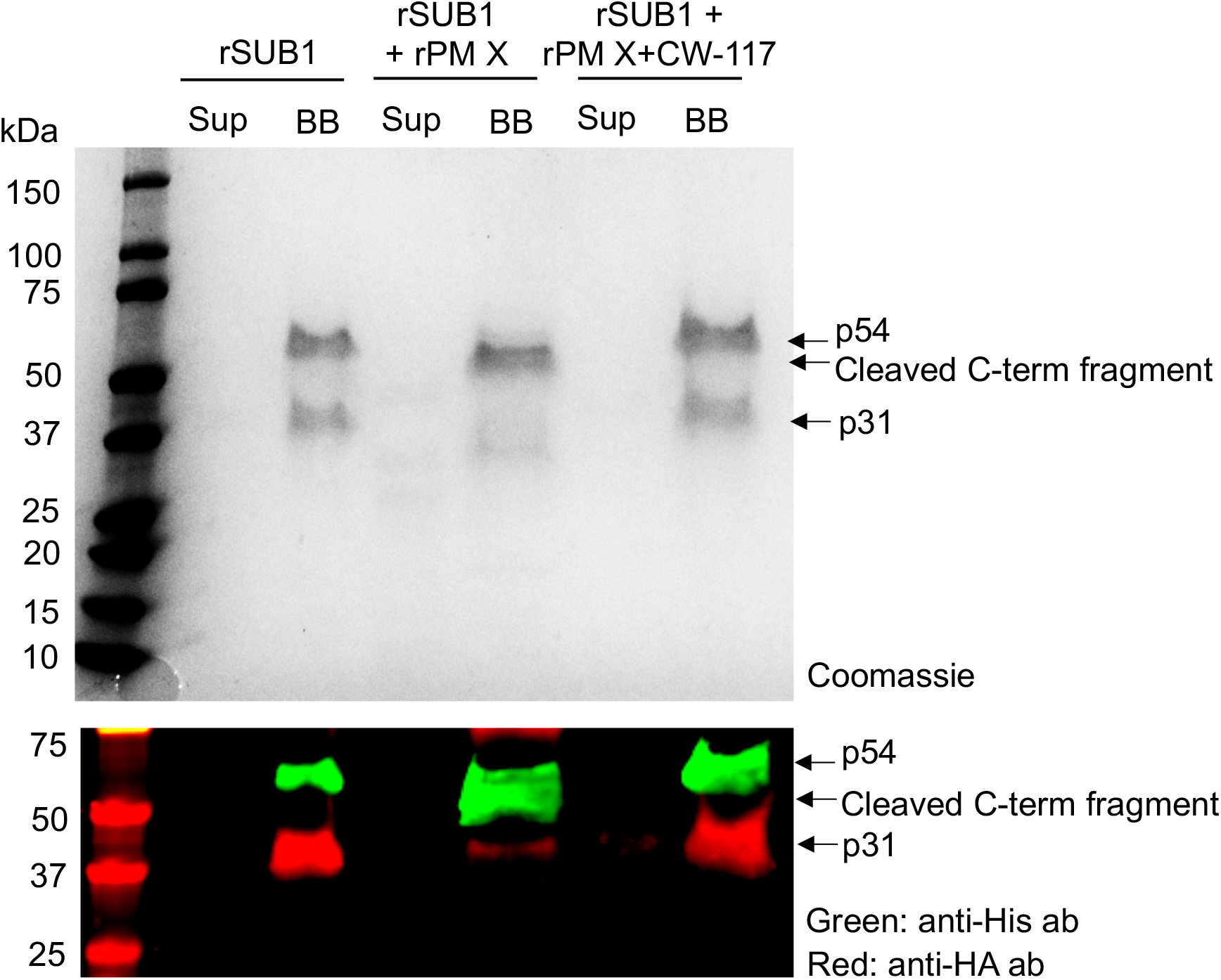
PM X cleaves both p31 and p54 of SUB1 *in vitro*. Coomassie blot (top) and western blot with indicated antibodies (bottom) showing the cleavage of p31 and the C-terminal p54 fragment from the p31/p54 complex of a mammalian expressed recombinant SUB1 (rSUB1). The rSUB1 was tagged at the N and C-terminal ends with 3x HA and 6x His respectively. The purified p31/p54 complex was immobilized on His trap magnetic beads and a PM X cleavage assay was performed on the beads with rPM X in the PM X activity buffer. 3 h post incubation, the supernatants (Sup) were separated by placing the reaction tubes on a magnet. The C-terminal fragments of rSUB1 that remained bead bound (BB) were eluted with buffer containing 500 mM imidazole. Both the Sup and BB fractions were boiled in SDS-PAGE buffer and analyzed. Experiment was repeated three times and shown are representative blots.

### p54 to p47 conversion involves both direct cleavage by PM X and SUB1 autocatalysis but is dispensable for parasite growth

An earlier study reported that the p54 to p47 conversion for rSUB1 occurs through autocatalytic cleavage after the aspartate within the ^242^SMLEVENDAE^251^ motif^11^. However, our *in vitro* cleavage assay as depicted in fig. 3 suggested that PM X could cleave both p31 and the p54 of SUB1. Consistent with this, previously a recombinant form of PM X has been shown to cleave the ^242^SMLEVENDAE^251^ peptide after the methionine^14^. Therefore, to elucidate the processing of the p54 to p47 further, we moved to cultured parasites and expressed either WT or cleavage site mutant forms of SUB1 as C-terminally V5-tagged second copies (Fig. 4A). For the autocatalytic cleavage mutant, we mutated the V^246^ to K and the D^249^ to L together in one construct (47^VK/DL^). This di-mutation was previously shown to abolish cleavage of a peptide substrate by rSUB1^22^. As a PM X cleavage mutant we mutated the LE to AA in the ^242^SMLE^245^ sequence (^242^SMAA) ^14^. Finally, in one construct we mutated both the p47 autocleavage and the proposed PM X cleavage sites together (Dbl^mut^). Besides looking at the processing of the different cleavage mutants, we wanted to determine if they were functional or not. For this, we expressed the second copy mutants in an endogenous SUB1 inducible knockout line, SUB1^loxP^. In this line, the parasites stably express the rapamycin (RAP)-inducible dimerizable Cre recombinase (DiCre)^23^. To excise the segment of the SUB1 gene encoding the catalytic residues upon RAP treatment, we modified the *sub1* locus with loxP sequences and simultaneously added 3x HA at the C-terminal end as described earlier^9,24^ (Supplementary Fig. 2A). PCR (Supplementary Fig. 2B, left) and Western blot (Supplementary Fig. 2B, right) demonstrated rapid and efficient RAP-induced excision of the floxed DNA sequences and ablation of SUB1 expression. As shown earlier^9^, addition of RAP to synchronized ring-stage cultures of the SUB1^loxP^ parasites led to a severe growth defect that was detectable from the second cycle on.

**Figure 4.**
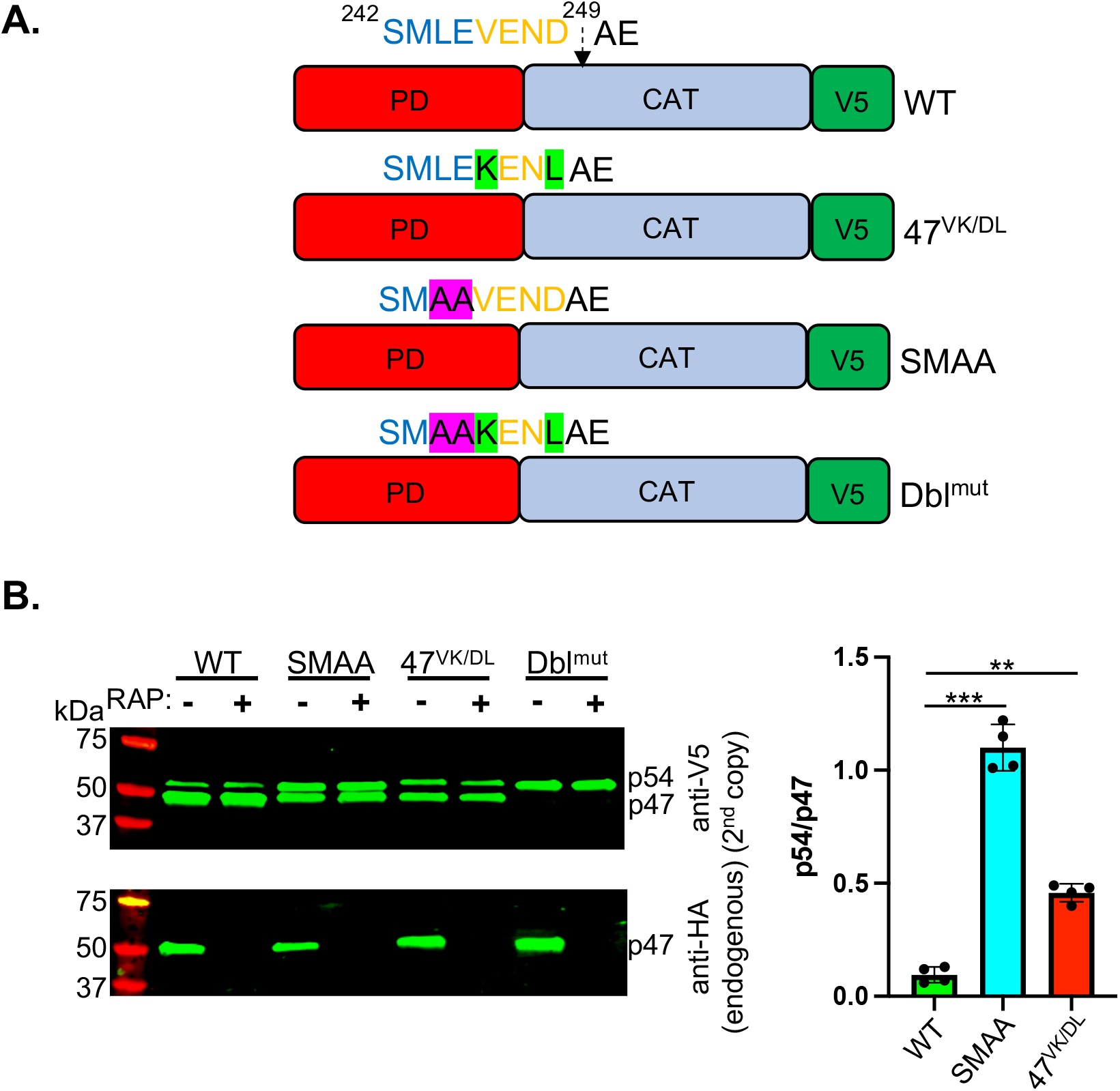
Conversion of p54 to p47 can be effected by SUB1 autocatalysis and PM X-mediated cleavage. **(A)** Schematics of the wild type (WT) or the different cleavage mutant SUB1 constructs that were tagged at the C-terminus with V5 epitope and expressed as second copies in the SUB1^loxP^ transgenic line. **(B)** Left: western blot showing the processing of the WT or the different cleavage mutants SUB1 expressed as second copies (top panel, V5) in presence (-RAP) or absence (+RAP) of the endogenous SUB1 (bottom panel, HA). Samples were harvested from synchronized 44-47 h old schizonts that were treated with 10 nM rapamycin (RAP) in the ring stage for 3 h. Right: Quantification from three independent western blots as on the left. Error bars represent standard deviation. Data were analyzed statistically by two-tailed Student’s *t*-test. **: *p* < 0.01, ***: *p* < 0.001.

To understand cleavage of the second copy SUB1 we analyzed the protein expression by western blot in synchronized schizonts (44-47 h old) that were either pretreated with RAP or not. Aliquots of the same samples were cultured further and tested for rescue of the growth defect due to depletion of endogenous SUB1. Under conditions when the endogenous SUB1 was processed to p47, the WT second copy was detected primarily as the p47 as well (Fig. 4B). However, both the 47^VK/DL^ and the ^242^SMAA mutants demonstrated inefficient conversion of p54 to p47. Moreover, the degree of the defect was more pronounced for the ^242^SMAA mutant than the 47^VK/DL^. Nevertheless, the fact that both were able to partially process to p47 suggested that the two sites could be alternatively used. Consistent with this, the Dbl^mut^ was detected only as the p54, suggesting a complete blockage in the p54 to p47 conversion. Importantly, each of the cleavage site mutants including the Dbl^mut^ was able to fully rescue growth in absence of endogenous SUB1 (Fig. 5). This suggested that the p54 to p47 conversion is dispensable for SUB1 functions in the parasites.

**Figure 5:**
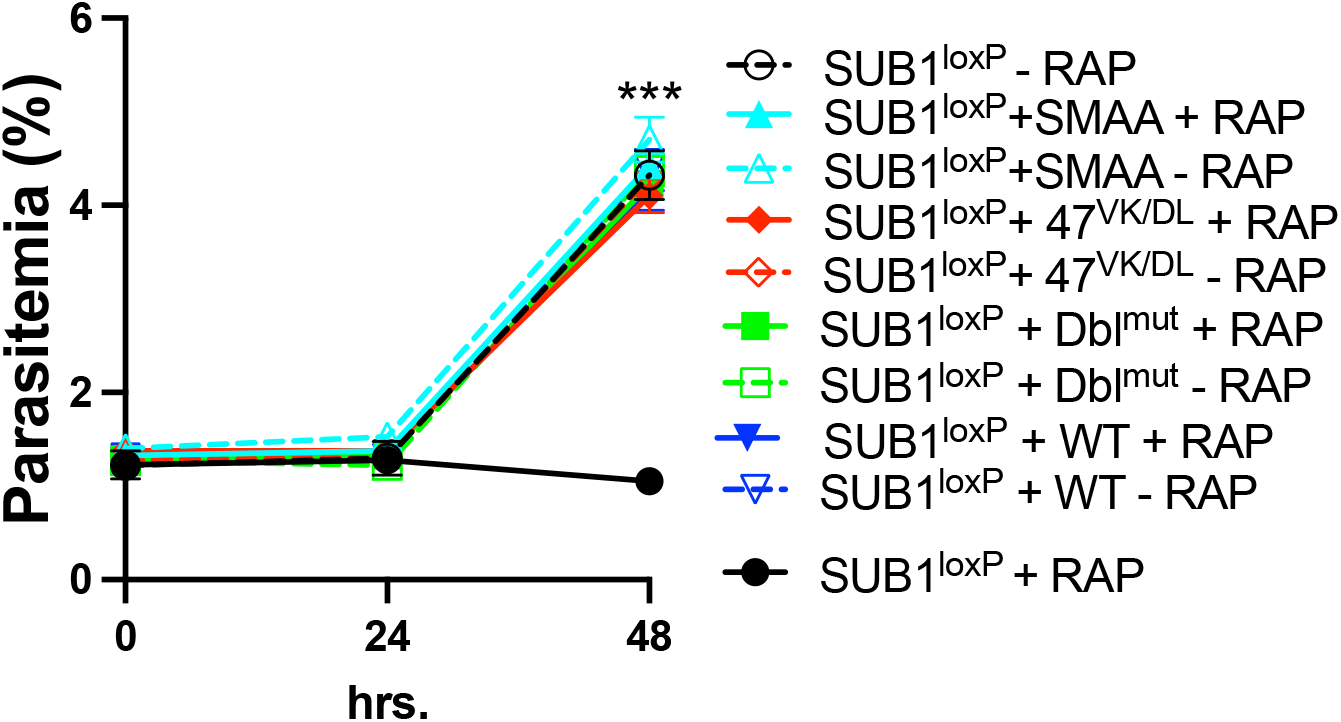
Conversion of p54 to p47 is dispensable for parasite growth. Synchronized parasites were cultured as in figure 4B. However, the initial parasitemia was set to 1%. Mean values from three independent experiments are shown and error bars represent standard deviations. Statistical two-tailed Student’s *t*-test was done to determine the significance of difference in growth between the parasites lacking endogenous SUB1 (SUB1^loxP^ + RAP) and parasites expressing only double cleavage mutants as in figure 4 (Dbl^mut^ + RAP). ***: *p* < 0.001.

### Primary processing of SUB1 is strictly autocatalytic and is required for intracellular trafficking of SUB1

Analysis of the p47 cleavage mutant forms of SUB1, as described in the previous section suggested two alternative pathways for p54 to p47 conversion: one PM X mediated, and another by autocatalysis. To test the same for the primary processing of SUB1 from p82 to p54, we carryout out similar experiments as in the previous section, but this time we mutated the V^214^ to K and the D^217^ to L within the proposed p54 autocleavage site (^213^LVSAD ↓ NID^220^, arrow indicates the scissile bond, Fig. 6A, 54^VK/DL^)^11^. While both the endogenous SUB1 and the WT second copy were processed to p47, the 54^VK/DL^ mutant accumulated in the precursor p82 form (Fig. 6B). Additionally, immunofluorescence microscopy (IFA) with mature schizonts showed that the WT second copy SUB1 but not the 54^VK/DL^ mutant was colocalized with the endogenous SUB1 in the exonemes (Fig. 6C, top and middle). The 54^VK/DL^ mutant, on the other hand, showed near complete colocalization with the ER marker BiP (Fig. 6C, bottom). Finally, consistent with the inhibitory function of p31, parasite growth assay revealed that the 54^VK/DL^ mutant failed to rescue the growth defect due to the depletion of the endogenous SUB1 (Supplementary Fig. 3). Together, these data suggested that unlike the secondary processing, the primary conversion of SUB1 precursor to p54 is strictly autocatalytic and is required for SUB1’s intracellular trafficking.

**Figure 6:**
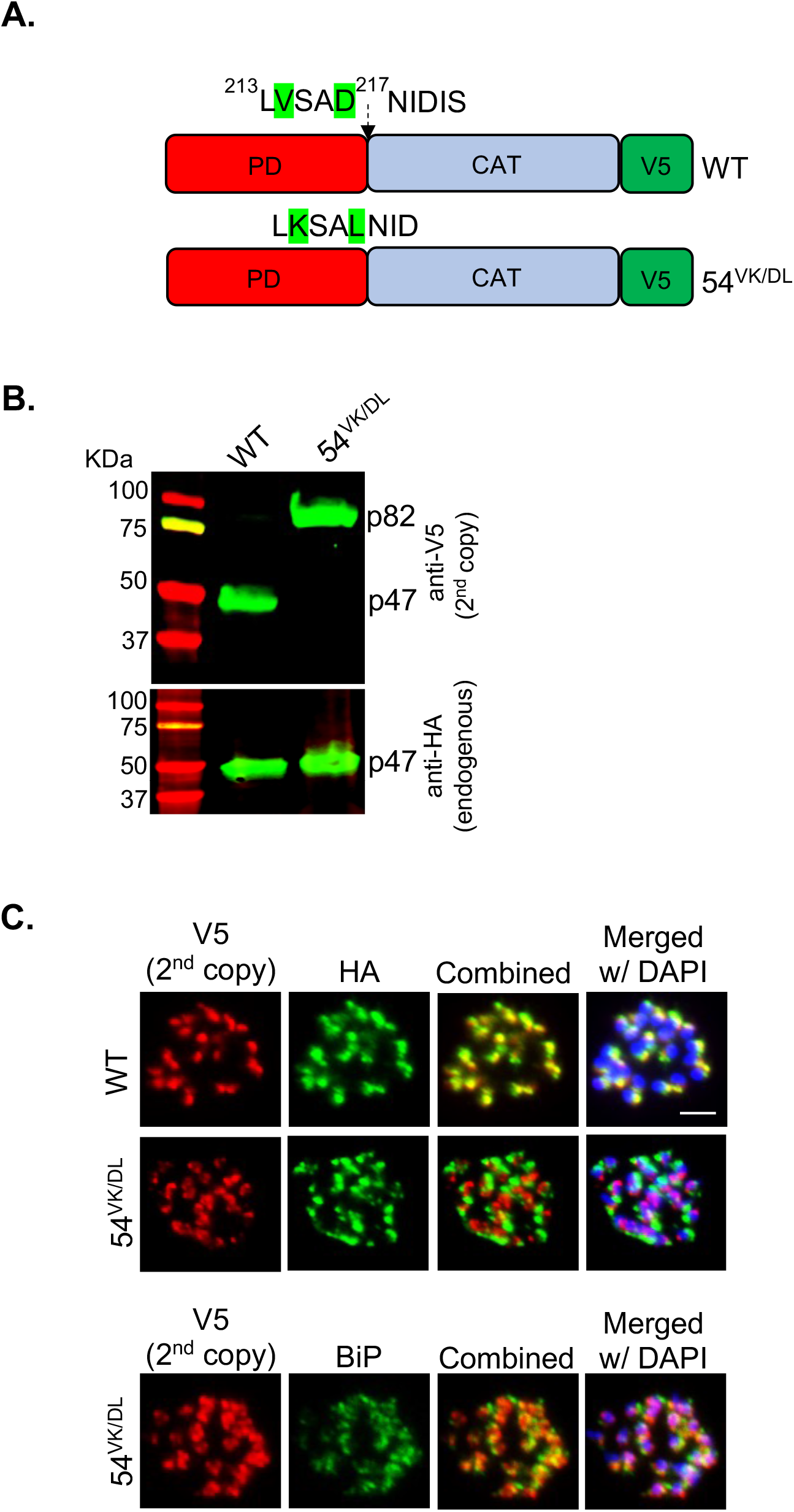
Primary processing of the SUB1precursor to p54 is autocatalytic and is required for intracellular trafficking. **(A)** Schematics of the wild-type (WT) or the p54 cleavage mutant (54^VK/DL^) constructs that were tagged at the C-terminus with V5 epitope and expressed as second copy in the SUB1^loxP^ transgenic line. **(B)** Western blot showing the processing of the WT or the 54^VK/DL^ mutant SUB1 expressed as second copies (top panel, V5) and the endogenous SUB1 (bottom panel, HA). Samples were harvested from synchronized 44-47 h old schizonts. Experiment was repeated three times. Shown is a representative blot. **(C)** Representative images from IFAs showing that the WT second copy (V5) but not the 54^VK/DL^ mutant (V5) was colocalized with the endogenous SUB1 (HA). The 54^VK/DL^ mutant, however, was colocalized with the ER marker BiP. Scale bar: 2 μm. Experiments were repeated two times.

## Discussion

Egress of malaria parasites from infected host RBCs is an essential process for propagation. The serine protease SUB1 plays a central role in this process by proteolytically activating several downstream effectors that mediate the breakdown of the PVM and the RBCM. SUB1, like other subtilisins is synthesized as a proenzyme (p82) with an N-terminal prodomain (PD). The PD facilitates the folding of the catalytic domain of SUB1 while acting as a tightly bound inhibitor for the cognate enzyme^19^. The existing model for proenzyme processing based on a recombinant SUB1 suggests that both the primary processing of p82 to the p31/p54 intermediate complex and the subsequent conversion into p47 are dependent on the autocatalytic activity of SUB1^11^. We and others have shown that the conversion of p54 to p47 in parasites is dependent on the upstream protease PM X^12–14^. Importantly, depletion of PM X in parasites renders SUB1 nonfunctional, suggesting a crucial role for PM X in the activation of the SUB1. In this current study we investigated the mechanism by which PM X regulates SUB1 activity. We showed that both p31 and p54 are direct substrates of PM X (Fig. 2 and 3). Further we showed that PM X-mediated proteolytic degradation of p31 in the p31/ p54 complex is the crucial step for SUB1 activation in *P. falciparum* (Fig. 5).

Using the *E. coli* expressed p31 we showed that PM X can cleave p31 at three sites, one towards the N terminus (^48^RFLE^51^), and the other two close to the C-terminus (^151^SFFQ^154^ and ^164^SFLE^167^) (Fig. 2). Of the two C-terminal sites, cleavage at ^164^SFLE^167^ was detected in WT p31. However, cleavage at the ^151^SFFQ^154^ site was detected only when the ^164^SFLE^167^ sequence was mutated. One possibility is that ^164^SFLE^167^ is the preferred sequence for PM X in WT p31. Alternatively, both the ^164^SFLE^167^ and the ^151^SFFQ^154^ sequences could be targeted by PM X on the same protein. These two cleavage sequences are only 13 amino acids apart from each other. Therefore, it is possible that the small polypeptide generated due to cleavage at these two sites ran off the SDS-PAGE, preventing us from detecting it. Importantly, all three cleavage sequences on p31 are conserved in SUB1 from multiple species of *Plasmodium*^21^. The ^48^RFLE^51^ sequence is part of an alpha helix that has been shown to fold over and interact with the active site in the crystal structure of PvSUB1^21^. The C-terminal cleavage sequences (^151^SFFQ^154^ and ^164^SFLE^167^), on the other hand belong to a segment of p31 which folds into a structure similar to the pro-region from bacterial subtilisins, and mediates the inhibition of the protease activity^20,21^. One consequence of p31 cleavage, therefore, could be the disruption of the interaction between p31 and p54. Secondly, it could lead to a change in the conformation of the inhibitory residues of p31 lodged in the active site. In fact, truncation at the C-terminal end of p31 has been shown to substantially reduce the inhibition of SUB1 activity *in vitro*^19^. Whether inside the parasites PM X targets the *in vitro*-identified cleavage sequences on p31 remains unknown. Mutations at these sequences in the parasites led to the accumulation of the mutant SUB1 in the precursor form (p82) that remained sequestered in the parasite ER, possibly due to the misfolding of the protease (data not shown). Nevertheless, the accumulation of full-length p31 in parasites only in the absence of PM X, together with the findings from the *in vitro* assays evoke a model in which SUB1 maturation involves direct cleavage of p31 by PM X to destabilize the inhibitory p31/p54 complex (Fig. 7).

**Figure 7:**
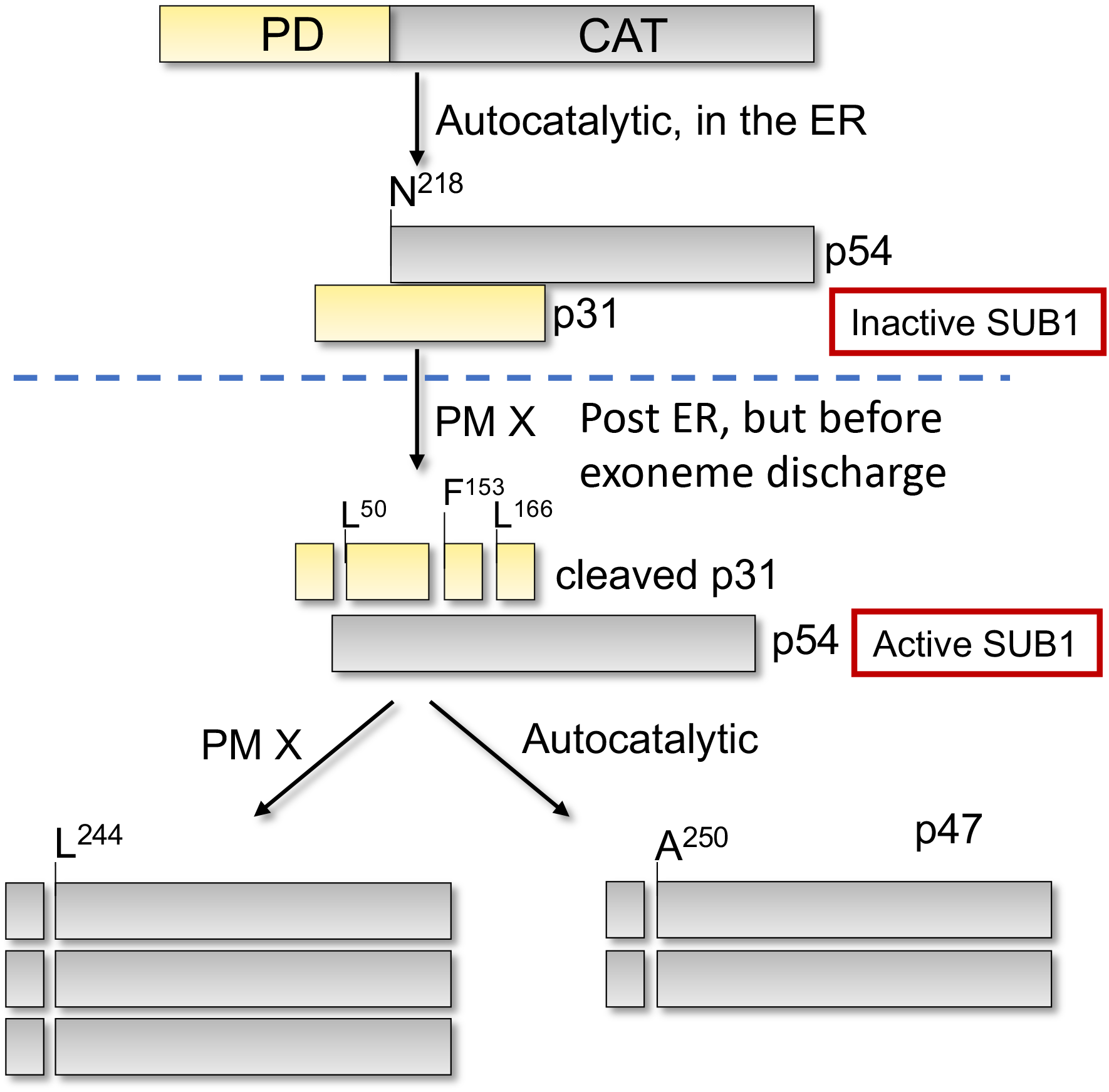
Model for the activation of SUB1 in *Plasmodium falciparum* as suggested by the present data.

Another important finding that comes from our study is that the conversion of the p54 to p47 is dispensable for parasite growth. Previously, p47 was assumed to be the active form of SUB1. Nevertheless, since the recombinantly made p54 was purified as a complex with p31, it was not possible to determine the activity of the p54. Moreover, our attempt to purify p54 without a PD also failed. Therefore, we took the advantage of the previously published data to mutate the potential p47 autocleavage site and determine the fate of the mutant SUB1 directly in the parasites^11^. This revealed that the conversion of p54 to the terminal p47 form can be effected by either a direct cleavage by PM X or by a less efficient autocatalysis. Currently, the cellular compartment where the second processing of SUB1 takes place remains unknown. However, in theory this must take place in a space that maintains enough acidity for PM X to be functional. Although the effect of pH in controlling the SUB1 activity has not been tested, a redox-sensitive, solvent exposed disulphide bond has been shown to be critical for SUB1 activity^20^. An acidic environment (potentially in the exoneme) could, therefore, maintain the processed SUB1 in an inactive state.

Unlike the p54 to p47 conversion, the primary processing of SUB1 in the parasite is strictly autocatalytic in nature (Fig. 6B). Importantly, a mutant SUB1 that failed to cleave itself at the PD-p54 junction was retained in the parasite ER. This suggested that like other subtilisins^25^, the primary processing of SUB1 is critical for intracellular trafficking of the protease. Whether trafficking through the Golgi requires distinct SUB1 folding intermediates remains to be tested.

Subtilisins in general are synthesized as inactive proenzymes that are activated upon reaching the site of action^25–29^. Typically, the PD facilitates folding of the rest of the proenzyme. Besides acting as a chaperone, the PD also acts as an endogenous inhibitor of the protease activity. When activation is required, there is a release and/ or autocatalytic degradation of the PD that can happen both in *cis* or in *trans*. For most subtilisins, the PD displacement has been proposed to be the rate-limiting step and occurs due to pH changes during translocation to the site of activity. Some of the PDs have been proposed to have pH sensors that trigger the process^30^. PfSUB1 appears to use a novel strategy for activation. It requires an endogenous activating protease, PM X that by destroying the propiece, relieves inhibition. Why the malaria parasite does it this way is unclear. Perhaps the environment of the exoneme is not conducive to the conformational change needed for automaturation.

## Materials and Methods

### Reagents

All primers were obtained from Integrated DNA Technologies. List of primers used in this study can be found in Supplementary table 1. Restriction enzymes were purchased from New England Biolabs. His trap magnetic sepharose beads were from Cytvia. The Expi293 cells, transfection reagent ExpiFectamine, Blasticidin S HCl, Ni-NTA beads, the mouse anti-HA antibody coated magnetic beads, and the rabbit anti-V5 antibody were from Thermo Fisher. The rat anti-FLAG antibody was from Novus Biologicals, the rabbit anti-HA antibody, the rabbit anti-His antibody, and rapamycin were from Sigma. The mouse anti-PM V antibody was previously described^31^. Anhydrotetracycline (aTc) was purchased from Cayman Chemical. DSM-1 was from Asinex.

### Generation of plasmids

To epitope tag the N-terminus of the SUB1 p31 at the endogenous locus in the previously reported transgenic PM X^apt^ parasites^12^, a human recodonized gene block was obtained from IDT. This gene block had the following elements in the order: signal peptide of SUB1-3xHA-remaining ORF of SUB1. To generate the donor vector for the CRISPR/ Cas9-mediated genome editing, 440 bp of the 5’UTR immediately upstream of the start *sub1* (left homologous region or LHR) and the 440 bp of the 3’UTR (right homologous region or RHR) were PCR amplified from the genomic DNA of the NF54 strain of *P. falciparum* using the primer pairs 272/ 273 and 270/ 271 respectively. The SUB1 gene block was PCR amplified using primer pair 274/ 275. An assembly PCR was performed using the primer pair 272/ 275 and a mixture of LHR, RHR, SUB1 gene block as template. The amplified fragment was subsequently cloned into the plasmid PM2GT^32^ to generate the plasmid PM2GT-HA-p31-SUB1. To generate SUB1 conditional knock out parasite line, a yPM2GT-SUB1^loxP^ plasmid was used following the principle described earlier^9^. The yPM2GT vector has been reported earlier^12^. First, the 5’ and 3’ homology arms were PCR amplified from NF54 gDNA using primer pairs 259/ 261 and 258/ 260 respectively. A Sera2 intron containing a loxP site in the middle^9^ was obtained as a gene block and further PCR amplified using the primer pair 262 and 264. A recodonized fragment of SUB1 ORF was obtained as a gene block and further PCR amplified using the primer pair 263/ 265. Finally, an assembly PCR was performed using the primer pair 259/ 265 and a mixture of 5’ and 3’ homology arms, sera2 intron and the recodonized fragment of SUB1 as the template. The amplified fragment was cloned into the yPM2GT plasmid that was digested with XhoI and NheI. A second Sera2 intron with loxP site was amplified using the primer pairs 276/ 277 and cloned into the above plasmid that was digested with SalI and EcoRI. This resulted into the final yPM2GT-SUB1^loxP^ plasmid. This plasmid was linearized with AflII and used as the donor vector for CRISPR/ Cas9 based genome editing.

To complement the SUB1 knockout, we modified the previously described plasmid pEOE-pAMA1-2X attP^13^. To make the expression of the second copy SUB1 stage-specific, we used a 2000 bp sequence located upstream of the PM X coding region (pPM X). This putative promoter-containing sequence was PCR amplified from the NF54 genomic DNA using the primer pair 278/279 and ligated between the AflII and XhoI restriction enzymes cut sites within pEOE-pAMA1-2X attP to generate the pEOE-pPM X-2X attP. The ORF of SUB1 was amplified using the primers 280 and 281. The amplified fragment was subsequently fused by Gibson assembly with the pEOE-pPM X-2X attP that was previously digested with XhoI and EagI. This resulted in the production of the final WT complementation construct pEOE-SUB1-V5-2XattP. Subsequent site-directed mutagenesis (QuickChange lightning, Agilent) on this plasmid used primers 266, 267, 268, and 269 to generate the different SUB1 cleavage mutant constructs.

To generate the bacteria expression plasmid to produce rp31 from *E. col*i, the pET28a plasmid (Novagen) was used. First, 3x HA sequence was PCR amplified from the yPM2GT-SUB1^loxP^ plasmid using the primer pair 280/281 and subsequently cloned between the NcoI and XhoI sites in the pET28a plasmid. A bacteria-recodonized version of the p31 sequence of SUB1 was then cloned into the modified pET28a at the XhoI site, thus labellig the N and the C termini of rp31 with 3x HA and 6x His respectively. The resultant pET28a-SUB1-p31 plasmid was used as a template to make the different rp31 cleavage site mutants by site directed mutagenesis using the primers 254, 255, 256.

Generation of the pHLSec-SUB1 plasmid to express SUB1 from the mammalian cells has been described earlier^12^. To epitope tag the N-terminus of p31, the 3x HA sequence was PCR amplified from the yPM2GT-SUB1^loxP^ plasmid using the primer pair 284/ 285 and cloned into the pHLSec-SUB1 plasmid at the AflIII site.

### Parasite culture, transfection and synchronization

NF54attB parasites with stably integrated Cre recombinase cassette (DiCre)^23^ and the resultant transgenic strains were cultured in human red blood cells using RPMI with 0.1% albumax, as previously described^33^. For the SUB1^loxP^ line, following electroporation, the transfectants were selected in the presence of 2μM DSM1^34^. For complementation of SUB1, plasmids carrying WT or mutant SUB1 coding sequences were independently co-transfected with a Bxb1 integrase plasmid^35^ into the SUB1^loxP^ parasites. Transfectants were selected with 5 nM WR99210^34^, together with DSM1. PM X^apt^ parasites^12^ were maintained in presence of 100 nM aTc and 5 μg/ml Blasticidin S^34^. For tagging the N-terminus of the endogenous SUB1, the PM X^apt^ parasites were co-transfected with the linearized plasmid PM2GT-HA-p31-SUB1 and the guide DNA plasmid. Transfected parasites were selected with 5 nM WR99210, together with 2.5 μg/ml blasticidin and 100 nM aTc. Integration of all plasmids were verified by PCR. For CRISPR/ Cas9 mediated editing of the *sub1* locus, we used the guide sequence: GCATTAACTAGTACATCAAA.

Highly synchronous ring-stage parasites were obtained as follows. High-parasitemia schizont cultures were passed through MACS LD magnet columns (Miltenyi Biotec) and schizonts were collected. These were then added to fresh uninfected RBCs resuspended in warm culture media. The cultures were shaken at 80 RPM for 3 h and the resulting parasites were synchronized using 5% sorbitol as described before^36^. To induce SUB1 knock out, synchronized ring-stage parasites (0-3 h old) with floxed *sub1* locus were treated with 10 nM rapamycin for 3 h. Parasites were then washed in prewarmed media and cultured to mature schizont stage (44 - 47 h post invasion). Genomic DNA was extracted and the excision of the floxed segment of SUB1 ORF was analyzed by PCR using the primer pair 282/ 283.

### Parasite growth assay

For growth curves, synchronized ring-stage parasites were diluted to an initial 1% parasitemia at 2% hematocrit. Cultures were split and rapamycin (10 nM) was added to appropriate wells. Control wells received equal volume of DMSO (vehicle). Parasitemia was determined every 24 hours using flow cytometry as described previously^12^.

### Expression and purification of recombinant SUB1 (rSUB1) from mammalian cells

rSUB1 was purified as a secreted protease from the mammalian cells using the protocol described before^22^, with some minor changes. Briefly, the Expi-HEK293 cells (Thermo Scientific) were transfected with the mammalian expression plasmid pHLSec containing the recodonized SUB1. For DNA delivery ExpiFectamine was used as per the manufacturer’s recommendation. Tunicamycin was added at a final concentration of 0.5 μg / ml immediately after transfection, and the cultures were grown for 72 hrs. Culture supernatants were then passed through a 0.22 μm filter and diluted with 3 culture volumes of 50 mM Tris-Cl pH 8.0, 100 mM NaCl, 10 mM imidazole. The diluted supernatant was purified using the Ni-NTA resin (Thermo Fisher). After extensive wash in the same buffer, the protein was eluted with 50 mM Tris-Cl pH 8.0, 100 mM NaCl, 400 mM imidazole. The eluted sample was then buffer exchanged using an Amicon ultra 3 kDa molecular wt. cutoff filter (Millipore, USA) and finally resuspended in 20 mM Tris pH 7.6, supplemented with glycerol to 10% (w/v), and stored in aliquots at −70 °C.

### Expression and purification of recombinant p31 (rp31) from E.coli

For rp31, the *E. coli* expression system was used. Briefly, the BL21 (DE3) competent cells were transformed with the pET28a plasmid containing the SUB1-p31 with 3xHA and 6xHis as the N and C-terminal tags respectively. 10 ml of bacterial culture was grown overnight at 37 °C, diluted to a 500 ml culture with a starting OD_600_ of 0.01. When the OD_600_ reached ~0.6, 250 uM IPTG was added, and the cultures were grown overnight at 18 °C. The cultures were then pelleted, and lysed in 0.2 g lysozyme, 100 mM Tris, PH 8.0 with 0.1% triton X-100 by sonication. The lysed bacteria were spun down at 12000 RPM for 10 mins. The lysate was diluted 20x in 100 mM NaCl, 50 mM Tris pH 8.0, 10 mM imidazole and passed over Ni-NTA column. The column was washed with 10 column volumes of buffer containing 100 mM NaCl, 50 mM Tris pH 8.0, 25 mM imidazole. The bound proteins were then eluted with 100 mM NaCl, 50 mM Tris pH 8.0, 500 mM imidazole. Finally, the proteins were buffer exchanged (100 mM NaCl, 50 mM Tris, pH 7.5) and concentrated with an Amicon ultra 3 kDa molecular wt. cutoff filter (Millipore, USA). The concentrated proteins were then immediately used for *in vitro* rPM X cleavage assay.

### In vitro rPM X cleavage assay

Activity of rPM X towards rp31 was carried out as previously described^13^ with minor modifications. Briefly, 250 ng of rPM X was incubated with 1 ug of rp31 in 40 ul of PM X activity buffer (150 mM NaCl, 25 mM MES, 25 mM Tris, pH 5.5) for 3 hours at 37 °C. To inhibit PM X activity, 1 μM CW-117^12,37^ was added to samples at pH 5.5. Following incubation, samples were mixed with 4x sample loading buffer and heated at 99 °C for 5 min. Fractions of equal volumes were subjected to SDS-PAGE and immunoblotting.

For cleavage of rSUB1 by rPM X, at first the former was immobilized onto His trap magnetic sepharose beads as per the manufacturer’s protocol. Protein-bound beads were washed 2x in 150 mM NaCl, 50 mM Tris, pH 7.6, and subsequently resuspended in PM X activity buffer. rPM X and CW-117 were added to appropriate tubes and reactions were incubated as above. Following incubation, the tubes were placed on magnet, and the supernatants were separated. The bead-bound proteins were eluted with buffer containing 300 mM NaCl, 50 mM Tris, 500 mM imidazole, pH 8.0. Both the supernatant and the eluted fractions were processed as above.

### Immunoprecipitation and western blot

To immunoprecipitate p31 from parasite lysates, highly synchronized PM X^apt^ parasites expressing N-terminally 3x HA tagged SUB1 were split into two plates. To one plate, aTc was added, and to the other equal volume of vehicle (DMSO) was added. 46 h post invasion, free schizonts were collected by saponin permeabilzation. Samples were subsequently lysed in 150 mM NaCl, 50 mM Tris, pH 7.5, 1% NP-40 supplemented with protease inhibitor cocktail. Insoluble material was removed by centrifugation. Clarified lysates were mixed with the dynabeads coated with mouse anti-HA antibody. Immunoprecipitation was carried out as per the manufacturer’s recommendation. Both the immunoprecipitated and the flow through fractions were mixed with 4x sample loading buffer and heated at 99 °C for 5 min. Aliquots of equal cell equivalents were subjected to SDS-PAGE and immunoblotting.

For SUB1 knockout western blots, following synchronization, the SUB1^loxP^ parasites expressing a second copy SUB1 or not were cultured in presence or absence of rapamycin as in the parasite growth assays, except the parasitemia was maintained around 5%. 46 hours post invasion samples were harvested as above, lysed with RIPA buffer, and insoluble fractions were removed by centrifugation. Cell lysates were then directly boiled with SDS-PAGE loading buffer followed by western blot.

Primary antibodies included rabbit anti-HA (1:1000), rabbit anti-SUB1 (1:1000), rabbit anti-HA (1:1000), rabbit anti-V5 (1:1000), mouse anti PM V (1:500), mouse anti-his (1:1000), and rat anti-FLAG (1:1000). For all, appropriate IRDye conjugated secondary antibodies were used at 1:10000 dilution. Blots were visualized on an Odyssey imaging system (Licor).

### Immunofluorescence assays (IFAs)

IFAs were performed as before^13^. Briefly, synchronized 44-47 h old schizont cultures were smeared on glass slides, fixed in 3.7 % paraformaldehyde for 15 min and blocked in 3 % BSA in PBS overnight at 4°C before antibody staining. The antibodies used for IFA were: rabbit anti-HA (1:500), mouse anti-V5 (1:500), mouse anti-BiP (1:500). The secondary antibodies were used as 1:2000 dilutions and were conjugated to Alexa Fluor 488 or 546 (Life Technologies). Cells were mounted with ProLong and 4’,6’-diamidino-2-phenylindole (DAPI) (Invitrogen) and imaged using a Zeiss Imager M2 Plus wide field fluorescence microscope, using a 63x objective and the Axiovision 4.8 software for epifluorescence imaging.

### Statistical analysis

Unless specified otherwise, all values in all figures are averaged from three independent biological repeats and error bars represent standard deviations. Differences were assessed by the two-tailed Student’s *t* test using the Microsoft Excel software. P values indicating statistical significance were grouped in all figures. All graphs were prepared using GraphPad Prism 8.

## Acknowledgments

This work was supported by grant AI138447 from the National Institute of Allergy and Infectious Diseases (NIAID) to D.E.G. We thank Dr. Josh Beck (Iowa State University) for the NF54-DiCre parasite line, Dr. Michael J. Blackman (The Francis Crick Institute, London) for anti-SUB1 antibody, Dr. Marvin J. Meyers (Saint Louis University) for CW-117, the protein facility of the Iowa State University for the Edman degradation assays. WR99210 was a gift from D. Jacobus from Jacobus Pharmaceutical Co. We also thank Barbara Vaupel for assistance with cloning, and Dr. Eva Istvan for useful suggestions.

## Notes

**Competing Interest Statement:** All authors declare no competing interests.

### Competing Interest Statement

The authors have declared no competing interest.

## References

1. WHO. WHO World Malaria Report 2020. Malaria report (2020).

2. Glushakova, S. et al. Rounding precedes rupture and breakdown of vacuolar membranes minutes before malaria parasite egress from erythrocytes. Cell. Microbiol. 20, e12868 (2018).

3. Glushakova, S., Yin, D., Li, T. & Zimmerberg, J. Membrane transformation during malaria parasite release from human red blood cells. Curr. Biol. (2005) doi:10.1016/j.cub.2005.07.067.

4. Glushakova, S. et al. New stages in the program of malaria parasite egress imaged in normal and sickle erythrocytes. Curr. Biol. 20, 1117–1121 (2010).

5. Dvorin, J. D. et al. A plant-like kinase in Plasmodium falciparum regulates parasite egress from erythrocytes. Science 328, 910–912 (2010).

6. Collins, C. R. et al. Malaria Parasite cGMP-dependent Protein Kinase Regulates Blood Stage Merozoite Secretory Organelle Discharge and Egress. PLoS Pathog. 9, (2013).

7. Absalon, S. et al. Calcium-Dependent Protein Kinase 5 Is Required for Release of Egress-Specific Organelles in Plasmodium falciparum. MBio 9, (2018).

8. Yeoh, S. et al. Subcellular Discharge of a Serine Protease Mediates Release of Invasive Malaria Parasites from Host Erythrocytes. Cell (2007) doi:10.1016/j.cell.2007.10.049.

9. Thomas, J. A. et al. A protease cascade regulates release of the human malaria parasite Plasmodium falciparum from host red blood cells. Nat. Microbiol. (2018) doi:10.1038/s41564-018-0111-0.

10. Das, S. et al. Processing of Plasmodium falciparum Merozoite Surface Protein MSP1 Activates a Spectrin-Binding Function Enabling Parasite Egress from RBCs. Cell Host Microbe (2015) doi:10.1016/j.chom.2015.09.007.

11. Sajid, M., Withers-Martinez, C. & Blackman, M. J. Maturation and specificity of Plasmodium falciparum subtilisin-like protease-1, a malaria merozoite subtilisin-like serine protease. J. Biol. Chem. 275, 631–641 (2000).

12. Nasamu, A. S. et al. Plasmepsins IX and X are essential and druggable mediators of malaria parasite egress and invasion. Science (80-.). 358, 518–522 (2017).

13. Mukherjee, S., Nguyen, S., Sharma, E. & Goldberg, D. E. Maturation and substrate processing topography of the Plasmodium falciparum invasion/egress protease plasmepsin X. Nat. Commun. 13, 4537 (2022).

14. Pino, P. et al. A multistage antimalarial targets the plasmepsins IX and X essential for invasion and egress. Science (80-.). (2017) doi:10.1126/science.aaf8675.

15. Favuzza, P. et al. Dual Plasmepsin-Targeting Antimalarial Agents Disrupt Multiple Stages of the Malaria Parasite Life Cycle. Cell Host Microbe (2020) doi:10.1016/j.chom.2020.02.005.

16. Yabuta, Y., Takagi, H., Inouye, M. & Shinde, U. Folding pathway mediated by an intramolecular chaperone: propeptide release modulates activation precision of pro-subtilisin. J. Biol. Chem. 276, 44427–44434 (2001).

17. Saouros, S. et al. Microneme protein 5 regulates the activity of Toxoplasma subtilisin 1 by mimicking a subtilisin prodomain. J. Biol. Chem. 287, 36029–36040 (2012).

18. Brydges, S. D. et al. Targeted deletion of MIC5 enhances trimming proteolysis of Toxoplasma invasion proteins. Eukaryot. Cell 5, 2174–2183 (2006).

19. Jean, L., Hackett, F., Martin, S. R. & Blackman, M. J. Functional characterization of the propeptide of Plasmodium falciparum subtilisin-like protease-1. J. Biol. Chem. 278, 28572–28579 (2003).

20. Withers-Martinez, C. et al. The malaria parasite egress protease SUB1 is a calcium-dependent redox switch subtilisin. Nat. Commun. 5, 3726 (2014).

21. Giganti, D. et al. A novel Plasmodium-specific prodomain fold regulates the malaria drug target SUB1 subtilase. Nat. Commun. 5, 4833 (2014).

22. Withers-Martinez, C. et al. Expression of recombinant Plasmodium falciparum subtilisin-like protease-1 in insect cells. Characterization, comparison with the parasite protease, and homology modeling. J. Biol. Chem. 277, 29698–29709 (2002).

23. Fierro, M. A., Hussain, T., Campin, L. J. & Beck, J. R. Knock-sideways by inducible ER retrieval reveals a novel extra-vacuolar function for <em>Plasmodium</em> PTEX component HSP101. bioRxiv 2022.10.02.510311 (2022) doi:10.1101/2022.10.02.510311.

24. Jones, M. L. et al. A versatile strategy for rapid conditional genome engineering using loxP sites in a small synthetic intron in Plasmodium falciparum. Sci. Rep. 6, 21800 (2016).

25. Anderson, E. D. et al. The ordered and compartment-specfific autoproteolytic removal of the furin intramolecular chaperone is required for enzyme activation. J. Biol. Chem. 277, 12879–12890 (2002).

26. Power, S. D., Adams, R. M. & Wells, J. A. Secretion and autoproteolytic maturation of subtilisin. Proc. Natl. Acad. Sci. U. S. A. 83, 3096–3100 (1986).

27. Kluskens, L. D. et al. Molecular characterization of fervidolysin, a subtilisin-like serine protease from the thermophilic bacterium Fervidobacterium pennivorans. Extremophiles 6, 185–194 (2002).

28. Strausberg, S., Alexander, P., Wang, L., Schwarz, F. & Bryan, P. Catalysis of a protein folding reaction: thermodynamic and kinetic analysis of subtilisin BPN’ interactions with its propeptide fragment. Biochemistry 32, 8112–8119 (1993).

29. Tanaka, S. et al. Crystal structure of unautoprocessed precursor of subtilisin from a hyperthermophilic archaeon: evidence for Ca2+-induced folding. J. Biol. Chem. 282, 8246–8255 (2007).

30. Shinde, U. & Thomas, G. Insights from bacterial subtilases into the mechanisms of intramolecular chaperone-mediated activation of furin. Methods Mol. Biol. 768, 59–106 (2011).

31. Banerjee, R. et al. Four plasmepsins are active in the Plasmodium falciparum food vacuole, including a protease with an active-site histidine. Proc. Natl. Acad. Sci. U. S. A. 99, 990–995 (2002).

32. Klemba, M., Beatty, W., Gluzman, I. & Goldberg, D. E. Trafficking of plasmepsin II to the food vacuole of the malaria parasite Plasmodium falciparum. J. Cell Biol. 164, 47–56 (2004).

33. Istvan, E. S. et al. Esterase mutation is a mechanism of resistance to antimalarial compounds. Nat. Commun. 8, 14240 (2017).

34. Istvan, E. S. et al. Plasmodium niemann-pick type C1-related protein is a druggable target required for parasite membrane homeostasis. Elife 8, 1–23 (2019).

35. Nkrumah, L. J. et al. Efficient site-specific integration in Plasmodium falciparum chromosomes mediated by mycobacteriophage Bxb1 integrase. Nat. Methods 3, 615–621 (2006).

36. Lambros, C. & Vanderberg, J. P. Synchronization of Plasmodium falciparum erythrocytic stages in culture. J. Parasitol. 65, 418–420 (1979).

37. Meyers, M. J. et al. Evaluation of aminohydantoins as a novel class of antimalarial agents. ACS Med. Chem. Lett. 5, 89–93 (2014).

