## Supplementary figures for "Activation of the *Plasmodium* egress effector subtilisin-like protease 1 is achieved by plasmepsin X destruction of the propiece"

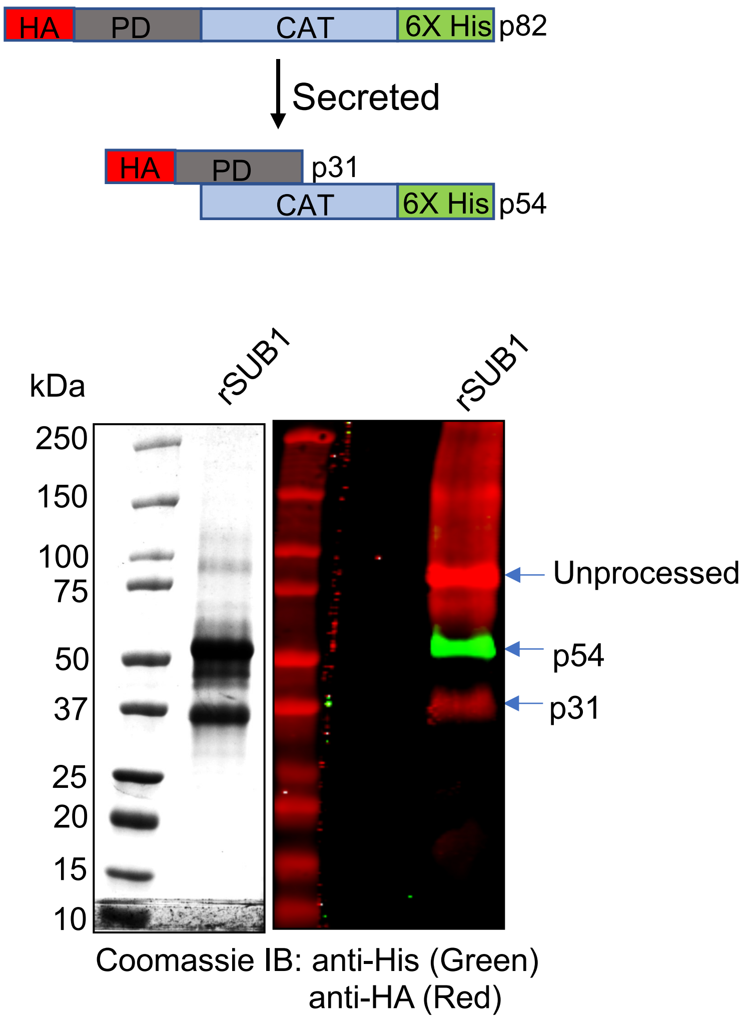


**Supplementary figure 1: Purified recombinant SUB1 (rSUB1) from mammalian Expi293 cells.** As described in the methods, rSUB1, that was tagged at the N and C terminal ends with 3x HA and 6x His tags were purified as a secreted protein. Treatment of the transfected cells with tunicamycin resulted in the correct folding of rSUB1, which was then secreted mostly as the semi-processed p31/p54 complex.

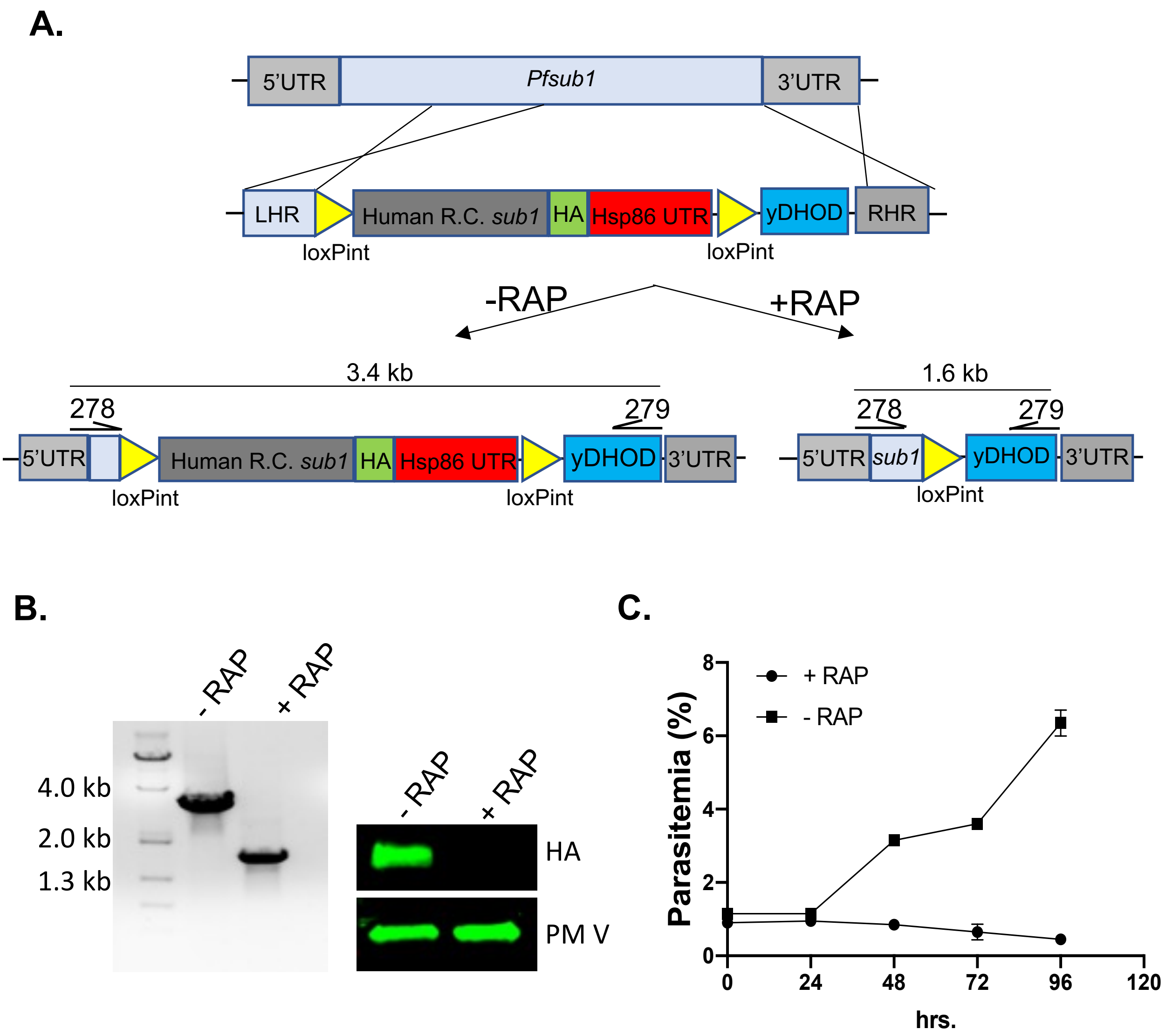


**Supplementary figure 2: Generation of the rapamycin (RAP) inducible SUB1 knock out (KO) parasite line.** (**A**) Schematic of the linear donor vector used to replace the endogenous locus with an HA-tagged human recodonized *sub1* by CRISPR/ Cas9 mediated genome editing. Also shown is the modified locus before and after treatment with RAP. Positions of the forward (278) and the reverse (279)primers used to check genome editing by PCR are shown. (**B**) Left: PCR, right: western blot showing excision of a segment of SUB1 ORF and the resulted loss of protein expression (HA) respectively. As a loading control for the western blot, expression of the ER resident protease plasmepsin V (PM V) was used. (**C**) Replication of mock (DMSO) treated (-RAP) or RAP treated (+RAP) parasites over a period of 96 h. Experiment was performed two times. Shown are the mean values, error bars represent standard deviation.

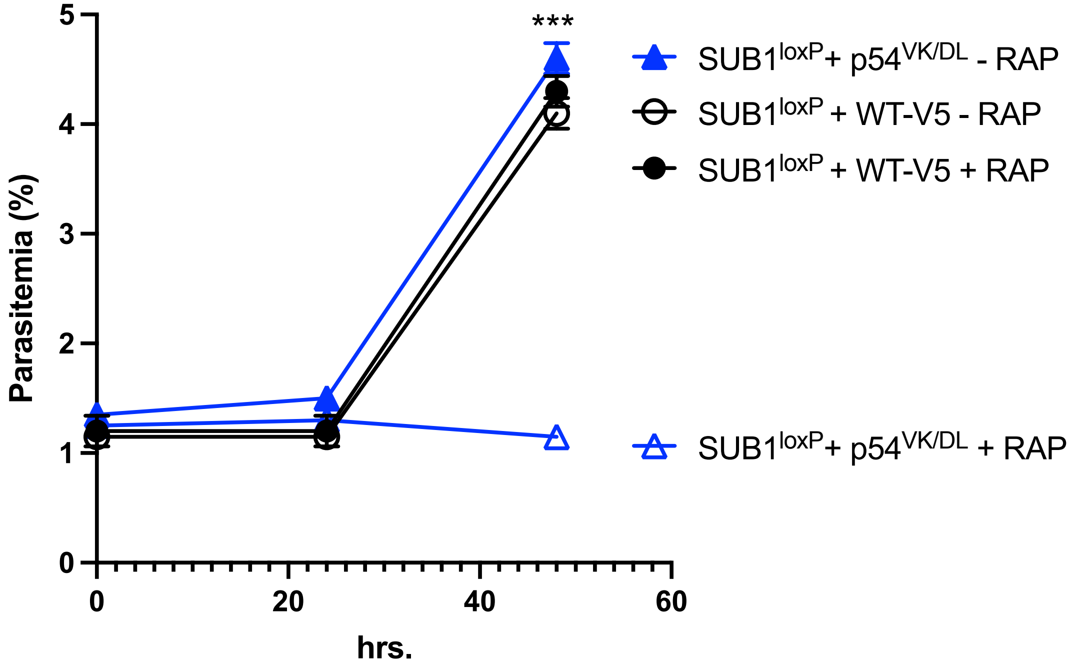


**Supplementary figure 3: Primary autocatalytic processing of SUB1 is essential for parasite growth.** Synchronized parasites from the Indicated lines were cultured were cultured as in figure 5. Mean values from three independent experiments are shown and error bars represent standard deviations. Statistical two-tailed Student's *t*-test was done to determine the significance of difference in growth between the $54^{VK/DL}$ mutant parasites grown in presence and absence of RAP . ***: $p < 0.001$.

### Supplementary table 1: List of primers used in this study.

| Primer # | Sequence | Notes |
|---|---|---|
| 254 | GATTGTGTCGGAATTACGCGCCGCAGAGAAGGTGGAGGATGTTATT | mutagenesis primer for making RFLE to RFAA in pET28a |
| 255 | GGAACCCGACGTATTGTCTTTCGCCGCATCTAAGGGCAACTTGTCAAAC | mutagenesis primer for making SFLE to AFAA in pET28a |
| 256 | GAATCATGCTACTACCCCCAGTTTTGCCGCAGAAAGTCTTTTGGAAC | mutagenesis primer for making SFFQ to SFAA in pET28a |
| 258 | GACACTATAGAACTCGAGTGGAAATGGCTTTTTAAATAATAT | forward PCR primer for amplification of SUB1 RHR from gDNA |
| 259 | GTTAATATTAGCAAGCACGTCTTAAGATGATGCTCAATAAAAAAGTTG | forward PCR primer for amplification of SUB1 LHR from gDNA |
| 260 | CAACTTTTTTATTGAGCATCATCTTAAGACGTGCTTGCTAATATTAAC | Reverse PCR primer for amplification of SUB1 RHR from gDNA |
| 261 | GTATATTATTTTTTTTATTTACCTTCCTTTAGTTCTATAATCAT | Reverse PCR primer for amplification of SUB1 LHR from gDNA |
| 262 | CAAATGTTTAGATTATTGTGTAAATAAAAAAAATAATATA | forward primer for amplification of sera2 intron |
| 263 | ATTTTATATTCTTTTAGATATCCCGCAATGCGCATATG | forward primer to assemble human rcoded SUB1 |
| 264 | CATATGCGCATTGCGGGATATCTAAAAGAATATAAAAT | reverse primer for amplification of sera2 intron |
| 265 | GTCCGGGACGTCGTACGGGTAGCTAGCATGGAGGTACCTACTCTTCTT | reverse primer to assemble human recoded SUB1 |
| 266 | GTTAATGATTATAAAAGTATGGCCGCAGTCGAAAATGATGCTGAAGAT | mutagenesis primer for SMLE to SMAA in pEOE |
| 267 | GATTATAAAAGTATGTTAGAAAAAGAAAATCTTGCTGAAGATTATGATAAAATG | mutagenesis primer for SUB1 p47 autocleavage mutant in pEOE |
| 268 | GTTAATGATTATAAAAGTATGGCCGCAAAAGAAAATCTTGCTGAAGATTATGATAAAATG | mutagenesis primer for SUB1 SMAA + p47 autocleavage mutant in pEOE |
| 269 | GATTGAATCAGATAAATTAAAAAGTGCACTTAATATTGATATAAGTGGTAT | mutagenesis primer for SUB1 p54 autocleavage mutant in pEOE |
| 270 | GATTTAGGTGACACTATAGAACTTGGAAATGGCTTTTTAAATAATAT | forward primer for SUB1 RHR in PM2GT |
| 271 | TATATTTGTATTTGTACCTTAAGACGTGCTTGCTAATATTAAC | reverse primer for SUB1 RHR in PM2GT |
| 272 | GTTAATATTAGCAAGCACGTCTTAAGGTACAAATACAAATATAAAT | forward primer for SUB1 LHR in PM2GT |
| 273 | CAACTTTTTTATTGAGCATCATGGTTTAAAAAAAAAATAAAATAAAA | reverse primer for SUB1 LHR in PM2GT |
| 274 | TTTTATTTTATTTTTTTTTTAAACCATGATGCTCAATAAAAAAGTTG | forward primer for SUB1 gblock in PM2GT |
| 275 | GACGCGGCCGTCACGTACGTCAATGGAGGTACCTACTCTTCTTTTTCC | reverse primer for SUB1 gblock in PM2GT |
| 276 | AAGCTTGCATGCCTGCAGGGTAAATAAAAAAAATAATATA | forward primer for 2nd sera2int |
| 277 | CATATTTATTAAATCTAGAATTCTAAAAGAATATAAAATA | reverse primer for 2nd sera2int |
| 278 | GTGCCACCTGACGTCGTTAAATAATTATTAATTTTCC | forward primer for PM X promoter amplification |
| 279 | CAACTTTTTTATTGAGCATCATCTCGAGTTCTGTCTTTATAACCG | reverse primer for PM X promoter amplification |
| 280 | AGTGGTGGTGGTGGTGGTGCTCGAGATCCGCACTAACCAGTTTA | forward primer for HA insertion in pET28a |
| 281 | AGGAGATATACCATGGTCGACACTATATACCCTTATGACGTTCCAGAC | reverse primer for HA insertion in pET28a |
| 282 | TCCATGTACCTAGGTGTGTG | SUB1loxP forward primer |
| 283 | AGATCAGTTCCGTGGGAAG | SUB1loxP reverse primer |
| 284 | GCGTAGCTGAAACCGGTTACCCTTATGACGTTCCAGACTAC | forward primer for adding 3x HA in pHLSec-SUB1 |
| 285 | CCGTTTTCCTCGGACCTCACTTCCTTAGCGTAATCAGGAACGTC | reverse primer for adding 3x HA in pHLSec-SUB1 |
