## Additional supplementary file 1 for "Activation of the *Plasmodium* egress effector subtilisin-like protease 1 is achieved by plasmepsin X destruction of the propiece"

### Iowa State University
### Protein Facility

#### PROTEIN/PEPTIDE SEQUENCE REPORT

**Date:** August 8, 2022

**To:** Sumit Mukherjee

**Sample Number:** 10022

**Sample Name:** 2

**Sample Preparation:** The membrane was washed with DI water and loaded onto the instrument for sequence analysis.

**Instrument:** Shimadzu PPSQ-53A

**Sequencing Method:** Edman Degradation

| Cycle Number | Amino Acid |
|---|---|
| 1 | M |
| 2 | R |
| 3 | E |
| 4 | L |

The major amino acid is listed first for each cycle. If you have any questions, feel free to contact me.

Prepared by: Joel Nott

### Protein/Peptide Sequencing Submission Form

**Tracking Number-1550:** 10022 Date: Aug-03-2022

Your sample ID 2

| | | |
|---|---|---|
| Name: SUMIT MUKHERJEE | Login: sumitmukh | Principal Investigator: Daniel Goldberg |
| Department/Company: Washington University School of Medicine | Phone #: 8062245077 | Fax #: |
| E-Mail Address: | Principal Investigator E-Mail Address: | |
| Mailing Address: 660 South Euclid Avenue, Department of Molecular Microbiology, Saint Louis, Missouri, 63110 | | |
| Account #: PR00136711 | | Assignee: |
| Business Purpose: | | |
| Billing Contact Name: Rachel Warhover | | |
| Billing Mailing Address: 4990 Children's Place, Infectious Diseases Division, Saint Louis, Missouri, 63110 | | |
| Billing Phone #: 314-454-8225 | Billing E-Mail Address: | |

✓ I agree to the terms and conditions present at http://www.biotech.iastate.edu/facilities/Agreements/ProteinTechnicalServicesAgreement.pdf

How many residues do you need? 4

**Sample Information**

Sample amount________moles;or ________microgramsM.W.________

For samples in solution: What solvent is the sample in?________

For samples electroblotted toPVDF: What membrane was used?

____Immobilon-P (.45 micron) (Millipore) ____Problot (.1 micron) (ABI)
____Westran (.45 micron) (Schleicher & Schuell) ____Trans-Blot (.1 micron) (Biorad)
____Immobilon-PSQ (.1 micron) (Millipore) ____Fluoratrans(.1 micron) (Pall Corp)

N-Terminal blocked: No________Do not know________ Yes________

Protein/PeptideModified: Yes, at________ with________

Cysteine modified: Yes________ If yes, what derivative?________ No________

Enzyme treatment: Yes________ What enzyme?________ Cleavage sites________

Radioactivity: Yes________ No________

Protein sequence known: Yes______ No______ DNA sequence known: Yes______No______

Describe purification stepsin detail, especially possible contaminants such as buffer, salts, and SDS:

If your sample was collectedon an HPLC, please attachthe chromatogram with AUFS, gradient, solvents, column and wavelength.

[Sequence Analysis]

| | | |
|---|---|---|
| Data Acquired | : | 8/7/2022 12:32:56 PM |
| Data Processed | : | 8/8/2022 7:25:40 AM |
| Reactor | : | 2 |
| Number of Cycles | : | 5 |
| Sequence Schedule | : | C:\PPSQ\SeqProg3_PDA_BGE\PVDF9-3G.sch |
| Sample Name | : | Sumit Mukherjee, 2 |
| Sample Amount(pmol) | : | 10.0 |
| Sample ID | : | 10022 |
| Operator Name | : | System Administrator |
| Data File | : | 10022_08-07-2022 |
| Start Number | : | 1 |
| Method File | : | 10022_08-07-2022.lcm |
| Batch File | : | 10022_08-07-2022.lcb |
| Data Folder Path | : | C:\LabSolutions\Data\Project1\PPSQ\10022_08-07-2022 |
| Number of Analyses | : | 5 / 5 |
| Standard File | : | C:\LabSolutions\Data\Project1\PPSQ\10022_08-07-2022\PTH-AA_08-04-2022_D01.lcd |
| Data Comment | | |

[Sequence]

M R E L

[Estimated Sequence]

| | 1 | 2 | 3 | 4 |
|---|---|---|---|---|
| 1st | M | R | E | L |
| 2nd | V | I | Y | K |
| 3rd | F | Q | G | N |
| 4th | S | D | K | W |
| Reliability(%) | 100.0 | 33.8 | 59.5 | 91.6 |

[Evaluated Value]

| | 1 | 2 | 3 | 4 |
|---|---|---|---|---|
| D | 0.45 | 131.06 | 0.63 | 0.81 |
| E | 0.29 | 96.45 | 4279.12 | 0.53 |
| N | 0.64 | 55.40 | 106.56 | 72.07 |
| S | 2.10 | 0.87 | 106.63 | 0.93 |
| T | 0.67 | 73.33 | 76.97 | 0.78 |
| Q | 0.38 | 131.38 | 0.58 | 0.87 |
| G | 0.68 | 63.75 | 235.22 | 0.73 |
| H | 0.77 | 1.00 | 4.00 | 0.76 |
| A | 0.67 | 0.99 | 0.72 | 0.90 |
| R | 0.09 | 1196.95 | 0.52 | 0.34 |
| Y | 0.79 | 18.13 | 765.12 | 0.31 |
| P | 0.35 | 18.99 | 85.04 | 0.32 |
| M | 3777.23 | 0.42 | 0.07 | 0.16 |
| V | 378.38 | 0.45 | 0.39 | 1.00 |

| | | | | |
|---|---|---|---|---|
| W | 0.49 | 0.83 | 0.89 | 2.16 |
| K | 0.33 | 74.37 | 115.06 | 622.33 |
| F | 52.96 | 0.37 | 54.74 | 0.41 |
| I | 0.07 | 354.46 | 0.46 | 0.09 |
| L | 0.76 | 82.85 | 0.53 | 5254.12 |

[Amount Yield(pmol)]

| | 1 | 2 | 3 | 4 |
|---|---|---|---|---|
| D | 5.77 | 6.62 | 1.63 | 0.00 |
| E | 1.59 | 3.61 | 123.63 | 0.00 |
| N | 1.75 | 1.36 | 2.08 | 90.04 |
| S | 3.38 | 0.00 | 6.65 | 74.77 |
| T | 2.76 | 5.11 | 3.43 | 0.00 |
| Q | 1.56 | 6.19 | 0.00 | 0.00 |
| G | 4.16 | 6.68 | 7.13 | 0.00 |
| H | 0.88 | 12.58 | 5.79 | 15.72 |
| A | 3.98 | 15.41 | 24.37 | 42.52 |
| R | 4.35 | 26.27 | 16.89 | 0.00 |
| Y | 3.14 | 12.09 | 48.31 | 0.00 |
| P | 0.00 | 0.00 | 78.66 | 0.00 |
| M | 90.65 | 0.00 | 27.28 | 0.00 |
| V | 8.40 | 54.36 | 99.50 | 17.19 |
| W | 2.91 | 32.67 | 11.57 | 4.13 |
| K | 6.55 | 65.63 | 22.15 | 17.56 |
| F | 0.00 | 83.11 | 25.35 | 8.19 |
| I | 2.90 | 68.56 | 0.00 | 0.00 |
| L | 7.47 | 68.69 | 0.00 | 88.05 |

[Percent Yield]

| | |
|---|---|
| Amino Acid | : A,V,L |
| Initial Yield(%) | : 880.53 |
| Repetitive Yield(%) | : 100.00 |
| Correlation Coef. | : 1.000 |
| Number of Data | : 1 |

[Repetitive Yield(%)]

Data File : PTH-AA_08-04-2022_D01.lcd
Sample Name : PTH-AA
Method File : PTH-AA_08-04-2022.lcm
Background Data File :

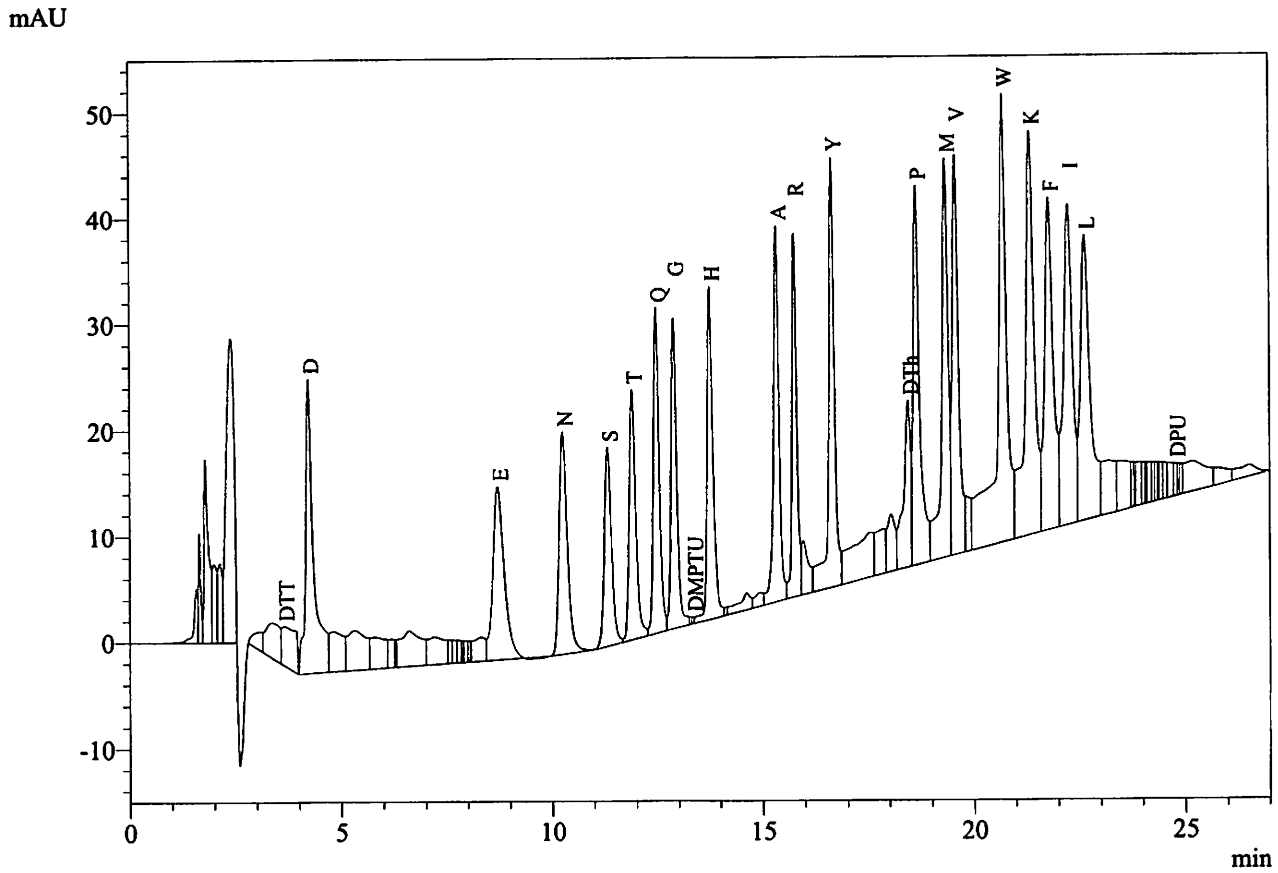


Peak Table
PDA Ch1 269nm

| Peak# | Name | Ret. Time | Area | Conc. |
|---|---|---|---|---|
| 9 | DTT | 3.639 | 86838 | 10.000 |
| 10 | D | 4.229 | 368538 | 10.000 |
| 26 | E | 8.709 | 301709 | 10.000 |
| 27 | N | 10.248 | 283149 | 10.000 |
| 28 | S | 11.321 | 219871 | 10.000 |
| 29 | T | 11.902 | 253806 | 10.000 |
| 30 | Q | 12.472 | 292815 | 10.000 |
| 31 | G | 12.889 | 273666 | 10.000 |
| 33 | DMPTU | 13.333 | 2040 | 10.000 |
| 34 | H | 13.739 | 303815 | 10.000 |
| 38 | A | 15.333 | 320725 | 10.000 |
| 39 | R | 15.757 | 270026 | 10.000 |
| 41 | Y | 16.643 | 398165 | 10.000 |
| 45 | DTh | 18.456 | 184729 | 10.000 |
| 46 | P | 18.650 | 382080 | 10.000 |
| 47 | M | 19.342 | 409815 | 10.000 |
| 48 | V | 19.578 | 389719 | 10.000 |
| 50 | W | 20.706 | 688571 | 10.000 |
| 51 | K | 21.353 | 578793 | 10.000 |
| 52 | F | 21.795 | 440406 | 10.000 |
| 53 | I | 22.255 | 445619 | 10.000 |
| 54 | L | 22.643 | 434982 | 10.000 |
| 69 | DPU | 24.779 | 16703 | 10.000 |
| Total | | | 7346580 | |

| | |
|---|---|
| Data File | : 10022_08-07-2022_D01.lcd |
| Sample Name | : Sumit Mukherjee, 2 |
| Method File | : 10022_08-07-2022.lcm |
| Background Data File | : |

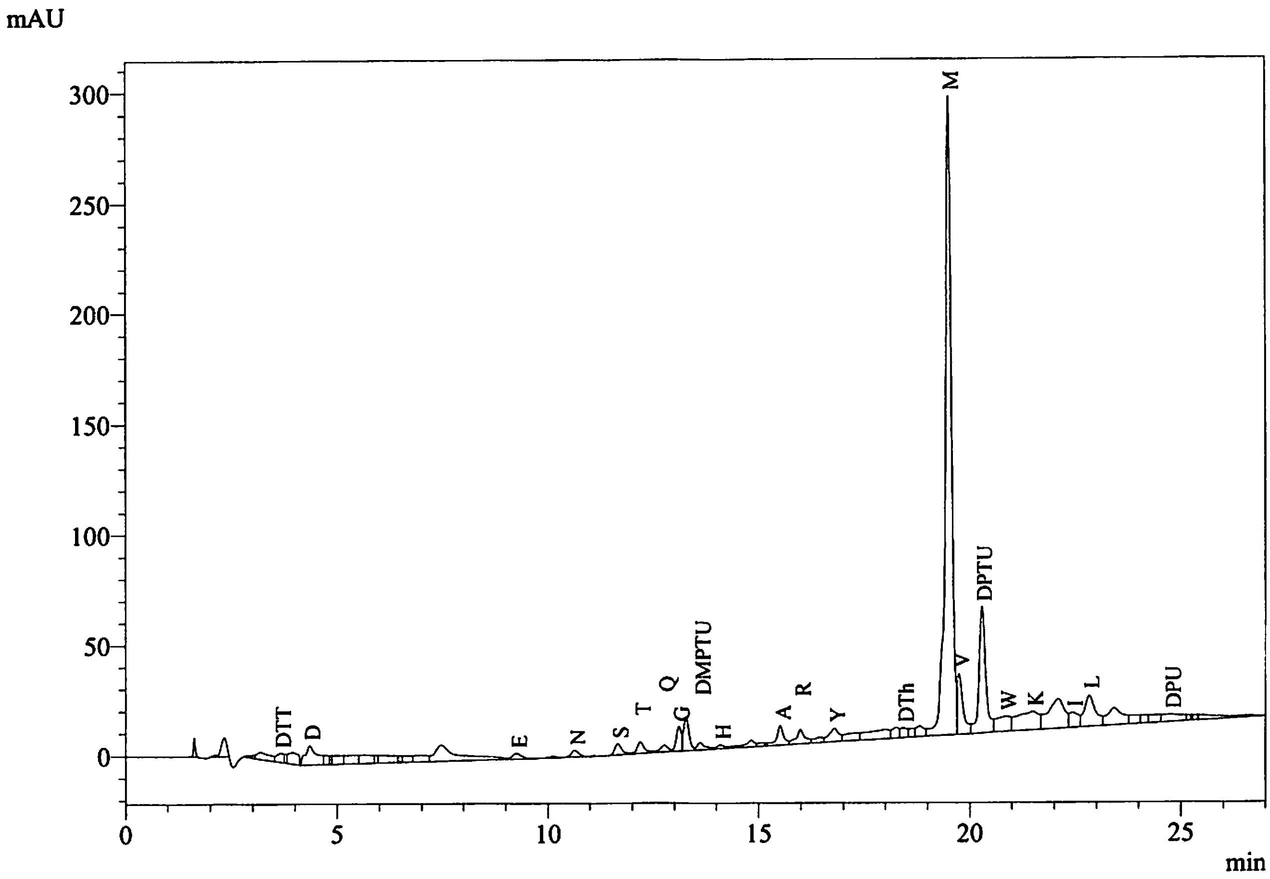


Peak Table
PDA Ch1 269nm

| Peak# | Name | Ret. Time | Area | Conc. |
|---|---|---|---|---|
| 6 | DTT | 3.652 | 49327 | 7.100 |
| 9 | D | 4.356 | 170151 | 5.771 |
| 22 | E | 9.248 | 38323 | 1.588 |
| 25 | N | 10.636 | 39561 | 1.746 |
| 27 | S | 11.654 | 59482 | 3.382 |
| 28 | T | 12.187 | 56008 | 2.758 |
| 30 | Q | 12.758 | 36545 | 1.560 |
| 31 | G | 13.099 | 90979 | 4.156 |
| 33 | DMPTU | 13.602 | 42583 | 260.863 |
| 34 | H | 14.087 | 21405 | 0.881 |
| 38 | A | 15.507 | 102028 | 3.976 |
| 39 | R | 15.983 | 94059 | 4.354 |
| 41 | Y | 16.792 | 100060 | 3.141 |
| 45 | DTh | 18.429 | 51909 | 3.513 |
| 48 | M | 19.511 | 2972116 | 90.654 |
| 49 | V | 19.747 | 261995 | 8.403 |
| 50 | DPTU | 20.293 | 638606 | |
| 51 | W | 20.865 | 160327 | 2.910 |
| 52 | K | 21.498 | 303176 | 6.548 |
| 54 | I | 22.434 | 103241 | 2.896 |
| 55 | L | 22.829 | 259834 | 7.467 |
| 60 | DPU | 24.758 | 106720 | 79.865 |
| Total | | | 5758436 | |

| | |
|---|---|
| Data File | : 10022_08-07-2022_D02.lcd |
| Sample Name | : Sumit Mukherjee, 2 |
| Method File | : 10022_08-07-2022.lcm |
| Background Data File | : 10022_08-07-2022_D01.lcd |

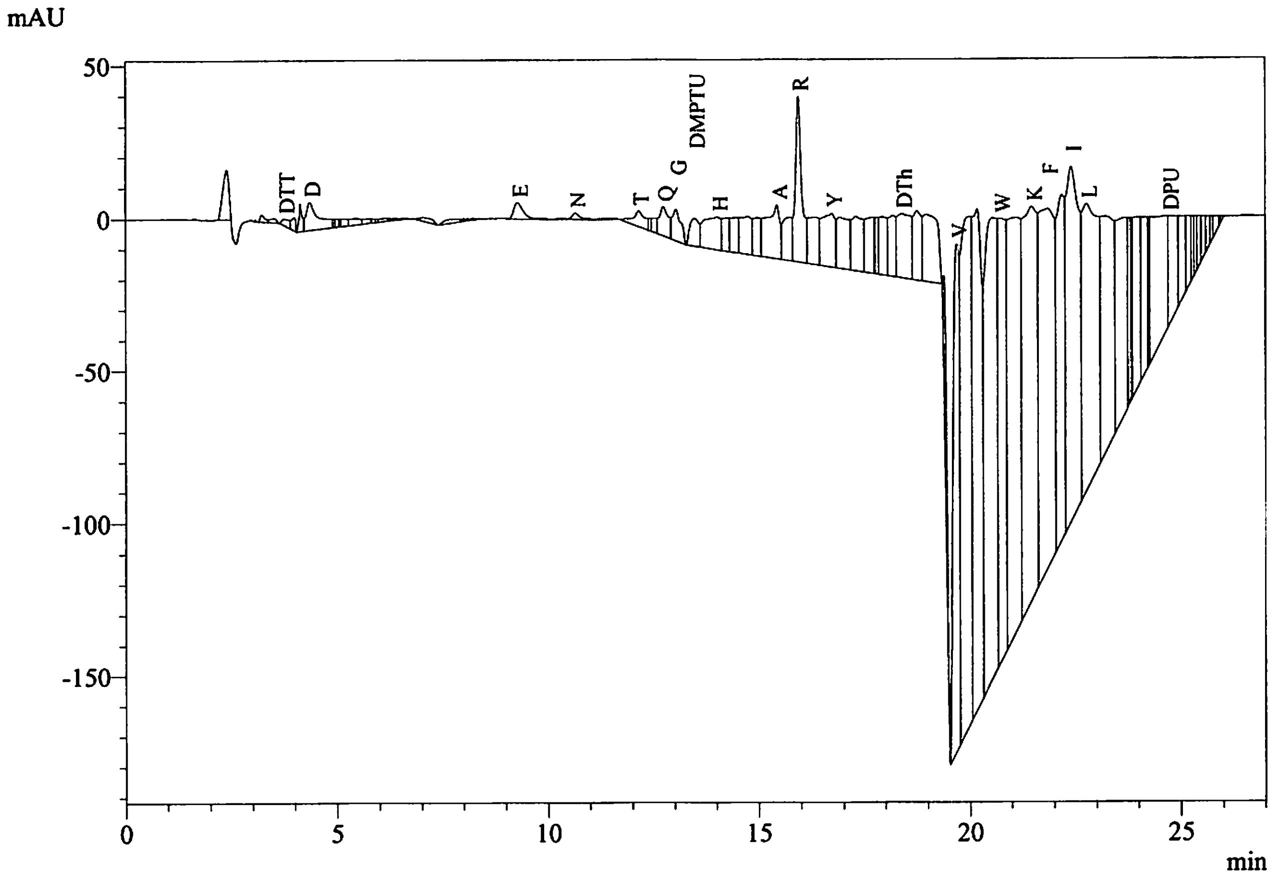


Peak Table
PDA Ch1 269nm

| Peak# | Name | Ret. Time | Area | Conc. |
|---|---|---|---|---|
| 5 | DTT | 3.749 | 28342 | 4.080 |
| 8 | D | 4.364 | 195321 | 6.625 |
| 25 | E | 9.285 | 87034 | 3.606 |
| 28 | N | 10.653 | 30719 | 1.356 |
| 33 | T | 12.160 | 103818 | 5.113 |
| 36 | Q | 12.744 | 145066 | 6.193 |
| 37 | G | 13.043 | 146327 | 6.684 |
| 38 | DMPTU | 13.485 | 137047 | 839.553 |
| 39 | H | 14.029 | 305683 | 12.577 |
| 44 | A | 15.442 | 395280 | 15.406 |
| 46 | R | 15.947 | 567572 | 26.274 |
| 48 | Y | 16.742 | 385043 | 12.088 |
| 56 | DTh | 18.408 | 469205 | 31.750 |
| 60 | V | 19.698 | 1694804 | 54.360 |
| 64 | W | 20.740 | 1799870 | 32.674 |
| 66 | K | 21.483 | 3038671 | 65.625 |
| 67 | F | 21.865 | 2928265 | 83.113 |
| 69 | I | 22.418 | 2444135 | 68.560 |
| 70 | L | 22.781 | 2390372 | 68.692 |
| 80 | DPU | 24.722 | 489488 | 366.313 |
| Total | | | 17782060 | |

| | |
|---|---|
| Data File | : 10022_08-07-2022_D03.lcd |
| Sample Name | : Sumit Mukherjee, 2 |
| Method File | : 10022_08-07-2022.lcm |
| Background Data File | : 10022_08-07-2022 D02.lcd |

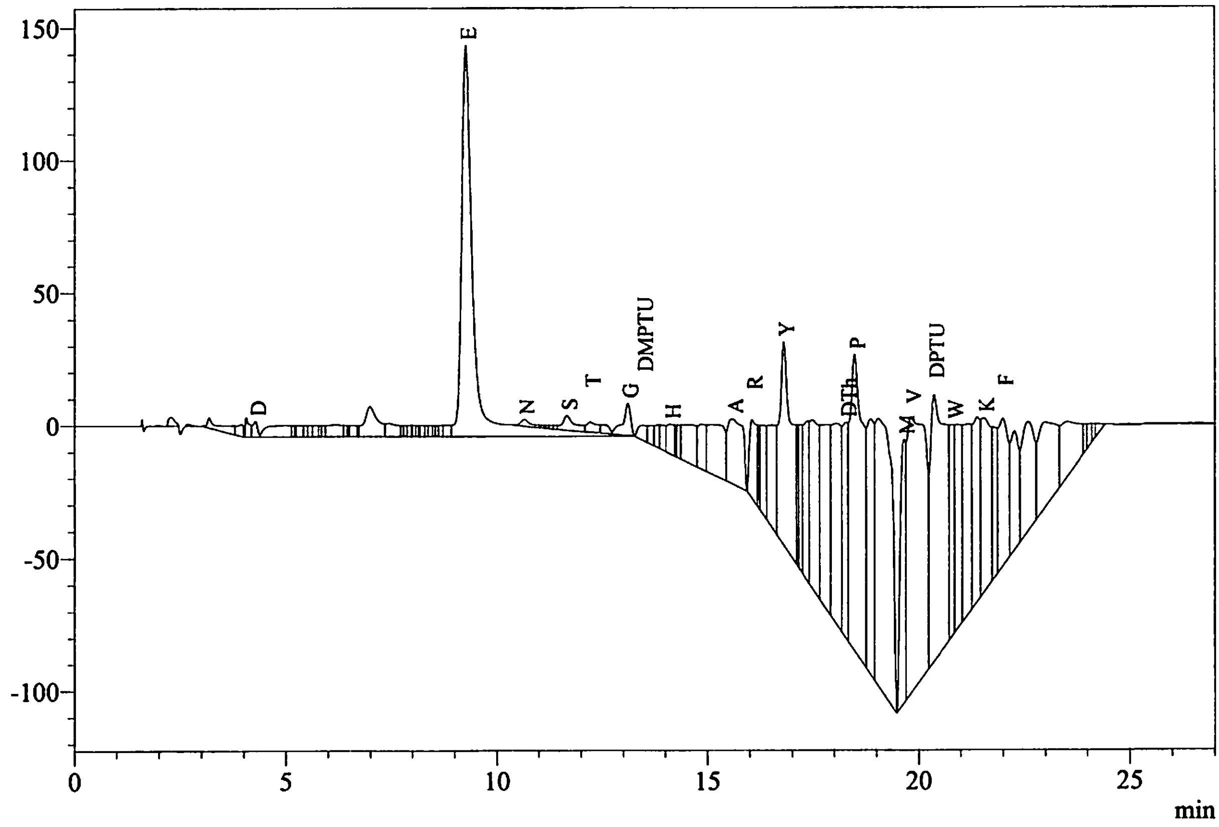


Peak Table
PDA Ch1 269nm

| Peak# | Name | Ret. Time | Area | Conc. |
|---|---|---|---|---|
| 8 | D | 4.284 | 47911 | 1.625 |
| 41 | E | 9.260 | 2983955 | 123.627 |
| 42 | N | 10.643 | 47012 | 2.075 |
| 49 | S | 11.659 | 116919 | 6.647 |
| 50 | T | 12.214 | 69625 | 3.429 |
| 52 | G | 13.099 | 156050 | 7.128 |
| 53 | DMPTU | 13.455 | 81839 | 501.348 |
| 57 | H | 14.110 | 140747 | 5.791 |
| 63 | A | 15.588 | 625413 | 24.375 |
| 64 | R | 16.051 | 364890 | 16.891 |
| 68 | Y | 16.801 | 1538837 | 48.310 |
| 75 | DTh | 18.286 | 718194 | 48.598 |
| 76 | P | 18.484 | 2404216 | 78.656 |
| 79 | M | 19.654 | 894345 | 27.279 |
| 80 | V | 19.822 | 3102215 | 99.502 |
| 81 | DPTU | 20.362 | 2549855 | |
| 82 | W | 20.818 | 637486 | 11.573 |
| 86 | K | 21.547 | 1025685 | 22.151 |
| 88 | F | 21.998 | 892988 | 25.346 |
| Total | | | 18398182 | |

| | |
|---|---|
| Data File | : 10022_08-07-2022_D04.lcd |
| Sample Name | : Sumit Mukherjee, 2 |
| Method File | : 10022_08-07-2022.lcm |
| Background Data File | : 10022_08-07-2022_D03.lcd |

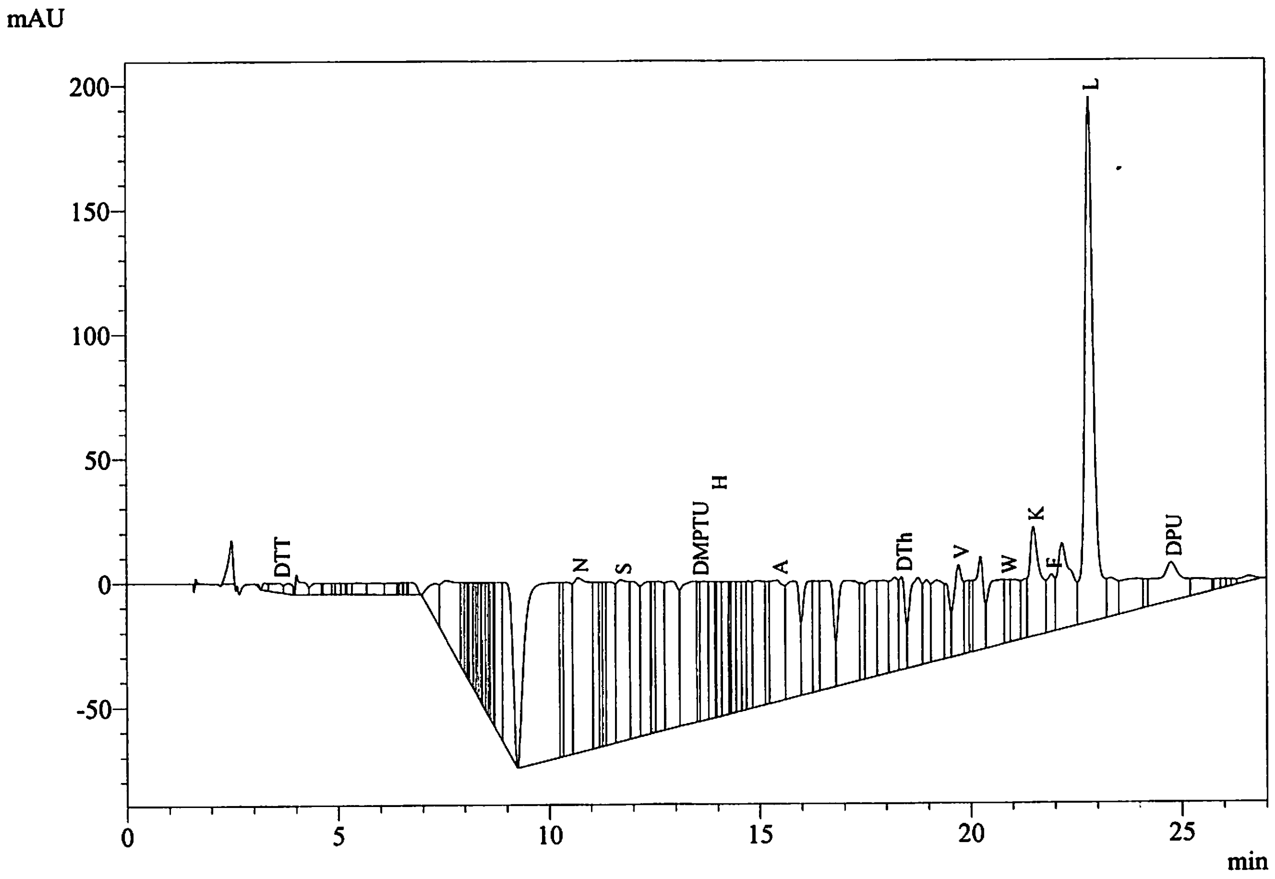


Peak Table
PDA Ch1 269nm

| Peak# | Name | Ret. Time | Area | Conc. |
|---|---|---|---|---|
| 5 | DTT | 3.608 | 72781 | 10.477 |
| 45 | N | 10.687 | 2039512 | 90.037 |
| 50 | S | 11.701 | 1315133 | 74.767 |
| 58 | DMPTU | 13.536 | 234324 | 1435.479 |
| 62 | H | 13.989 | 381983 | 15.716 |
| 71 | A | 15.424 | 1091083 | 42.524 |
| 81 | DTh | 18.378 | 367885 | 24.894 |
| 86 | V | 19.720 | 535926 | 17.189 |
| 91 | W | 20.862 | 227542 | 4.131 |
| 94 | K | 21.503 | 813292 | 17.564 |
| 95 | F | 21.927 | 288596 | 8.191 |
| 97 | L | 22.827 | 3064113 | 88.053 |
| 101 | DPU | 24.760 | 649633 | 486.160 |
| Total | | | 11081804 | |
