## Additional supplementary file 2 for "Activation of the *Plasmodium* egress effector subtilisin-like protease 1 is achieved by plasmepsin X destruction of the propiece"

### Iowa State University
### Protein Facility

#### PROTEIN/PEPTIDE SEQUENCE REPORT

**Date:** August 8, 2022

**To:** Sumit Mukherjee

**Sample Number:** 10021

**Sample Name:** 1

**Sample Preparation:** The membrane was washed with DI water and loaded onto the instrument for sequence analysis.

**Instrument:** Shimadzu PPSQ-53A

**Sequencing Method:** Edman Degradation

| Cycle Number | Amino Acid |
|---|---|
| 1 | L |
| 2 | E |
| 3 | K |
| 4 | V |

The major amino acid is listed first for each cycle. If you have any questions, feel free to contact me.

[Sequence Analysis]

| | |
|---|---|
| Data Acquired | : 8/7/2022 7:44:11 AM |
| Data Processed | : 8/8/2022 7:12:16 AM |
| Reactor | : 1 |
| Number of Cycles | : 5 |
| Sequence Schedule | : C:\PPSQ\SeqProg3_PDA_BGE\PVDF9-3G.sch |
| Sample Name | : Sumit Mukherjee, 1 |
| Sample Amount(pmol) | : 10.0 |
| Sample ID | : 10021 |
| Operator Name | : System Administrator |
| Data File | : 10021_08-07-2022 |
| Start Number | : 1 |
| Method File | : 10021_08-07-2022.lcm |
| Batch File | : 10021_08-07-2022.lcb |
| Data Folder Path | : C:\LabSolutions\Data\Project1\PPSQ\10021_08-07-2022 |
| Number of Analyses | : 5 / 5 |
| Standard File | : C:\LabSolutions\Data\Project1\PPSQ\10021_08-07-2022\PTH-AA_08-04-2022_D01.lcd |
| Data Comment | |

[Sequence]

L E K V

[Estimated Sequence]

| | 1 | 2 | 3 | 4 |
|---|---|---|---|---|
| 1st | L | E | K | V |
| 2nd | V | Q | G | N |
| 3rd | M | D | N | D |
| 4th | F | K | D | S |
| Reliability(%) | 50.4 | 100.0 | 34.2 | 100.0 |

[Evaluated Value]

| | 1 | 2 | 3 | 4 |
|---|---|---|---|---|
| D | 0.52 | 111.78 | 74.32 | 51.71 |
| E | 0.04 | 2354.96 | 0.44 | 0.41 |
| N | 0.63 | 48.08 | 76.71 | 96.84 |
| S | 26.93 | 0.74 | 70.38 | 31.26 |
| T | 0.85 | 12.35 | 44.72 | 0.76 |
| Q | 0.26 | 120.93 | 0.74 | 1.78 |
| G | 0.83 | 69.72 | 464.85 | 0.60 |
| H | 3.12 | 0.61 | 0.09 | 1.05 |
| A | 59.62 | 0.78 | 0.93 | 4.59 |
| R | 0.55 | 53.49 | 0.72 | 0.69 |
| Y | 0.91 | 16.46 | 9.76 | 0.79 |
| P | 0.93 | 6.50 | 8.39 | 0.92 |
| M | 240.18 | 0.21 | 0.72 | 6.82 |
| V | 528.61 | 0.49 | 0.94 | 4108.36 |

| | | | | |
|---|---|---|---|---|
| W | 0.59 | 2.18 | 3.14 | 0.80 |
| K | 0.81 | 78.69 | 1492.02 | 0.49 |
| F | 152.99 | 0.38 | 0.46 | 6.23 |
| I | 114.81 | 0.48 | 0.54 | 1.00 |
| L | 2695.89 | 0.31 | 0.23 | 0.94 |

[Amount Yield(pmol)]

| | 1 | 2 | 3 | 4 |
|---|---|---|---|---|
| D | 1.15 | 0.00 | 0.94 | 0.00 |
| E | 2.21 | 189.10 | 0.00 | 0.00 |
| N | 1.41 | 1.22 | 64.57 | 39.69 |
| S | 3.09 | 0.00 | 68.81 | 29.01 |
| T | 1.41 | 1.04 | 43.51 | 19.62 |
| Q | 0.79 | 2.30 | 0.00 | 20.67 |
| G | 5.25 | 1.62 | 73.55 | 26.26 |
| H | 0.00 | 0.00 | 0.00 | 29.43 |
| A | 4.19 | 0.00 | 43.45 | 28.43 |
| R | 2.79 | 1.55 | 29.18 | 50.19 |
| Y | 3.65 | 0.42 | 26.29 | 31.12 |
| P | 2.84 | 3.53 | 17.46 | 37.32 |
| M | 4.48 | 0.00 | 13.55 | 12.85 |
| V | 13.64 | 0.00 | 33.45 | 96.16 |
| W | 0.00 | 8.92 | 16.08 | 0.00 |
| K | 6.41 | 23.45 | 45.35 | 0.00 |
| F | 4.56 | 0.00 | 18.87 | 37.23 |
| I | 0.00 | 69.62 | 28.62 | 24.55 |
| L | 47.42 | 0.00 | 0.00 | 18.72 |

[Percent Yield]

| | |
|---|---|
| Amino Acid | : A,V,L |
| Initial Yield(%) | : 374.66 |
| Repetitive Yield(%) | : 126.57 |
| Correlation Coef. | : 1.000 |
| Number of Data | : 2 |

[Repetitive Yield(%)]

| | |
|---|---|
| Data File | : PTH-AA_08-04-2022_D01.lcd |
| Sample Name | : PTH-AA |
| Method File | : PTH-AA_08-04-2022.lcm |
| Background Data File | : |

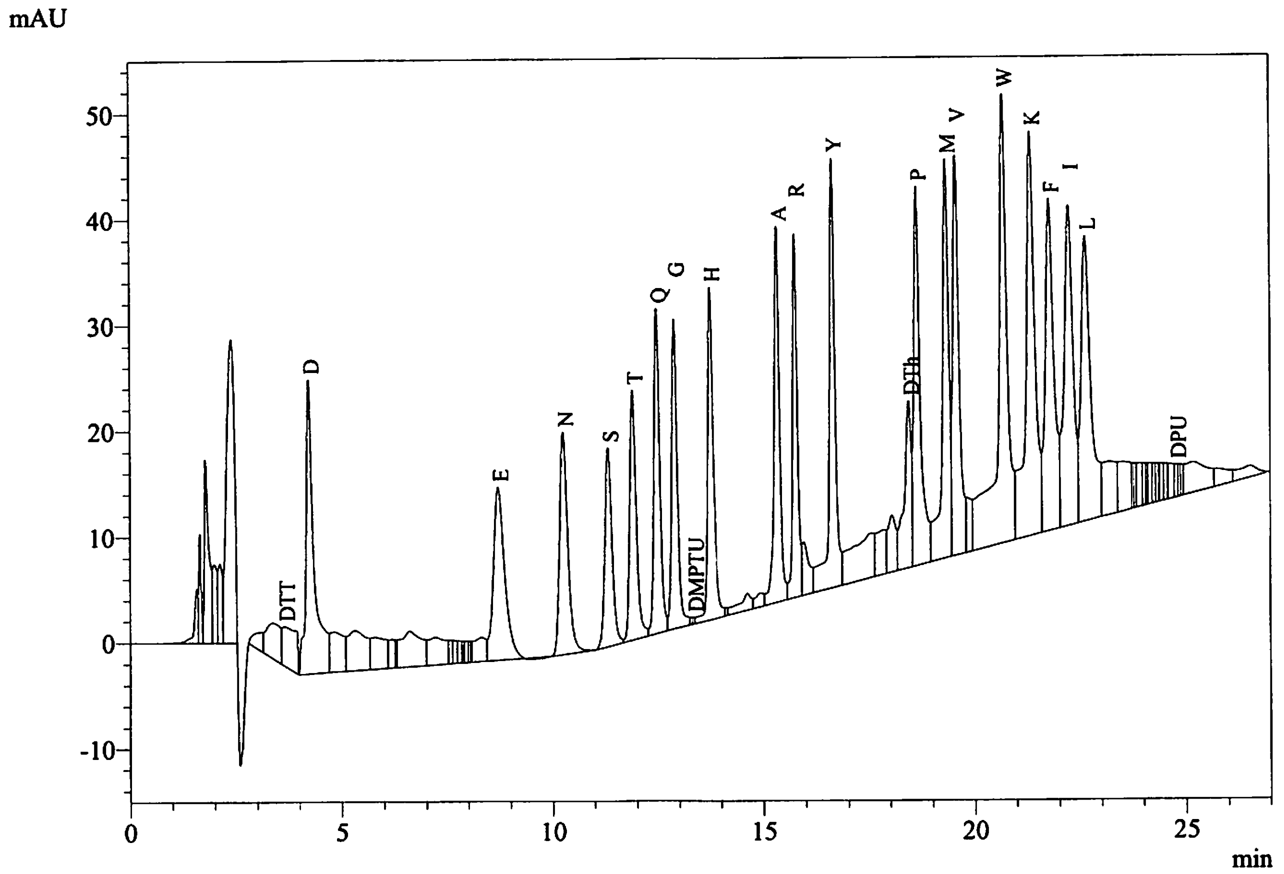


Peak Table
PDA Ch1 269nm

| Peak# | Name | Ret. Time | Area | Conc. |
|---|---|---|---|---|
| 9 | DTT | 3.639 | 86838 | 10.000 |
| 10 | D | 4.229 | 368538 | 10.000 |
| 26 | E | 8.709 | 301709 | 10.000 |
| 27 | N | 10.248 | 283149 | 10.000 |
| 28 | S | 11.321 | 219871 | 10.000 |
| 29 | T | 11.902 | 253806 | 10.000 |
| 30 | Q | 12.472 | 292815 | 10.000 |
| 31 | G | 12.889 | 273666 | 10.000 |
| 33 | DMPTU | 13.333 | 2040 | 10.000 |
| 34 | H | 13.739 | 303815 | 10.000 |
| 38 | A | 15.333 | 320725 | 10.000 |
| 39 | R | 15.757 | 270026 | 10.000 |
| 41 | Y | 16.643 | 398165 | 10.000 |
| 45 | DTh | 18.456 | 184729 | 10.000 |
| 46 | P | 18.650 | 382080 | 10.000 |
| 47 | M | 19.342 | 409815 | 10.000 |
| 48 | V | 19.578 | 389719 | 10.000 |
| 50 | W | 20.706 | 688571 | 10.000 |
| 51 | K | 21.353 | 578793 | 10.000 |
| 52 | F | 21.795 | 440406 | 10.000 |
| 53 | I | 22.255 | 445619 | 10.000 |
| 54 | L | 22.643 | 434982 | 10.000 |
| 69 | DPU | 24.779 | 16703 | 10.000 |
| Total | | | 7346580 | |

Data File : 10021_08-07-2022_D01.lcd
Sample Name : Sumit Mukherjee, 1
Method File : 10021_08-07-2022.lcm
Background Data File :

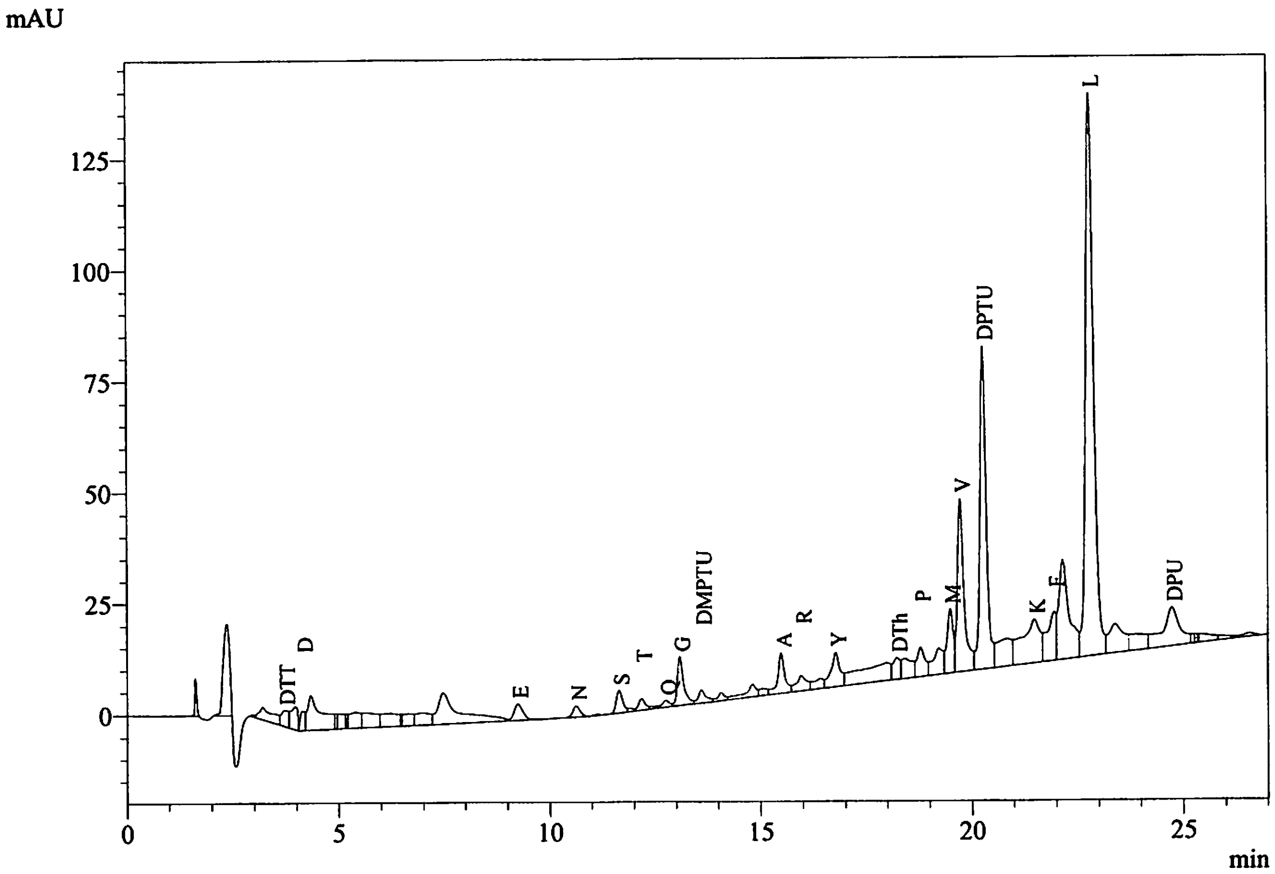


Peak Table
PDA Ch1 269nm

| Peak# | Name | Ret. Time | Area | Conc. |
|---|---|---|---|---|
| 5 | DTT | 3.718 | 42759 | 6.155 |
| 7 | D | 4.148 | 33902 | 1.150 |
| 19 | E | 9.237 | 53397 | 2.212 |
| 21 | N | 10.620 | 32005 | 1.413 |
| 23 | S | 11.638 | 54292 | 3.087 |
| 25 | T | 12.171 | 28601 | 1.409 |
| 28 | Q | 12.745 | 18609 | 0.794 |
| 29 | G | 13.077 | 114938 | 5.250 |
| 30 | DMPTU | 13.585 | 37846 | 231.846 |
| 34 | A | 15.482 | 107625 | 4.195 |
| 35 | R | 15.959 | 60355 | 2.794 |
| 37 | Y | 16.775 | 116167 | 3.647 |
| 39 | DTh | 18.231 | 57753 | 3.908 |
| 41 | P | 18.789 | 86806 | 2.840 |
| 43 | M | 19.490 | 146771 | 4.477 |
| 44 | V | 19.731 | 425211 | 13.638 |
| 45 | DPTU | 20.274 | 754594 | |
| 47 | K | 21.486 | 296851 | 6.411 |
| 48 | F | 21.955 | 160536 | 4.556 |
| 50 | L | 22.810 | 1650239 | 47.423 |
| 53 | DPU | 24.738 | 255113 | 190.916 |
| Total | | | 4534369 | |

Data File : 10021_08-07-2022_D02.lcd
Sample Name : Sumit Mukherjee, 1
Method File : 10021_08-07-2022.lcm
Background Data File : 10021_08-07-2022_D01.lcd

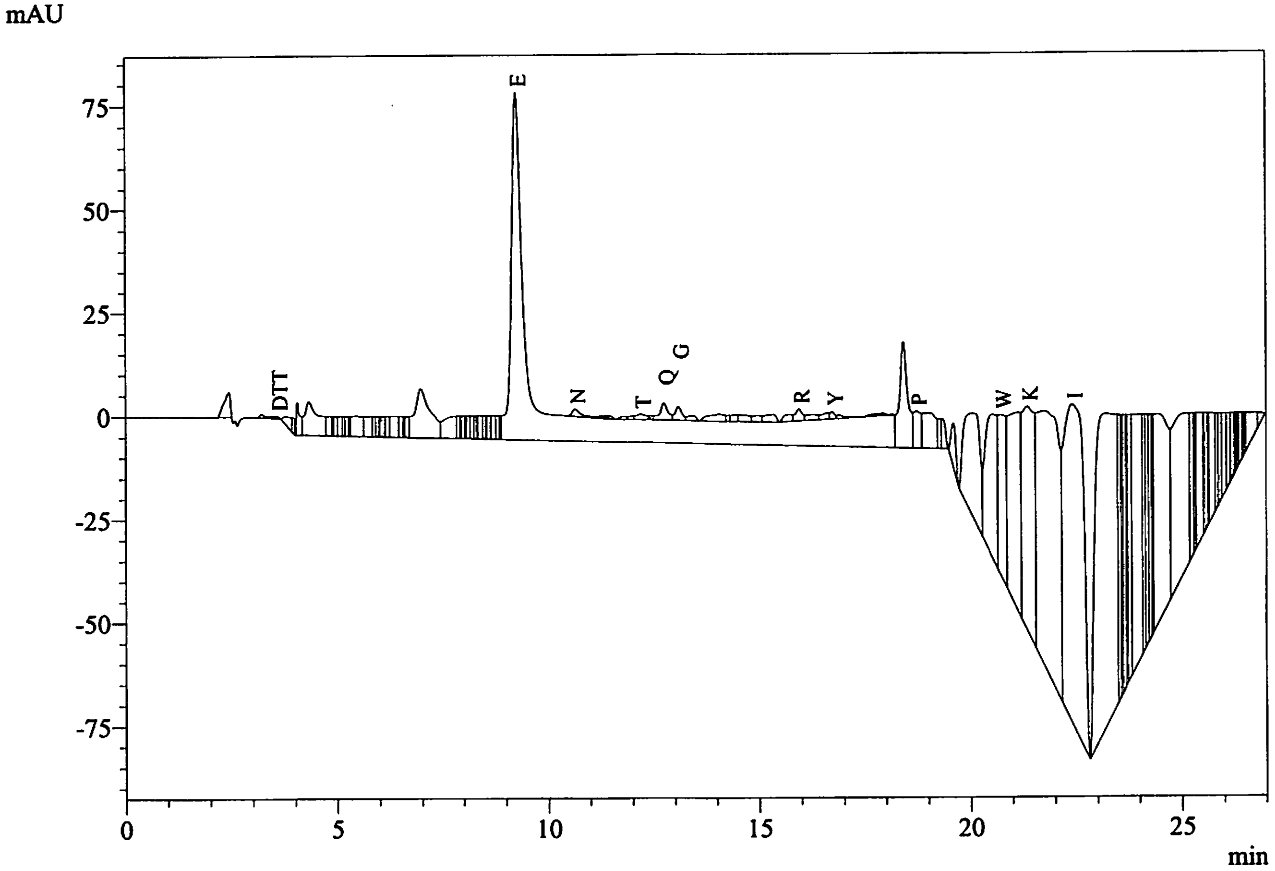


Peak Table
PDA Ch1 269nm

| Peak# | Name | Ret. Time | Area | Conc. |
|---|---|---|---|---|
| 5 | DTT | 3.579 | 2311 | 0.333 |
| 46 | E | 9.246 | 4564192 | 189.097 |
| 47 | N | 10.640 | 27685 | 1.222 |
| 55 | T | 12.188 | 21069 | 1.038 |
| 58 | Q | 12.746 | 53922 | 2.302 |
| 59 | G | 13.086 | 35387 | 1.616 |
| 68 | R | 15.947 | 33483 | 1.550 |
| 71 | Y | 16.735 | 13342 | 0.419 |
| 77 | P | 18.740 | 107918 | 3.531 |
| 84 | W | 20.744 | 491596 | 8.924 |
| 86 | K | 21.366 | 1085618 | 23.446 |
| 88 | I | 22.422 | 2481924 | 69.620 |
| Total | | | 8918445 | |

| | |
|---|---|
| Data File | : 10021_08-07-2022_D03.lcd |
| Sample Name | : Sumit Mukherjee, 1 |
| Method File | : 10021_08-07-2022.lcm |
| Background Data File | : 10021_08-07-2022_D02.lcd |

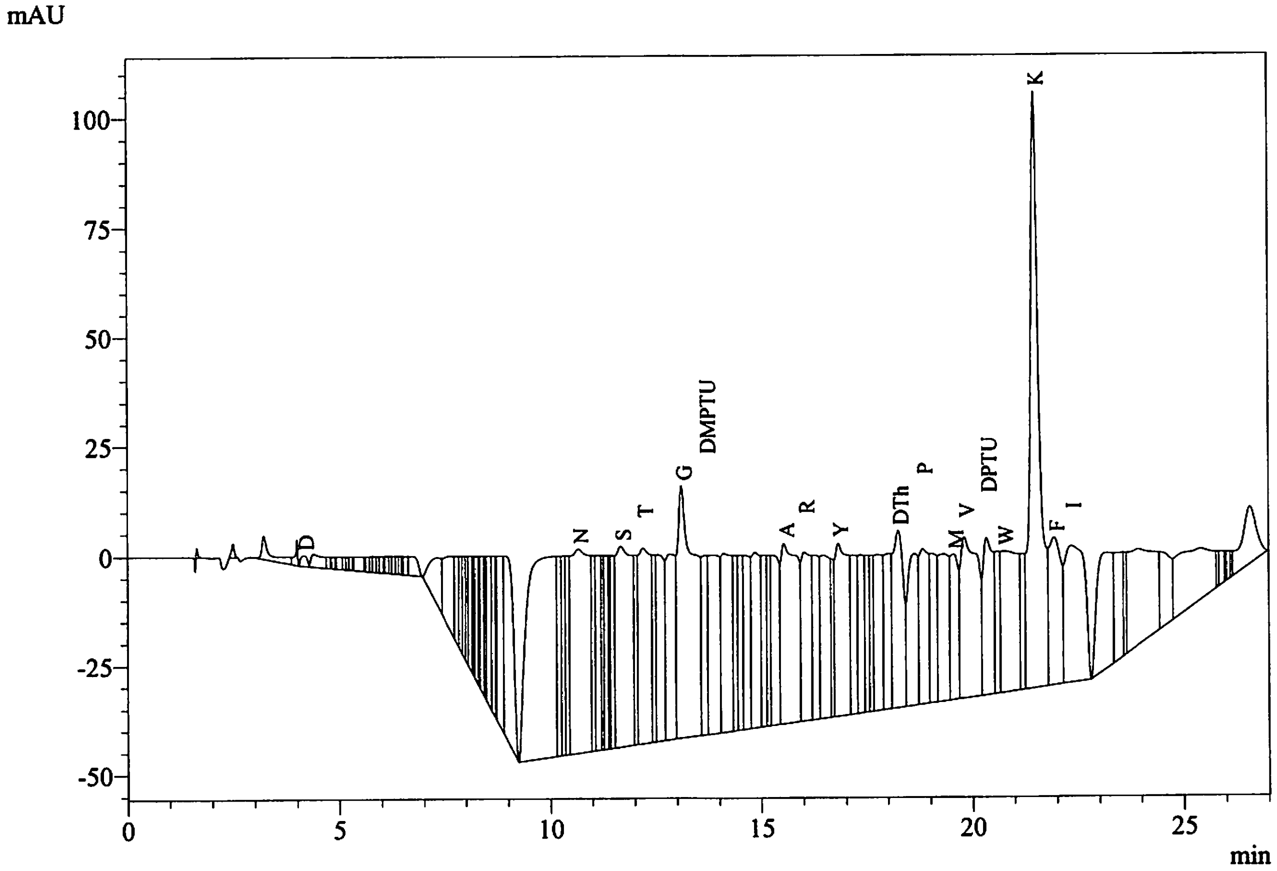


Peak Table
PDA Ch1 269nm

| Peak# | Name | Ret. Time | Area | Conc. |
|---|---|---|---|---|
| 5 | D | 4.160 | 27667 | 0.938 |
| 55 | N | 10.664 | 1462684 | 64.572 |
| 62 | S | 11.674 | 1210391 | 68.813 |
| 64 | T | 12.202 | 883439 | 43.510 |
| 68 | G | 13.108 | 1610261 | 73.550 |
| 69 | DMPTU | 13.673 | 394203 | 2414.899 |
| 79 | A | 15.541 | 1114876 | 43.451 |
| 80 | R | 16.023 | 630317 | 29.179 |
| 84 | Y | 16.825 | 837317 | 26.287 |
| 91 | DTh | 18.256 | 697935 | 47.227 |
| 93 | P | 18.830 | 533648 | 17.459 |
| 96 | M | 19.556 | 444101 | 13.546 |
| 97 | V | 19.812 | 1042949 | 33.452 |
| 98 | DPTU | 20.338 | 599996 | |
| 100 | W | 20.746 | 885905 | 16.082 |
| 102 | K | 21.487 | 2099747 | 45.348 |
| 103 | F | 21.947 | 664758 | 18.868 |
| 104 | I | 22.345 | 1020403 | 28.623 |
| Total | | | 16160598 | |

| | |
|---|---|
| Data File | : 10021_08-07-2022_D04.lcd |
| Sample Name | : Sumit Mukherjee, 1 |
| Method File | : 10021_08-07-2022.lcm |
| Background Data File | : 10021_08-07-2022_D03.lcd |

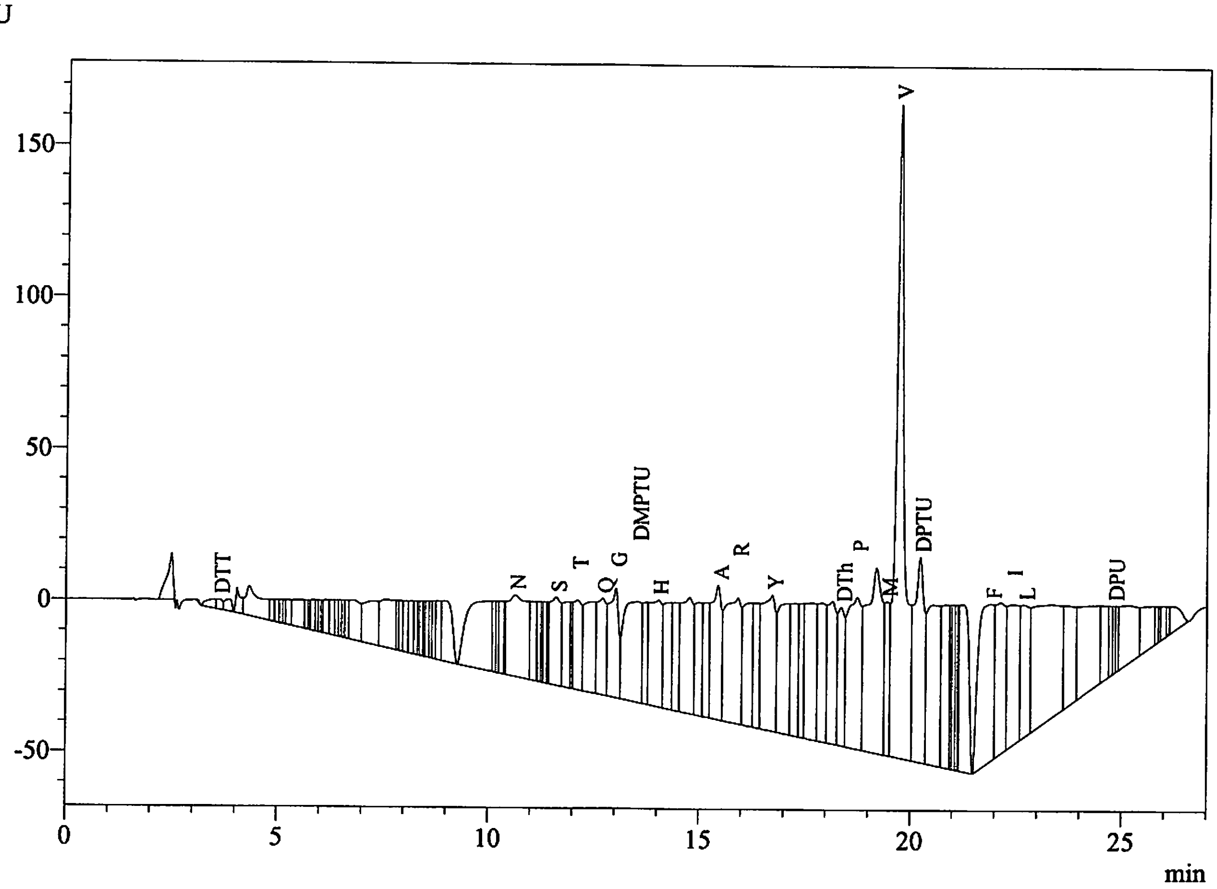


Peak Table
PDA Ch1 269nm

| Peak# | Name | Ret. Time | Area | Conc. |
|---|---|---|---|---|
| 4 | DTT | 3.617 | 31793 | 4.576 |
| 49 | N | 10.622 | 898963 | 39.686 |
| 56 | S | 11.601 | 510350 | 29.014 |
| 59 | T | 12.108 | 398348 | 19.619 |
| 61 | Q | 12.700 | 484267 | 20.673 |
| 62 | G | 13.016 | 574960 | 26.262 |
| 63 | DMPTU | 13.524 | 959796 | 5879.745 |
| 65 | H | 14.022 | 715344 | 29.432 |
| 71 | A | 15.442 | 729541 | 28.433 |
| 72 | R | 15.912 | 1084148 | 50.187 |
| 75 | Y | 16.724 | 991315 | 31.121 |
| 82 | DTh | 18.364 | 513028 | 34.715 |
| 83 | P | 18.742 | 1140819 | 37.323 |
| 85 | M | 19.445 | 421416 | 12.854 |
| 86 | V | 19.717 | 2998176 | 96.165 |
| 87 | DPTU | 20.225 | 1128736 | |
| 95 | F | 21.907 | 1311731 | 37.231 |
| 97 | I | 22.406 | 875274 | 24.552 |
| 98 | L | 22.699 | 651539 | 18.723 |
| 104 | DPU | 24.811 | 100028 | 74.857 |
| Total | | | 16519570 | |
