## Additional supplementary file 3 for "Activation of the *Plasmodium* egress effector subtilisin-like protease 1 is achieved by plasmepsin X destruction of the propiece"

[Sequence Analysis]

| | |
|---|---|
| Data Acquired | : 8/7/2022 5:19:52 PM |
| Data Processed | : 8/8/2022 7:31:33 AM |
| Reactor | : 3 |
| Number of Cycles | : 5 |
| Sequence Schedule | : C:\PPSQ\SeqProg3_PDA_BGE\PVDF9-3G.sch |
| Sample Name | : Sumit Mukherjee, 3 |
| Sample Amount(pmol) | : 10.0 |
| Sample ID | : 10023 |
| Operator Name | : System Administrator |
| Data File | : 10023_08-07-2022 |
| Start Number | : 1 |
| Method File | : 10023_08-07-2022.lcm |
| Batch File | : 10023_08-07-2022.lcb |
| Data Folder Path | : C:\LabSolutions\Data\Project1\PPSQ\10023_08-07-2022 |
| Number of Analyses | : 5 / 5 |
| Standard File | : C:\LabSolutions\Data\Project1\PPSQ\10023_08-07-2022\PTH-AA_08-04-2022_D01.lcd |
| Data Comment | |

[Sequence]
L E S K

[Estimated Sequence]

| | 1 | 2 | 3 | 4 |
|---|---|---|---|---|
| 1st | L | E | S | K |
| 2nd | S | P | V | L |
| 3rd | G | R | N | F |
| 4th | D | K | W | V |
| Reliability(%) | 27.6 | 26.6 | 71.6 | 100.0 |

[Evaluated Value]

| | 1 | 2 | 3 | 4 |
|---|---|---|---|---|
| D | 837.44 | 0.43 | 0.57 | 0.87 |
| E | 0.13 | 3794.90 | 0.23 | 0.41 |
| N | 198.26 | 0.39 | 139.46 | 0.63 |
| S | 2084.85 | 0.12 | 1628.95 | 0.31 |
| T | 420.58 | 0.23 | 0.99 | 23.43 |
| Q | 102.93 | 0.52 | 0.44 | 7.17 |
| G | 982.48 | 0.28 | 0.92 | 23.45 |
| H | 649.09 | 0.28 | 0.37 | 0.84 |
| A | 497.66 | 0.25 | 0.66 | 48.10 |
| R | 0.83 | 0.77 | 0.31 | 0.74 |
| Y | 392.68 | 0.37 | 0.77 | 0.90 |
| P | 0.01 | 1523.60 | 0.16 | 0.16 |
| M | 810.57 | 0.11 | 0.59 | 14.64 |
| V | 332.34 | 0.37 | 207.66 | 132.18 |

| | | | | |
|---|---|---|---|---|
| W | 114.62 | 0.15 | 60.90 | 0.79 |
| K | 17.20 | 0.72 | 0.79 | 3824.95 |
| F | 601.35 | 0.09 | 0.42 | 235.27 |
| I | 273.73 | 0.37 | 0.65 | 56.28 |
| L | 5939.41 | 0.09 | 0.31 | 340.43 |

[Amount Yield(pmol)]

| | 1 | 2 | 3 | 4 |
|---|---|---|---|---|
| D | 24.49 | 0.00 | 0.00 | 0.00 |
| E | 12.11 | 249.76 | 0.00 | 0.00 |
| N | 5.94 | 0.00 | 505.66 | 0.00 |
| S | 42.78 | 0.00 | 306.08 | 0.00 |
| T | 10.31 | 0.00 | 80.89 | 38.21 |
| Q | 3.87 | 0.00 | 85.74 | 31.31 |
| G | 22.36 | 0.00 | 0.00 | 35.43 |
| H | 13.28 | 98.77 | 80.07 | 25.08 |
| A | 10.45 | 0.00 | 71.59 | 35.85 |
| R | 8.36 | 94.20 | 0.00 | 21.82 |
| Y | 8.57 | 0.00 | 30.96 | 16.59 |
| P | 3.17 | 176.70 | 0.00 | 0.00 |
| M | 14.18 | 0.00 | 0.00 | 9.59 |
| V | 10.67 | 0.00 | 72.85 | 0.00 |
| W | 5.55 | 0.00 | 71.81 | 3.54 |
| K | 5.89 | 154.00 | 0.00 | 36.12 |
| F | 10.81 | 60.41 | 23.97 | 5.14 |
| I | 0.00 | 184.12 | 27.84 | 5.96 |
| L | 83.67 | 0.00 | 0.00 | 13.23 |

[Percent Yield]

| | | |
|---|---|---|
| Amino Acid | : | A,V,L |
| Initial Yield(%) | : | 836.69 |
| Repetitive Yield(%) | : | 100.00 |
| Correlation Coef. | : | 1.000 |
| Number of Data | : | 1 |

[Repetitive Yield(%)]

| | |
|---|---|
| Data File | : PTH-AA_08-04-2022_D01.lcd |
| Sample Name | : PTH-AA |
| Method File | : PTH-AA_08-04-2022.lcm |
| Background Data File | : |

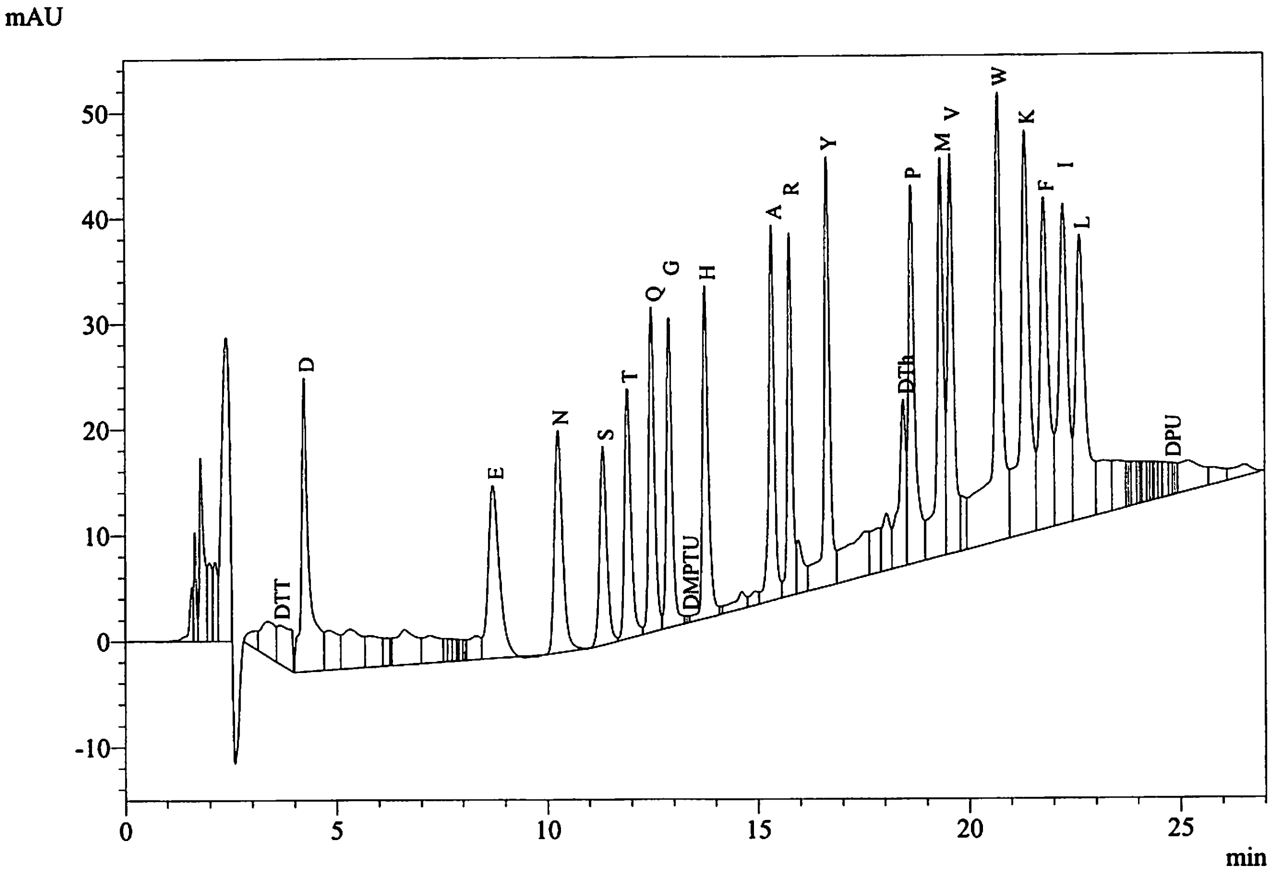


Peak Table
PDA Ch1 269nm

| Peak# | Name | Ret. Time | Area | Conc. |
|---|---|---|---|---|
| 9 | DTT | 3.639 | 86838 | 10.000 |
| 10 | D | 4.229 | 368538 | 10.000 |
| 26 | E | 8.709 | 301709 | 10.000 |
| 27 | N | 10.248 | 283149 | 10.000 |
| 28 | S | 11.321 | 219871 | 10.000 |
| 29 | T | 11.902 | 253806 | 10.000 |
| 30 | Q | 12.472 | 292815 | 10.000 |
| 31 | G | 12.889 | 273666 | 10.000 |
| 33 | DMPTU | 13.333 | 2040 | 10.000 |
| 34 | H | 13.739 | 303815 | 10.000 |
| 38 | A | 15.333 | 320725 | 10.000 |
| 39 | R | 15.757 | 270026 | 10.000 |
| 41 | Y | 16.643 | 398165 | 10.000 |
| 45 | DTh | 18.456 | 184729 | 10.000 |
| 46 | P | 18.650 | 382080 | 10.000 |
| 47 | M | 19.342 | 409815 | 10.000 |
| 48 | V | 19.578 | 389719 | 10.000 |
| 50 | W | 20.706 | 688571 | 10.000 |
| 51 | K | 21.353 | 578793 | 10.000 |
| 52 | F | 21.795 | 440406 | 10.000 |
| 53 | I | 22.255 | 445619 | 10.000 |
| 54 | L | 22.643 | 434982 | 10.000 |
| 69 | DPU | 24.779 | 16703 | 10.000 |
| Total | | | 7346580 | |

| | |
|---|---|
| Data File | : 10023_08-07-2022_D01.lcd |
| Sample Name | : Sumit Mukherjee, 3 |
| Method File | : 10023_08-07-2022.lcm |
| Background Data File | : |

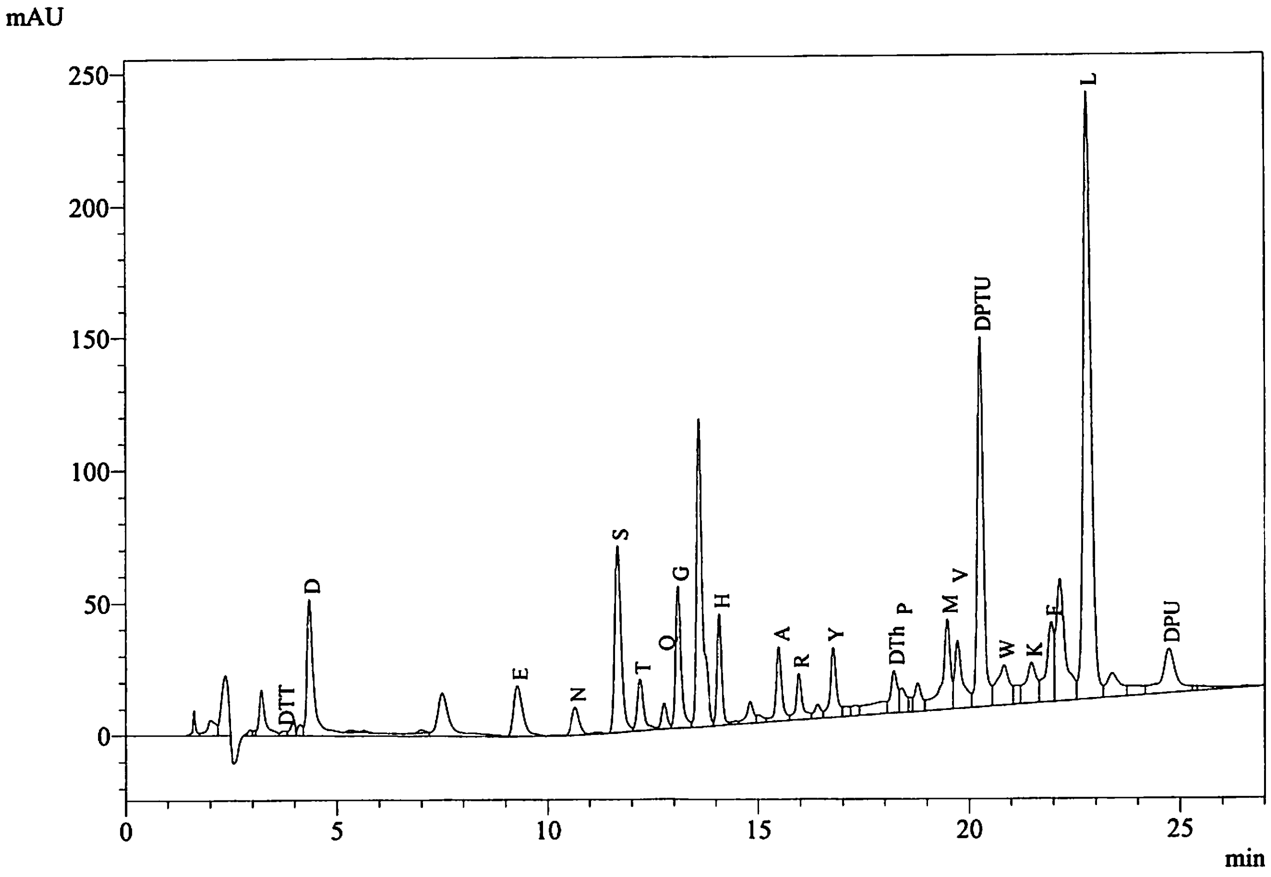


Peak Table
PDA Ch1 269nm

| Peak# | Name | Ret. Time | Area | Conc. |
|---|---|---|---|---|
| 7 | DTT | 3.732 | 18348 | 2.641 |
| 10 | D | 4.347 | 722107 | 24.492 |
| 15 | E | 9.273 | 292360 | 12.113 |
| 16 | N | 10.647 | 134573 | 5.941 |
| 18 | S | 11.659 | 752419 | 42.776 |
| 19 | T | 12.185 | 209336 | 10.310 |
| 20 | Q | 12.756 | 90735 | 3.873 |
| 21 | G | 13.086 | 489593 | 22.363 |
| 23 | H | 14.072 | 322846 | 13.283 |
| 27 | A | 15.481 | 268019 | 10.446 |
| 28 | R | 15.954 | 180593 | 8.360 |
| 30 | Y | 16.772 | 272914 | 8.568 |
| 34 | DTh | 18.218 | 179248 | 12.129 |
| 35 | P | 18.405 | 97031 | 3.174 |
| 38 | M | 19.479 | 464923 | 14.181 |
| 39 | V | 19.723 | 332611 | 10.668 |
| 40 | DPTU | 20.264 | 1388645 | |
| 41 | W | 20.833 | 305818 | 5.552 |
| 43 | K | 21.480 | 272746 | 5.890 |
| 44 | F | 21.946 | 380823 | 10.809 |
| 46 | L | 22.804 | 2911544 | 83.669 |
| 49 | DPU | 24.736 | 398288 | 298.063 |
| Total | | | 10485522 | |

| | |
|---|---|
| Data File | : 10023_08-07-2022_D02.lcd |
| Sample Name | : Sumit Mukherjee, 3 |
| Method File | : 10023_08-07-2022.lcm |
| Background Data File | : 10023_08-07-2022_D01.lcd |

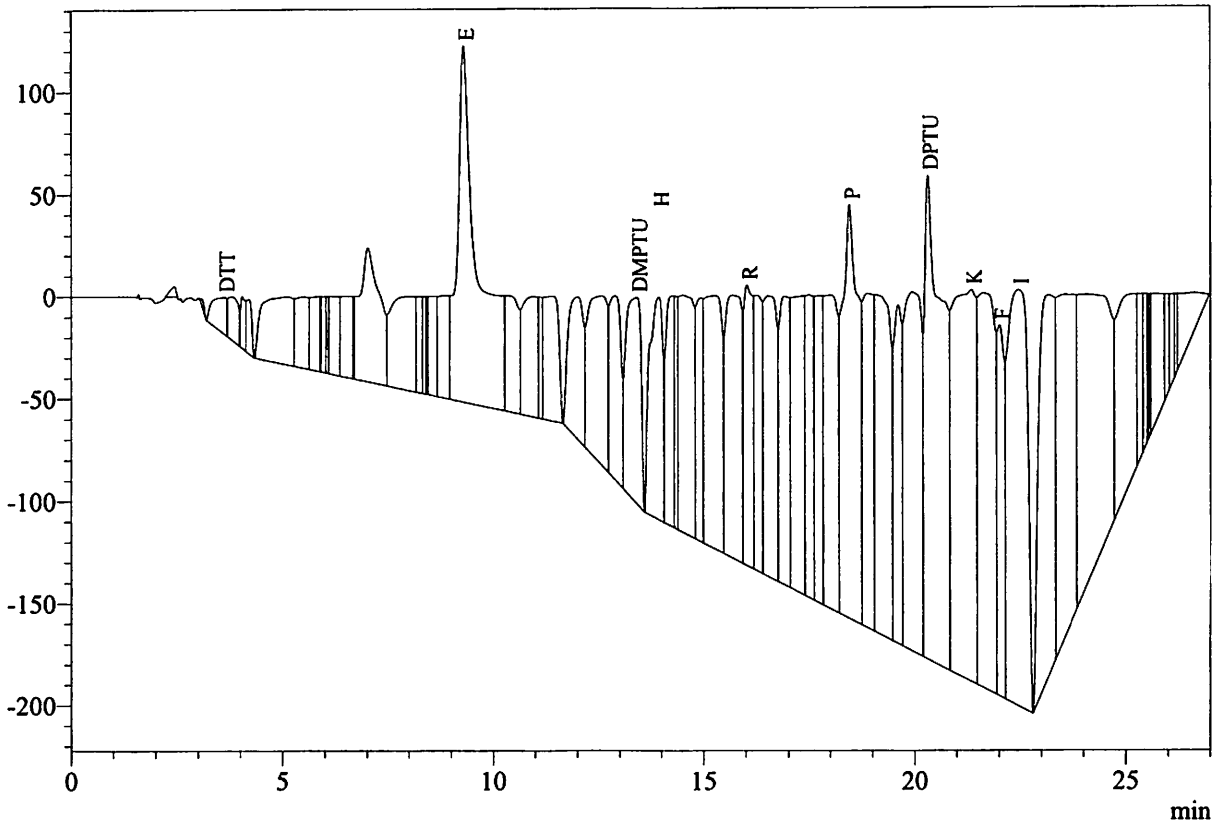


Peak Table
PDA Ch1 269nm

| Peak# | Name | Ret. Time | Area | Conc. |
|---|---|---|---|---|
| 4 | DTT | 3.615 | 385916 | 55.551 |
| 24 | E | 9.297 | 6028348 | 249.758 |
| 32 | DMPTU | 13.405 | 2427254 | 14869.448 |
| 33 | H | 13.928 | 2400610 | 98.769 |
| 40 | R | 16.025 | 2034845 | 94.197 |
| 48 | P | 18.461 | 5401151 | 176.702 |
| 53 | DPTU | 20.316 | 7228865 | |
| 54 | K | 21.353 | 7130963 | 154.005 |
| 56 | F | 22.020 | 2128261 | 60.406 |
| 57 | I | 22.463 | 6563814 | 184.121 |
| Total | | | 41730027 | |

| | |
|---|---|
| Data File | : 10023_08-07-2022_D03.lcd |
| Sample Name | : Sumit Mukherjee, 3 |
| Method File | : 10023_08-07-2022.lcm |
| Background Data File | : 10023_08-07-2022_D02.lcd |

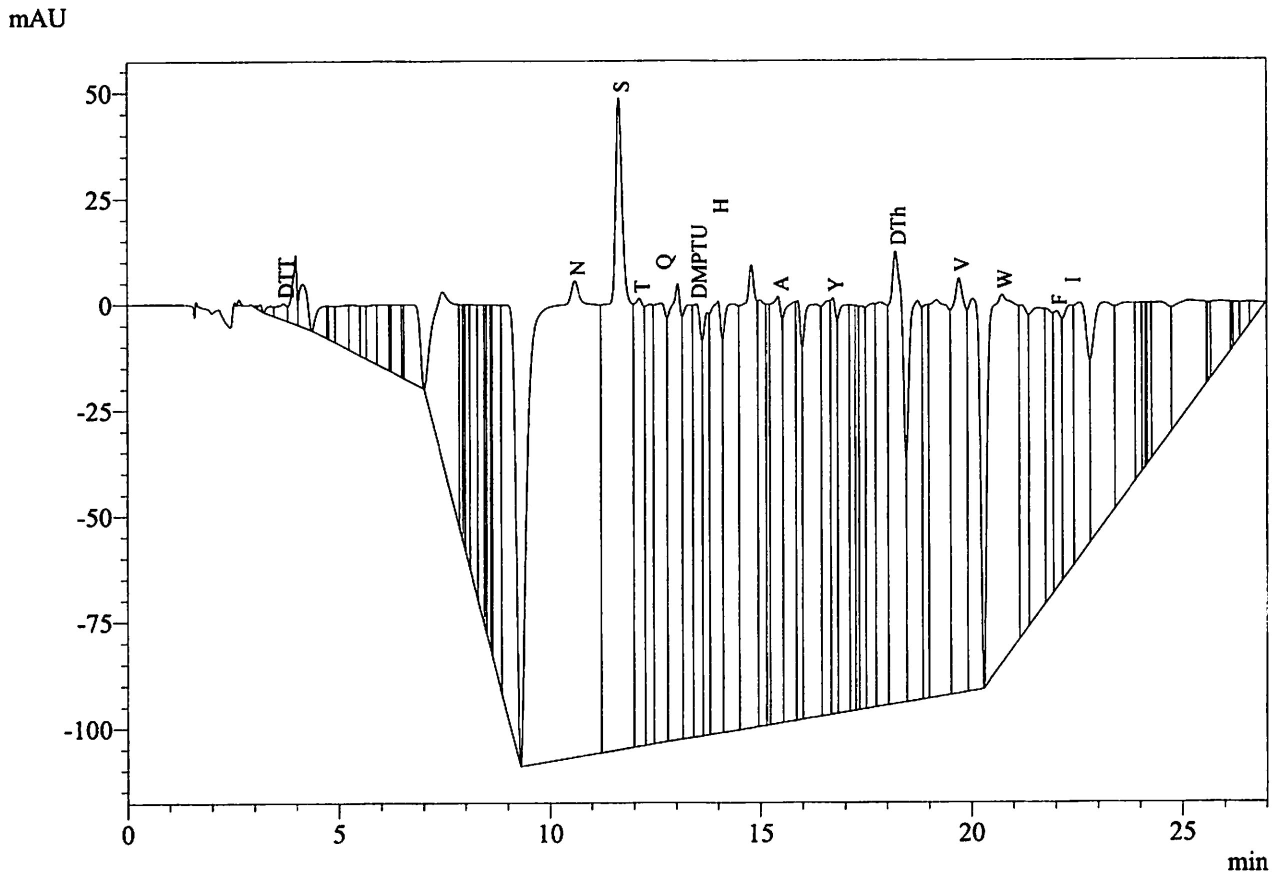


Peak Table
PDA Ch1 269nm

| Peak# | Name | Ret. Time | Area | Conc. |
|---|---|---|---|---|
| 7 | DTT | 3.692 | 62288 | 8.966 |
| 33 | N | 10.603 | 11454258 | 505.664 |
| 34 | S | 11.654 | 5383928 | 306.084 |
| 35 | T | 12.138 | 1642415 | 80.889 |
| 37 | Q | 12.659 | 2008472 | 85.740 |
| 40 | DMPTU | 13.491 | 1371945 | 8404.586 |
| 42 | H | 14.011 | 1946047 | 80.067 |
| 47 | A | 15.434 | 1836814 | 71.588 |
| 52 | Y | 16.732 | 986138 | 30.959 |
| 59 | DTh | 18.221 | 2425043 | 164.095 |
| 63 | V | 19.722 | 2271217 | 72.848 |
| 65 | W | 20.753 | 3955496 | 71.806 |
| 69 | F | 22.041 | 844417 | 23.967 |
| 70 | I | 22.356 | 992345 | 27.836 |
| Total | | | 37180820 | |

| | |
|---|---|
| Data File | : 10023_08-07-2022_D04.lcd |
| Sample Name | : Sumit Mukherjee, 3 |
| Method File | : 10023_08-07-2022.lcm |
| Background Data File | : 10023_08-07-2022_D03.lcd |

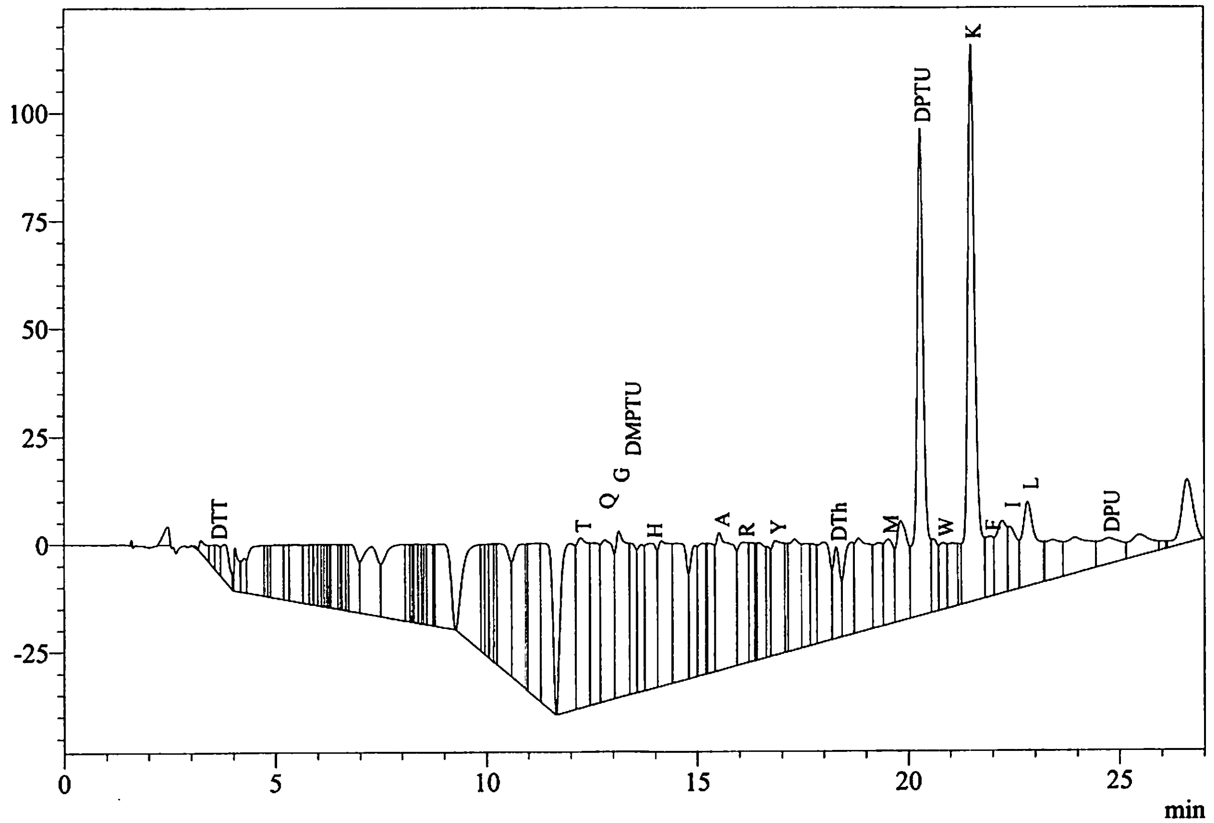


Peak Table
PDA Ch1 269nm

| Peak# | Name | Ret. Time | Area | Conc. |
|---|---|---|---|---|
| 6 | DTT | 3.599 | 50161 | 7.221 |
| 53 | T | 12.232 | 775846 | 38.211 |
| 55 | Q | 12.805 | 733546 | 31.314 |
| 56 | G | 13.137 | 775681 | 35.430 |
| 57 | DMPTU | 13.416 | 363001 | 2223.760 |
| 59 | H | 13.920 | 609648 | 25.083 |
| 66 | A | 15.523 | 919927 | 35.853 |
| 67 | R | 16.102 | 471411 | 21.822 |
| 72 | Y | 16.853 | 528602 | 16.595 |
| 78 | DTh | 18.301 | 245635 | 16.621 |
| 82 | M | 19.525 | 314338 | 9.588 |
| 84 | DPTU | 20.292 | 1373452 | |
| 86 | W | 20.835 | 194822 | 3.537 |
| 89 | K | 21.505 | 1672692 | 36.125 |
| 90 | F | 21.945 | 181224 | 5.144 |
| 92 | I | 22.413 | 212437 | 5.959 |
| 93 | L | 22.832 | 460558 | 13.235 |
| 97 | DPU | 24.755 | 228104 | 170.704 |
| Total | | | 10111086 | |
