## Additional supplementary file 4 for "Activation of the *Plasmodium* egress effector subtilisin-like protease 1 is achieved by plasmepsin X destruction of the propiece"

### Iowa State University
### Protein Facility

#### PROTEIN/PEPTIDE SEQUENCE REPORT

**Date:** September 3, 2022

**To:** Sumit Mukherjee

**Sample Number:** 10048

**Sample Name:** Sample 1

**Sample Preparation:** The membrane was washed with DI water and loaded onto the instrument for sequence analysis.

**Instrument:** Shimadzu PPSQ-53A

**Sequencing Method:** Edman Degradation

| Cycle Number | Amino Acid |
|---|---|
| 1 | F |
| 2 | Q |
| 3 | E |
| 4 | S |

The major amino acid is listed first for each cycle. If you have any questions, feel free to contact me.

[Sequence Analysis]

| | | |
|---|---|---|
| Data Acquired | : | 9/2/2022 11:54:53 AM |
| Data Processed | : | 9/2/2022 5:03:18 PM |
| Reactor | : | 1 |
| Number of Cycles | : | 5 |
| Sequence Schedule | : | C:\PPSQ\SeqProg3_PDA_BGE\PVDF9-3G.sch |
| Sample Name | : | Sumit Mukherjee, Sample 1 |
| Sample Amount(pmol) | : | 10.0 |
| Sample ID | : | 10048 |
| Operator Name | : | System Administrator |
| Data File | : | 10048_09-02-2022 |
| Start Number | : | 1 |
| Method File | : | 10048_09-02-2022.lcm |
| Batch File | : | 10048_09-02-2022.lcb |
| Data Folder Path | : | C:\LabSolutions\Data\Project1\PPSQ\10048_09-02-2022 |
| Number of Analyses | : | 5 / 5 |
| Standard File | : | C:\LabSolutions\Data\Project1\PPSQ\10048_09-02-2022\PTH-AA_08-31-2022_D01.lcd |
| Data Comment | | |

[Sequence]

F Q E S

[Estimated Sequence]

| | 1 | 2 | 3 | 4 |
|---|---|---|---|---|
| 1st | F | Q | E | S |
| 2nd | M | Y | Y | A |
| 3rd | L | E | P | I |
| 4th | G | A | D | N |
| Reliability(%) | 100.0 | 71.3 | 49.2 | 100.0 |

[Evaluated Value]

| | 1 | 2 | 3 | 4 |
|---|---|---|---|---|
| D | 0.49 | 67.06 | 112.94 | 80.60 |
| E | 0.09 | 958.89 | 6047.47 | 0.16 |
| N | 0.72 | 35.44 | 97.61 | 85.88 |
| S | 0.13 | 0.46 | 0.29 | 2700.62 |
| T | 0.61 | 37.82 | 66.43 | 0.77 |
| Q | 0.00 | 12323.40 | 0.12 | 0.18 |
| G | 8.28 | 0.79 | 0.95 | 0.87 |
| H | 0.54 | 0.48 | 0.64 | 0.00 |
| A | 0.32 | 230.92 | 0.76 | 123.47 |
| R | 1.47 | 0.72 | 0.00 | 1.72 |
| Y | 0.17 | 1689.43 | 1346.08 | 0.23 |
| P | 0.47 | 81.09 | 127.78 | 0.59 |
| M | 112.95 | 0.35 | 47.74 | 23.10 |
| V | 0.51 | 97.90 | 77.34 | 0.66 |

| | | | | |
|---|---|---|---|---|
| W | 3.64 | 0.75 | 1.57 | 5.03 |
| K | 0.70 | 26.08 | 94.56 | 0.70 |
| F | 8298.50 | 0.09 | 0.25 | 0.76 |
| I | 0.73 | 30.85 | 56.92 | 91.68 |
| L | 19.74 | 0.80 | 75.51 | 0.94 |

[Amount Yield(pmol)]

| | 1 | 2 | 3 | 4 |
|---|---|---|---|---|
| D | 2.50 | 3.48 | 5.06 | 14.54 |
| E | 2.30 | 35.46 | 195.36 | 0.00 |
| N | 1.78 | 4.80 | 5.82 | 665.84 |
| S | 3.30 | 0.00 | 0.00 | 431.17 |
| T | 1.43 | 4.84 | 221.79 | 0.00 |
| Q | 0.83 | 142.42 | 0.00 | 0.00 |
| G | 10.01 | 1.07 | 358.31 | 197.96 |
| H | 0.50 | 0.18 | 86.63 | 46.41 |
| A | 2.40 | 2.94 | 107.89 | 103.95 |
| R | 0.00 | 0.00 | 0.00 | 27.71 |
| Y | 5.56 | 21.31 | 97.09 | 0.00 |
| P | 2.96 | 0.00 | 0.00 | 0.00 |
| M | 7.01 | 0.00 | 85.08 | 28.92 |
| V | 3.51 | 9.37 | 51.17 | 29.54 |
| W | 3.15 | 0.00 | 14.67 | 0.00 |
| K | 5.81 | 204.49 | 113.17 | 16.82 |
| F | 151.32 | 0.00 | 0.00 | 21.29 |
| I | 0.38 | 300.88 | 57.25 | 30.68 |
| L | 3.68 | 0.00 | 101.29 | 28.61 |

[Percent Yield]

| | | |
|---|---|---|
| Amino Acid | : | A,V,L |
| Initial Yield(%) | : | 0.00 |
| Repetitive Yield(%) | : | 0.00 |
| Correlation Coef. | : | 0.000 |
| Number of Data | : | 0 |

[Repetitive Yield(%)]

| | |
|---|---|
| Data File | : PTH-AA_08-31-2022_D01.lcd |
| Sample Name | : PTH-AA |
| Method File | : PTH-AA_08-31-2022.lcm |
| Background Data File | : |

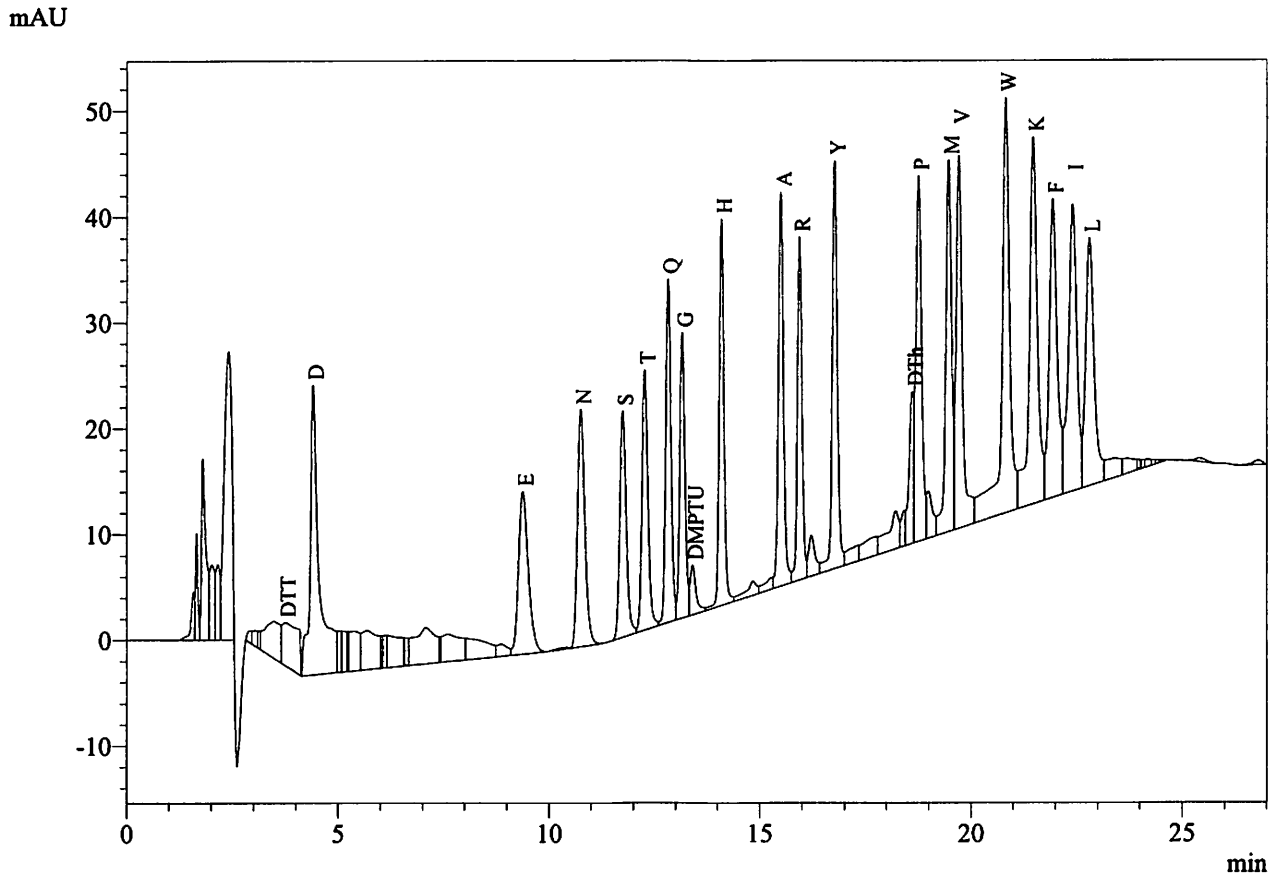


Peak Table
PDA Ch1 269nm

| Peak# | Name | Ret. Time | Area | Conc. |
|---|---|---|---|---|
| 11 | DTT | 3.743 | 113234 | 10.000 |
| 12 | D | 4.411 | 406405 | 10.000 |
| 28 | E | 9.370 | 248612 | 10.000 |
| 31 | N | 10.753 | 271970 | 10.000 |
| 32 | S | 11.744 | 226315 | 10.000 |
| 33 | T | 12.262 | 239710 | 10.000 |
| 34 | Q | 12.817 | 280338 | 10.000 |
| 35 | G | 13.148 | 228667 | 10.000 |
| 36 | DMPTU | 13.397 | 45273 | 10.000 |
| 37 | H | 14.089 | 285627 | 10.000 |
| 40 | A | 15.493 | 307920 | 10.000 |
| 41 | R | 15.935 | 244091 | 10.000 |
| 43 | Y | 16.766 | 325871 | 10.000 |
| 48 | DTh | 18.614 | 110915 | 10.000 |
| 49 | P | 18.763 | 321335 | 10.000 |
| 51 | M | 19.471 | 338785 | 10.000 |
| 52 | V | 19.711 | 362583 | 10.000 |
| 53 | W | 20.830 | 525709 | 10.000 |
| 54 | K | 21.476 | 465873 | 10.000 |
| 55 | F | 21.940 | 369418 | 10.000 |
| 56 | I | 22.410 | 380874 | 10.000 |
| 57 | L | 22.804 | 326467 | 10.000 |
| Total | | | 6425995 | |

| | |
|---|---|
| Data File | : 10048_09-02-2022_D01.lcd |
| Sample Name | : Sumit Mukherjee, Sample 1 |
| Method File | : 10048_09-02-2022.lcm |
| Background Data File | : |

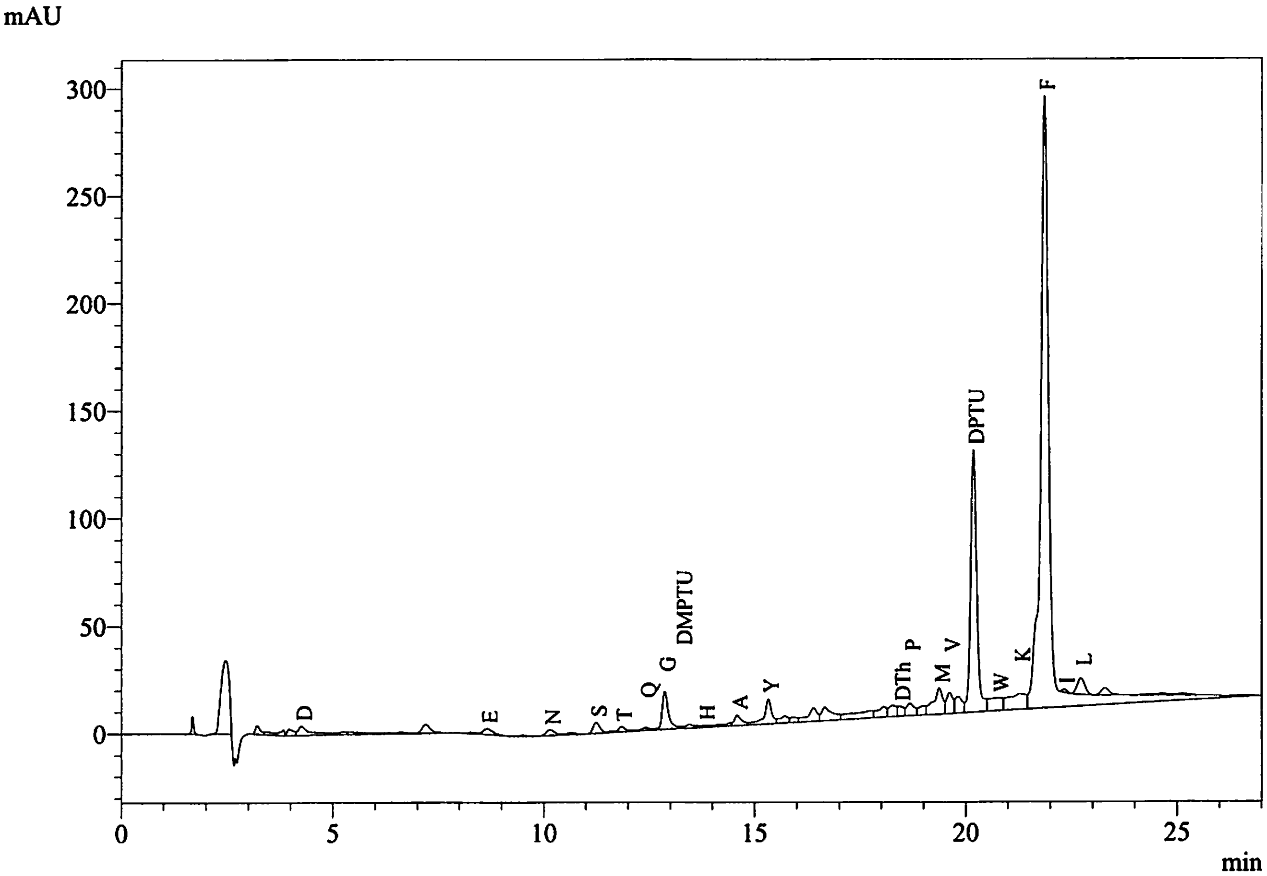


Peak Table
PDA Ch1 269nm

| Peak# | Name | Ret. Time | Area | Conc. |
|---|---|---|---|---|
| 7 | D | 4.256 | 81256 | 2.499 |
| 18 | E | 8.653 | 45759 | 2.301 |
| 20 | N | 10.139 | 38748 | 1.781 |
| 23 | S | 11.239 | 59708 | 3.298 |
| 25 | T | 11.838 | 27500 | 1.434 |
| 26 | Q | 12.412 | 18606 | 0.830 |
| 28 | G | 12.862 | 183194 | 10.014 |
| 29 | DMPTU | 13.254 | 7984 | 2.204 |
| 32 | H | 13.809 | 11406 | 0.499 |
| 35 | A | 14.583 | 59123 | 2.400 |
| 36 | Y | 15.323 | 144952 | 5.560 |
| 44 | DTh | 18.441 | 46191 | 5.206 |
| 45 | P | 18.678 | 75980 | 2.956 |
| 47 | M | 19.370 | 189979 | 7.010 |
| 48 | V | 19.614 | 101787 | 3.509 |
| 50 | DPTU | 20.177 | 1231602 | |
| 51 | W | 20.775 | 132658 | 3.154 |
| 52 | K | 21.278 | 216631 | 5.813 |
| 53 | F | 21.868 | 4471988 | 151.318 |
| 54 | I | 22.329 | 11645 | 0.382 |
| 55 | L | 22.723 | 96238 | 3.685 |
| Total | | | 7252933 | |

| | |
|---|---|
| Data File | : 10048_09-02-2022_D02.lcd |
| Sample Name | : Sumit Mukherjee, Sample 1 |
| Method File | : 10048_09-02-2022.lcm |
| Background Data File | : 10048_09-02-2022_D01.lcd |

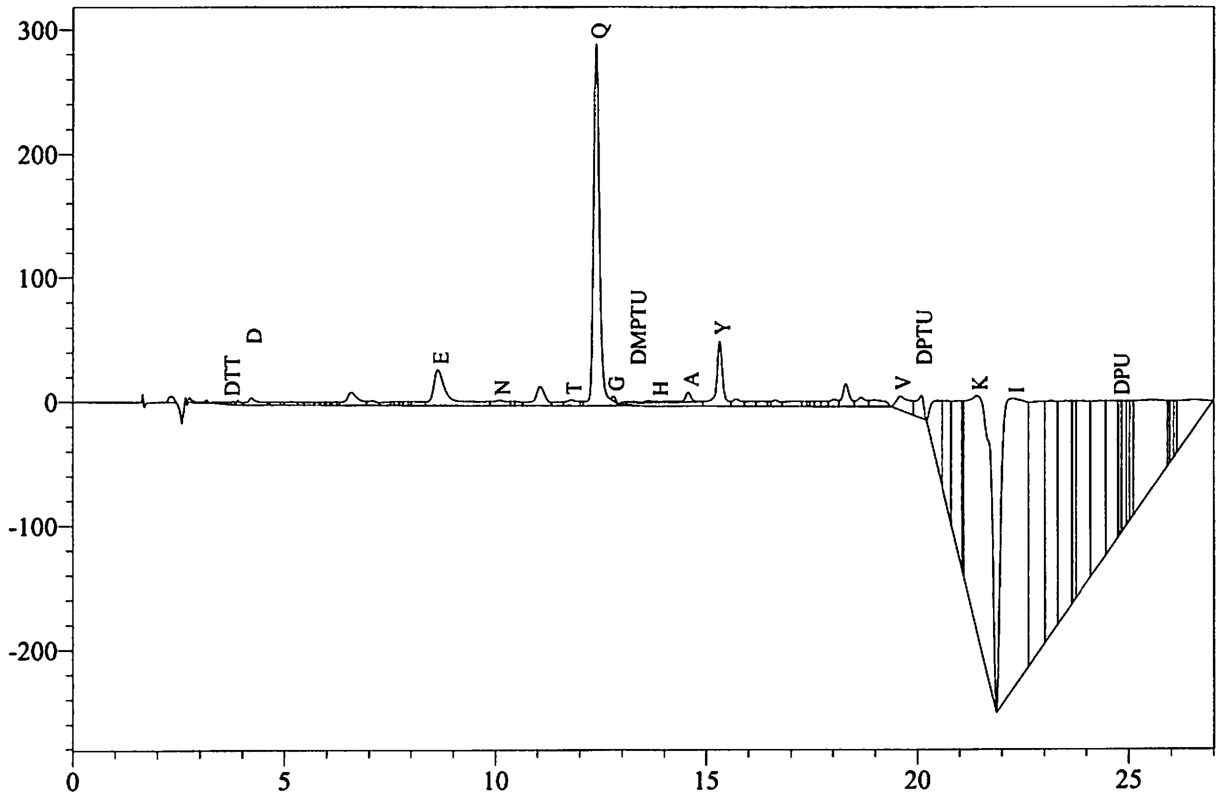


Peak Table
PDA Ch1 269nm

| Peak# | Name | Ret. Time | Area | Conc. |
|---|---|---|---|---|
| 10 | DTT | 3.688 | 15384 | 1.698 |
| 13 | D | 4.214 | 113023 | 3.476 |
| 36 | E | 8.636 | 705256 | 35.460 |
| 37 | N | 10.089 | 104526 | 4.804 |
| 42 | T | 11.794 | 92723 | 4.835 |
| 44 | Q | 12.386 | 3194107 | 142.422 |
| 45 | G | 12.795 | 19618 | 1.072 |
| 47 | DMPTU | 13.318 | 11789 | 3.255 |
| 49 | H | 13.840 | 4165 | 0.182 |
| 54 | A | 14.570 | 72479 | 2.942 |
| 55 | Y | 15.318 | 555484 | 21.308 |
| 71 | V | 19.594 | 271928 | 9.375 |
| 72 | DPTU | 20.095 | 224176 | |
| 77 | K | 21.410 | 7621402 | 204.492 |
| 78 | I | 22.271 | 9167650 | 300.875 |
| 87 | DPU | 24.768 | 314724 | |
| Total | | | 22488435 | |

Data File : 10048_09-02-2022_D03.lcd
Sample Name : Sumit Mukherjee, Sample 1
Method File : 10048_09-02-2022.lcm
Background Data File : 10048_09-02-2022_D02.lcd

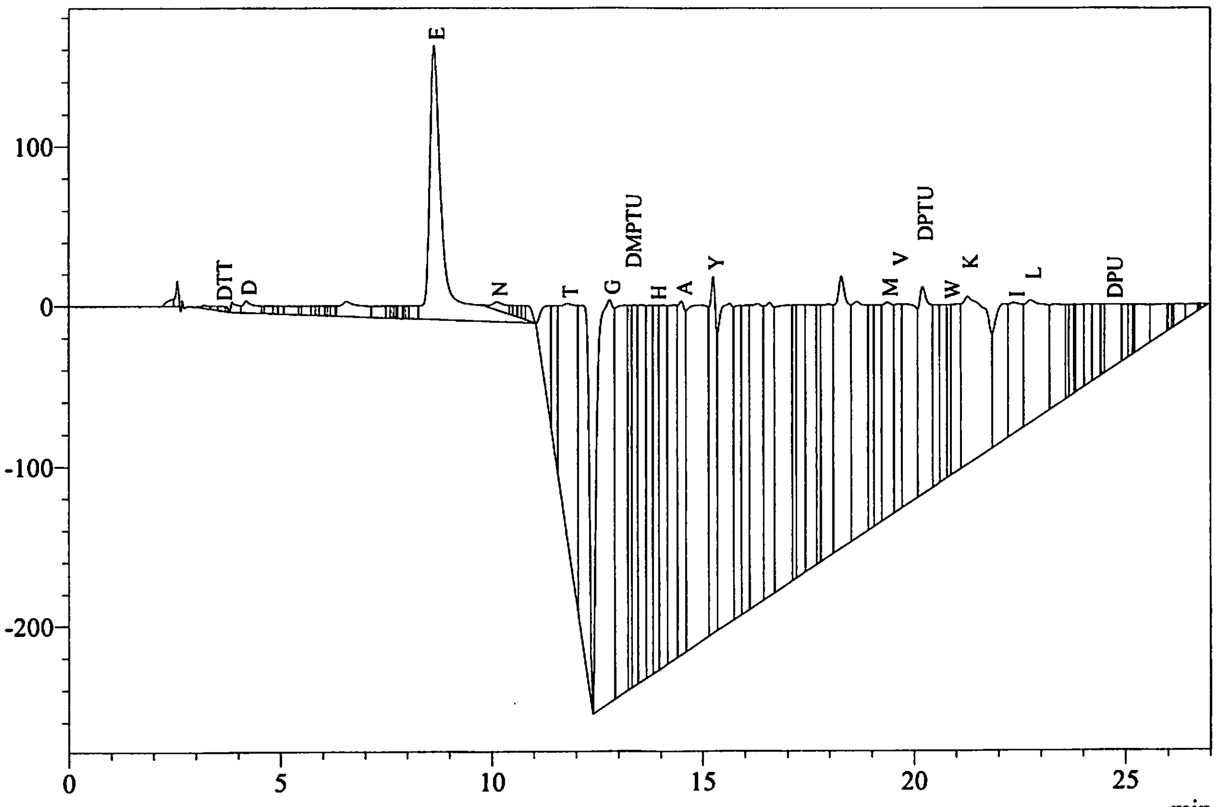


Peak Table
PDA Ch1 269nm

| Peak# | Name | Ret. Time | Area | Conc. |
|---|---|---|---|---|
| 7 | DTT | 3.612 | 32263 | 3.562 |
| 10 | D | 4.191 | 164662 | 5.065 |
| 36 | E | 8.637 | 3885593 | 195.364 |
| 39 | N | 10.119 | 126689 | 5.823 |
| 47 | T | 11.800 | 4253315 | 221.795 |
| 49 | G | 12.794 | 6554707 | 358.310 |
| 51 | DMPTU | 13.301 | 1304303 | 360.121 |
| 55 | H | 13.890 | 1979396 | 86.625 |
| 58 | A | 14.504 | 2657689 | 107.889 |
| 60 | Y | 15.256 | 2531152 | 97.092 |
| 76 | M | 19.375 | 2305937 | 85.081 |
| 77 | V | 19.633 | 1484130 | 51.165 |
| 79 | DPTU | 20.209 | 2596111 | |
| 82 | W | 20.859 | 616855 | 14.667 |
| 84 | K | 21.283 | 4217974 | 113.174 |
| 86 | I | 22.370 | 1744280 | 57.246 |
| 87 | L | 22.766 | 2645355 | 101.287 |
| 97 | DPU | 24.695 | 969397 | |
| Total | | | 40069806 | |

| | |
|---|---|
| Data File | : 10048_09-02-2022_D04.lcd |
| Sample Name | : Sumit Mukherjee, Sample 1 |
| Method File | : 10048_09-02-2022.lcm |
| Background Data File | : 10048_09-02-2022_D03.lcd |

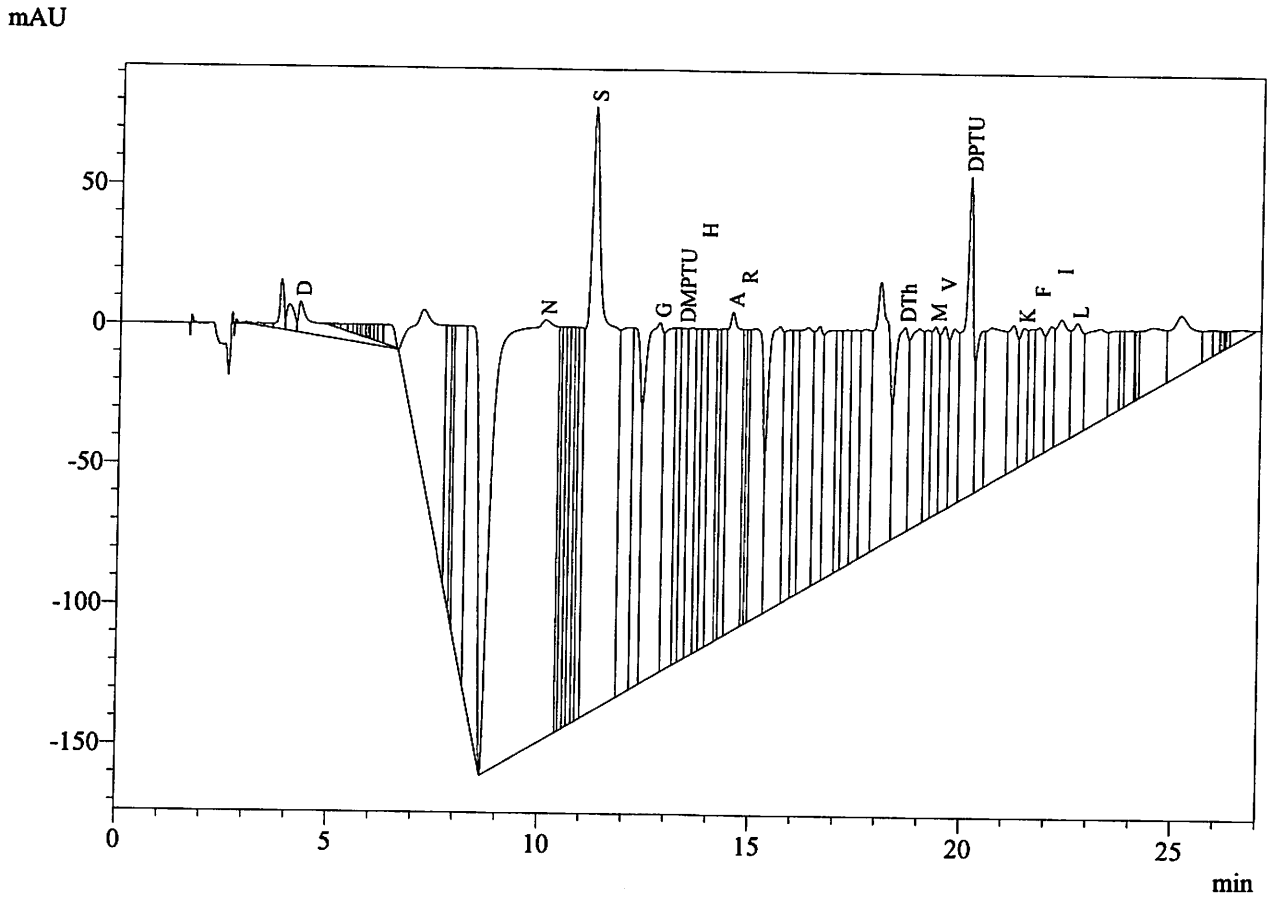


Peak Table
PDA Ch1 269nm

| Peak# | Name | Ret. Time | Area | Conc. |
|---|---|---|---|---|
| 10 | D | 4.253 | 472680 | 14.538 |
| 32 | N | 10.057 | 14487144 | 665.842 |
| 39 | S | 11.214 | 7806437 | 431.171 |
| 42 | G | 12.768 | 3621306 | 197.957 |
| 45 | DMPTU | 13.356 | 1175989 | 324.693 |
| 48 | H | 13.854 | 1060557 | 46.414 |
| 52 | A | 14.510 | 2560547 | 103.945 |
| 53 | R | 14.816 | 541185 | 27.714 |
| 67 | DTh | 18.586 | 1566119 | 176.499 |
| 70 | M | 19.305 | 783938 | 28.925 |
| 71 | V | 19.529 | 857000 | 29.545 |
| 73 | DPTU | 20.116 | 1699329 | |
| 77 | K | 21.405 | 626729 | 16.816 |
| 79 | F | 21.770 | 629053 | 21.285 |
| 81 | I | 22.287 | 934942 | 30.684 |
| 82 | L | 22.662 | 747197 | 28.609 |
| Total | | | 39570152 | |
